# Optimising network modelling methods for fMRI

**DOI:** 10.1101/741595

**Authors:** Usama Pervaiz, Diego Vidaurre, Mark W. Woolrich, Stephen M. Smith

**Affiliations:** Oxford Centre for Functional MRI of the Brain (FMRIB), Wellcome Centre for Integrative Neuroimaging, Nuffield Department of Clinical Neurosciences, University of Oxford, Oxford OX3 9DU, United Kingdom; Oxford Centre for Human Brain Activity (OHBA), Wellcome Centre for Integrative Neuroimaging, Department of Psychiatry, University of Oxford, Oxford, OX3 7JX, United Kingdom

**Keywords:** Functional Connectivity, Connectome, Netmat, Riemannian geometry, Deep Learning, Convolutional Neural Networks

## Abstract

A major goal of neuroimaging studies is to develop predictive models to analyse the relationship between whole brain functional connectivity patterns and behavioural traits. However, there is no single widely-accepted standard pipeline for analyzing functional connectivity. The common procedure for designing functional connectivity based predictive models entails three main steps: parcellating the brain, estimating the interaction between defined parcels, and lastly, using these integrated associations between brain parcels as features fed to a classifier for predicting non-imaging variables e.g., behavioural traits, demographics, emotional measures, etc. There are also additional considerations when using correlation-based measures of functional connectivity, resulting in three supplementary steps: utilising Riemannian geometry tangent space parameterization to preserve the geometry of functional connectivity; penalizing the connectivity estimates with shrinkage approaches to handle challenges related to short time-series (and noisy) data; and removing confounding variables from brain-behaviour data. These six steps are contingent on each-other, and to optimise a general framework one should ideally examine these various methods simultaneously. In this paper, we investigated strengths and short-comings, both independently and jointly, of the following measures: parcellation techniques of four kinds (categorized further depending upon number of parcels), five measures of functional connectivity, the decision of staying in the ambient space of connectivity matrices or in tangent space, the choice of applying shrinkage estimators, six alternative techniques for handling confounds and finally four novel classifiers/predictors. For performance evaluation, we have selected two of the largest datasets, UK Biobank and the Human Connectome Project resting state fMRI data, and have run more than 9000 different pipeline variants on a total of ∼14000 individuals to determine the optimum pipeline. For independent performance validation, we have run some best-performing pipeline variants on ABIDE and ACPI datasets (∼1000 subjects) to evaluate the generalisability of proposed network modelling methods.

## 1. Introduction

It is valuable to understand large-scale networks of brain activity. Most of the literature is focused on recognizing brain networks that are spatially and temporally linked to a given task [1]. In this paper, we aim to explore brain networks measured from resting state functional magnetic resonance imaging (rfMRI). In contrast to task-based fMRI, during rfMRI the brain is not occupied by a pre-defined task. However, studies have shown that the brain activity at rest is not random [2, 3, 4], but is hierarchically organized in time and significantly correlated with behaviour [5].

The analysis of rfMRI data largely relies on estimates of functional connectivity. Functional connectivity is defined as the statistical association between time-series of anatomically distinct brain regions [6], which in fMRI is typically calculated as zero-lag correlation. To simplify, if two brain regions have blood-oxygen-level-dependent (BOLD) signals that are correlated, it means they are functionally connected. Functional connectivity is dynamic as it can be observed to change over time [5, 7, 8]. Contrasting with this metric is static connectivity, which is an average of all dynamic connectivity states, and is thereby is less noisy. Static (or average) connectivity is also simpler to estimate and interpret, and is at this point in more common usage. The methodological research conducted in this article is purely based on exploring static functional connectivity.

A number of research studies have shown the utility of functional connectivity in identifying neuropsychiatric disorders such as Alzheimer’s disease [9], depression [10], social anxiety disorder [11] autism spectrum disorder [12], Parkinson’s disease [13] and schizophrenia [14]. Resting state functional connectivity also has been shown to correlate with emotional measures [15] and behavioral performance [16, 17]. Overall, functional connectivity is gaining visibility as a salient tool for assessing functional brain organization and as an important biomarker for neurological disorders.

As the rfMRI scan is composed of thousands of voxels at each time-point per subject, this poses extreme challenges to working with raw voxel-wise time-series data. To avoid working in high dimensional space (and to increase interpretability and effective SNR), the standard processing procedure entails the determination of parcels which are defined over a group of voxels sharing similar timecourses. However, the method by which the brain is segmented into these different regions, or parcels, before estimating functional connectivity is far from standardized [19]. Some of the existing parcellation techniques are based on mapping anatomical or functional atlases onto an individual’s brain [20, 21, 22]. Other brain parcellation techniques are more data driven and attempt to derive parcels based upon common features within the data [23, 24, 25]. After parcellating the brain, the next step is estimating functional connectivity. Functional connectivity is conventionally estimated through taking a full (Pearson) correlation between brain parcels over time [18]. The use of full correlation in deriving these estimates however, has some drawbacks as this method does not distinguish direct vs. indirect pathways by which two brain regions may be connected [26]. Additionally, the utilization of this method results in numerous other limitations related to low SNR, primarily because of short scanning sessions. This paper provides a detailed account of several functional connectivity estimation techniques that may mitigate these challenges, which consequently can result in enhanced sensitivity in signal detection.

After derivation of brain parcels and functional connectivity estimates, there remains further decisions of how to optimally utilize these in conjunction with other data, such as behavioral metrics. The commonly followed procedure is to directly use elements of correlation-based functional connectivity estimates as features for regression or classification algorithms. This direct usage however has fallen subject to criticism, as correlation-based functional connectivity does not naturally form a Euclidean space [27, 28]. In this paper, we evaluate an arguably more principled approach to represent correlation-based functional connectivity using Riemannian geometry [27, 28]. This method is founded upon mathematical principles which involve manipulating correlation-based estimates in the tangential space of Riemannian manifold [30] where Euclidean geometry can be applied.

Additionally, there is also a concern of noise causing an increase in inter- and intra-subject variance in functional connectivity estimates. One method to alleviate this is to apply conventional regularisation techniques in the ambient space when functional connectivity is estimated. The other approach is to apply covariance shrinkage estimators [31] in the tangential space of Riemannian manifold. In this paper, we have evaluated the effects of covariance shrinkage techniques both in the ambient space and the tangent space (isotropic [32] and non-isotropic covariance shrinkage techniques [33]) on the prediction of individual demographic and behavioural phenotypes. Moreover, we have assessed the performance of shrinkage approaches in following framework, (i) no shrinkage in the ambient space or the tangent space, (ii) shrinkage in the ambient space but no shrinkage in the tangent space, (iii) no shrinkage in the ambient space but shrinkage in the tangent space, and (iv) shrinkage approaches both in the ambient space and the tangent space.

To predict individual phenotypes from rfMRI data, the elements of the functional connectivity matrix, where each element represents the strength of connection between two parcels, can be used as input features for a machine learning classifier (or continuous variable predictor). In this paper, we compare the performance of a selection of existing state of the art methods with several proposed deep learning based architectures. We also demonstrate a baseline application for using neural networks in brain functional connectivity modelling.

There is currently no general agreement on a single common pipeline for estimating and using functional connectivity. We aim to help with this situation by answering several major questions:

1. How should we parcellate the brain? Should this technique be data driven or based upon hard parcellation? How many parcels should we derive?
2. How should we estimate correlation-based functional connectivity? Should we remain with full, or partial correlation, or do we have better choices?
3. Should we preserve the geometry and shape of correlation-based functional connectivity elements within a Riemannian framework, and what would be the optimal way to do this?
4. Should we utilize covariance shrinkage estimators? Should shrinkage occur in ambient space of covariances or in tangent space?
5. How should we remove the effect of confounding variables in rfMRI?
6. Which classifier/predictor provides optimal prediction of age, sex and other non-imaging variables using these functional connectivity estimates? Is a deep learning approach of value?

The questions raised above may be treated as six steps to derive a framework for estimation and accurate usage of functional connectivity estimates. This paper will attempt to address the above questions in order to propose a comprehensive pipeline that takes into account several central methodological issues when estimating and using functional connectivity.

For performance evaluation of the different methods, we have chosen two rich datasets which contain both rfMRI and phenotypic data. The first is 13301 subjects from UK Biobank (UKB) [36] and other is 1003 healthy subjects from the Human Connectome project (HCP) [37]. Our assessment criteria are based on the prediction accuracy of age, sex, fluid intelligence and neuroticism score, using correlation-based functional connectivity estimates. To establish the most optimum and reliable pipeline, we have run more than 9000 combinations of different steps of methodology detailed below. This computation has involved thousands of hours of computing on both CPUs and GPUs. The overall methodology is summarized in Figure 1.

**Figure 1:**
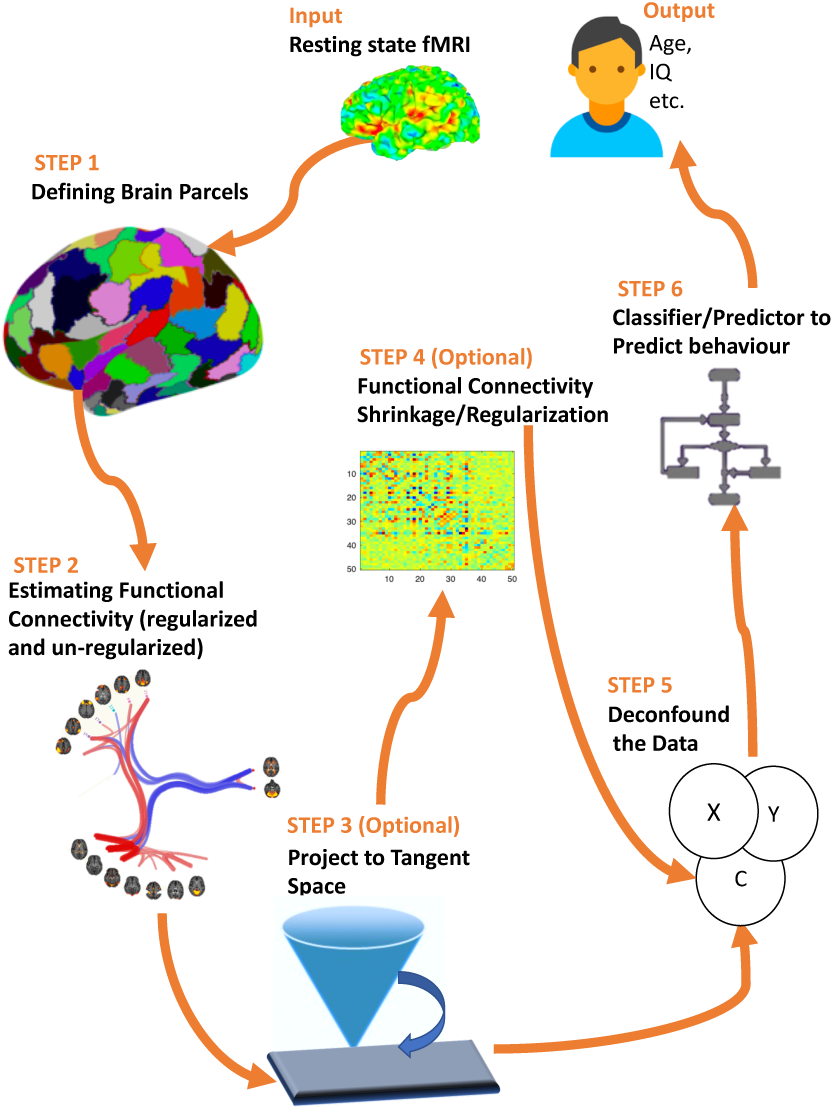
Flow chart summarizing the six major steps of the methodology framework for rfMRI analysis.

## 2. Methods

### 2.1. Finding the best parcellation approach

The standard procedure for parcellation entails the grouping of voxels sharing similar time-courses into parcels. A parcel can be any spatial region of the brain, associated with a single time course (found by averaging the spontaneous fluctuation over all voxels within the parcel). For our analyses, we considered the following spatial properties in parcels, 1) parcels may or may not be allowed to overlap, 2) parcels can have both positive and negative spatial weight, or be binary masks 3) parcels may be composed from multiple disconnected regions.

Following the above criteria, we estimated parcels (spatial maps and associated time-series) across the brain using four different methods. In the literature, parcels can also be referred as nodes, modes, networks or region of interests, but here for simplicity and continuity, we will only refer to them as parcels. We only consider rfMRI-driven parcellations, as there is plenty of existing evidence that these outperform parcellations derived from pre-defined structural atlases [38, 39, 40].

#### 2.1.1. Data-driven Parcellation: SHEN functional Parcels

The simplest way to identify parcels is to extract time courses from pre-defined, labelled regions, which often come from anatomical atlases. For resting state studies, a preferred method is to use a functional brain atlas based on correlated BOLD signal, rather than anatomical distinctions [39]. In this study, we applied a functional brain atlas known as the SHEN parcellation [20] which covers the cortex, subcortex and cerebellum. To derive the SHEN atlas, a spectral clustering method was applied which ensured the functional homogeneity within each parcel [20]. SHEN parcellation is based on rfMRI data of 79 healthy normal volunteers and provides more coherent resting state time-course estimates when compared with anatomical atlases. The total number of parcels in SHEN parcellation is 268. SHEN is a volumetric parcellation, and for both UKB and HCP data, we used the preprocessed released data in standard volumetric space after ICA FIX denoising [41].

#### 2.1.2. Data-driven Parcellation: YEO functional Parcels

We applied another functional brain atlas known as the YEO parcellation [42]. The YEO parcellation is based on rfMRI data from 1489 subjects which were registered using surface-based alignment. To derive the YEO atlas, a gradient weighted Markov random field approach was employed which integrates local gradient and global similarity approaches. This ensured the parcels were functionally and connectionally homogeneous, and also agreed with the boundaries of certain cortical areas defined using histology and visuotopic fMRI. The YEO parcellation is available at multiple resolution levels, and we have applied the YEO parcellation where the total number of parcels is 100. The YEO parcellation is available both in volumetric and grayordinate space. For HCP data, we use the minimally-preprocessed released data in grayordinates, and for UKB data, we used the preprocessed released data in standard volumetric space. UKB has not yet been processed with a pipeline like HCP, but future UKB data releases will include versions of the data transformed into grayordinate space. For now, this helps to span the space of approaches commonly taken (some volumetric, some surface-based). YEO parcellation is often known as the “Schaefer” parcellation, from the name of the first author [42].

#### 2.1.3. Data Driven Soft Parcellation: Spatially Independent Parcels

We also applied a popular data-driven parcellation scheme called Group Independent Component Analysis, or GroupICA [34, 35] to our data. There are two popular variants of GroupICA, spatial ICA (sICA), which identifies spatially independent components, and temporal ICA (tICA) [43], which put restrictions on temporal dynamics. The parcels identified by tICA are coerced to have orthogonal time-series which therefore precludes further network analyses of the kind covered in this paper. Therefore, we opted for sICA and applied ^2^ sICA by concatenating the time courses from all subjects. The ICA parcels can be spatially overlapping and non-contiguous. The dimensionality (d) of sICA corresponds to the number of desired parcels. For HCP data, we applied sICA at d = 15, 50 and 200, and for UKB, d = 25 and 100. With UKB data, we disregarded four components from ICA 25, and 45 components from ICA 100 as they were judged to be artefacts ^3^. This left us with d = 21, and 55 for UKB^4^. For HCP data we use the minimally-preprocessed released data, in grayordinates (cortical surface vertices and subcortical voxels), with ICA FIX denoising having been applied [41]. For UKB data, we used the preprocessed released data in standard volumetric space after ICA FIX denoising.

#### 2.1.4. Data Driven Parcellation: Probabilistic Functional Parcels

While ICA based approaches provide a choice between spatial or temporal independence, these approaches can be problematic as the brain is probably not perfectly segregated either in space or time, and they typically to do not have any modelling of between-subject variability. We applied another framework used for identifying large-scale probabilistic functional modes (PROFUMO) [45, 46] to identify parcels which are allowed to be correlated with each other in space and/or time, and which explicitly models between-subject variability in the parcels.. In contrast to the above methods, therefore, PROFUMO does not require parcels to be orthogonal or non-overlapping. We applied PROFUMO on the HCP for d = 50, but we are not yet able to apply PROFUMO on the UKB data due to prohibitive computational cost with the larger cohort.

### 2.2. Finding the best correlation-based functional connectivity estimate method

We extracted a timeseries signal for each subject, *X* ∈ (ℝ)^*T ×d*^, where *T* corresponds to the number of timepoints and *d* correponds to the number of parcels for each subject. Functional connectivity estimates are also referred to in the literature as parcellated connectomes or netmats, but here we will refer to these estimates as functional connectivity. Table 1 lists symbols for the different correlation-based functional connectivity estimates we evaluated.

**Table 1:** The symbols used for various functional connectivity estimates in this paper.

| Term | Symbol |
| --- | --- |
| Empirical covariance | $\check{\Sigma}$ |
| Full correlation | $\check{\Sigma}$ |
| Precision matrix | $\hat{\Omega}$ |
| Tikhonov precision matrix | $\bar{\Omega}$ |
| Oracle approximate shrinkage precision matrix | $\dot{\Omega}$ |
| Partial correlation | $\rho$ |
| Any of above functional connectivity matrix | $C$ |
| Any of above functional connectivity matrix in tangent space | $\vec{C}$ |

#### 2.2.1. Estimating the Functional Connectivity

##### Empirical Covariance

Functional connectivity is calculated by estimating the empirical (sample) covariance matrix of timeseries signal for each subject. The range of elements of the empirical covariance matrix, 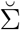, lies between −∞ and +∞ and can be defined as:

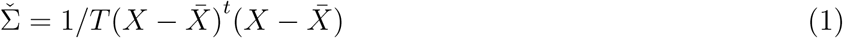

##### Full Correlation

The full correlation coefficient, or Pearson’s correlation coefficient can be defined in terms of covariance. Mathematically, full correlation 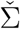 is a standardized form of covariance. The value of correlation takes place between -1 and +1. Conversely, the value of covariance lies between −∞ and +∞.

##### Partial Correlation

Although commonly used, using full correlation to derive connectivity estimates is problematic, as it does not distinguish whether two parcels of brain are directly connected or indirectly connected through another parcel. To mitigate this, we define partial correlation, which is correlation between the time series of two parcels after adjusting for the time series of all other network parcels. Partial correlation is calculated from the inverse of the covariance matrix, also known as the precision matrix. We calculated inverse covariance using cholesky decomposition [47] as illustrated in Equation 2. For this derivation, we first factorized 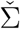 into an upper triangular *U* that satisfies 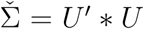 and then calculated the precision matrix 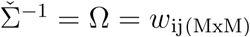. To convert precision into partial correlation *ρ*, there is a sign flip of the non-diagonal elements and also normalization as shown in Equation 3 where i and j are nodes.

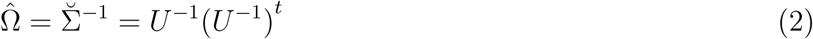

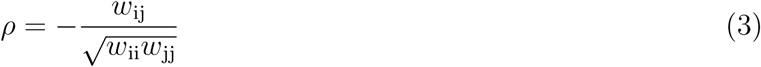

##### Tikhonov Partial Correlation

As partial correlation involves the calculation of the inverse of covariance matrix, this method becomes problematic when there are not considerably more timepoints than nodes. Several approaches based on regularization have been proposed in the literature to address this unstable matrix inversion problem. We implemented the Tikhonov regularization (also referred as L_2_ ridge regression) [48] as shown in Equation 4. This equation involves the addition of a regularization term Γ = *αI*, a multiple of the identity matrix. The scalar *α* is optimized here by minimizing the root mean square distance ^5^ between regularised subject precision matrices 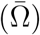 and the group average of the unregularised subject precision matrices 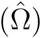. We optimised a single *α* for all the subjects.

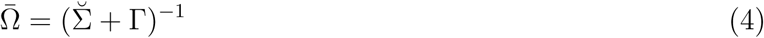

##### Sparse Group Gaussian Graphical Modelling (SGGGM) Partial Correlation

Tikhnov regularization allows for a stable inversion for the covariance matrix; however, it uniformly shrinks off-diagonal elements. Sparse Group Graphical Gaussian Modelling (SGGGM) [49] learns group-level connectivity in a data-driven way using an L_1_ prior and regularization of each subject’s connectivity towards the group mean using an L_2_ prior. The advantage of SGGGM is that it supports intra-subject inverse covariance being similar, but not exactly the same as the group inverse covariance estimate in a similar manner to [50]. The inverse covariance 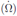 fed into SGGGM, is calculated using the regularization technique known as oracle approximating shrinkage (OAS), as described in the original paper [49], and is well regularized. SGGGM is posed as a regularized consensus optimization problem which can be solved using an alternating direction method of multipliers (ADMM) [51]. In Equation 5, ‖ ‖. corresponds to the Frobenius norm penalty, applied to the difference between the subjects’ matrices and the group. 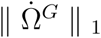 is used for enforcing sparsity on the group prior and 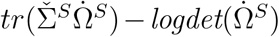 is the representation of each subject’s own observation. Parameter *λ*_1_ controls sparsity and *λ*_2_ regulates the group consistency. We optimized these hyper-parameters (*λ*_1_ and *λ*_2_) by minimizing the root mean square distance between regularised subject precision matrices estimated using SGGGM and the group average of the unregularised subject precision matrices 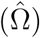.

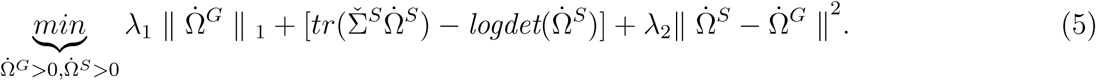

#### 2.2.2. Tangent Space Parameterization

Covariance matrices ^6^by construction are symmetric and live in space of positive definitive (PD) matrices. The PD matrix is a symmetric matrix whose eigenvalues are all positive. The standard practice is to directly vectorize (unwrap) these covariance matrices and then feed them into a machine learning predictor/classifier, however there are a few downfalls to this practice. One issue is that PD matrices do not naturally form a Euclidean space, meaning mathematical operations like subtraction do not apply, if geometry is to be preserved (and the PD nature preserved). For example, the difference of two PD matrices does not correspond to the PD covariance matrix of a signal. *C*_1_, *C*_2_ ∈ *Sym*^+^_n_ ⇏ *C*_1_ − *C*_2_ ∈ *Sym*^+^_n_ [27]. Another problem is that in the space of the positive definitive cone, where covariance matrices exist, elements of covariance are inter-correlated. This goes against the underlying assumption for some predictors/classifiers, which assume features are uncorrelated. Lastly, these vectorized connectivity estimates do not follow a normal distribution, which violates the assumptions of some methods, like Linear Discriminant Analysis (LDA).

If we follow the geometry of PD matrices, we note that the space of PD matrices forms a differentiable manifold *M*. PD matrices therefore cannot directly be treated as an element of Euclidean space; therefore, to correctly apply mathematical formulations on these matrices, we utilise Riemannian geometry [29]. For each covariance matrix *C* in manifold *M*, we can define a scalar product in the associated tangent space (T) as *MT*_*C*_. This tangent space is homomorphic to the manifold, and therefore we can approximate the Riemannian distance in the manifold by using Euclidean distance in the tangent space. This allows us to apply simple Euclidean mathematical operations in the tangent space without breaking the geometry of connectivity matrices. Moreover, many popular classification/prediction algorithms (for example, Neural Network, LDA and SVM) cannot be directly implemented to act in a Riemannian manifold. As the projected connectivity matrices in the tangent space are treated as Euclidean objects, such classification algorithms can be readily applied in the associated tangent space.

Another important consideration for the tangent space parametrization is that the projection requires a reference point in the manifold (this is the point where the tangent plane touches the manifold) that should be close to the subject’s projected covariance matrix. If we have a different reference point for each subject, each subject’s covariance matrix would be projected to a different tangent plane. To ensure that all projected covariance matrices lie in the same tangent plane, we must find a reference covariance matrix close to all subjects’ covariance estimates. This reference covariance matrix could be an average of the whole set of covariance matrices [53] and is referred to here as *C*_*G*_. Following the logarithmic map for projecting any covariance matrix to its tangent plane [54], any covariance matrix *C* can be projected to tangent space as shown in Equation 6, where *log*_*m*_ is matrix logarithm. Once the covariance matrices are in tangent space, they are no longer linked by the PD constraint, and hence these uncorrelated features are more useful for classifiers/predictors.

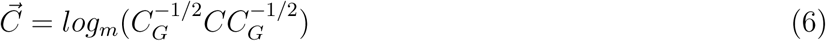

##### Reference Point in Tangent Space

*C*_*G*_ symbolizes the group reference estimate which may be estimated in different ways. The choice of reference PD matrix would normally be a group mean estimate as stated in Table 2. Euclidean mean is the average of all covariance matrices whereas harmonic mean is the reciprocal of the arithmetic mean of the reciprocals. The Euclidean and harmonic means are calculated in the manifold space. Log Euclidean is calculated by first applying matrix logarithm, then computing the mean, and lastly applying the matrix exponential to bring back the mean to the manifold. The Riemannian mean is introduced in [55] and can be computed in three iterative steps: (1) projecting connectivity estimates to the tangent space (initialized by using the arithmetic mean, *C*_*e*_), (2) calculating the arithmetic mean in the tangent space, 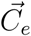, and (3) mapping the arithmetic mean back to the manifold i.e., 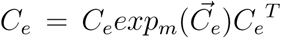. These three steps are repeated until convergence 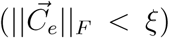 [28]. The Kullback mean is the geometric mean between harmonic mean and arithmetic mean. It is not evident in the literature which mean would lead to better representation of the manifold and therefore, we considered all these alternatives for reference point estimation.

**Table 2:** Reference Mean Estimation. *C*_*i*_ denotes the functional connectivity matrix for subject i.

| Reference Mean | Equation |
| --- | --- |
| Euclidean | $C_e = \frac{1}{N} \sum_i C_i$ |
| Harmonic | $C_h = (\frac{1}{N} \sum_i C_i^{-1})^{-1}$ |
| Log Euclidean | $C_{le} = \exp_m (\frac{1}{N} \sum_i \log_m C_i)$ |
| Riemannian Mean | $C_r = \arg \min (\sum_i \delta_R(C_e C_i)^2)$ |
| Kullback | $C_k = C_e^{1/2} \left( C_e^{-1/2} C_h C_e^{-1/2} \right)^\alpha C_e^{1/2}$ |

#### 2.2.3. Shrinkage in Tangent Space

To further reduce estimation variability, we can apply regularization in the tangent space. The Ledoit-Wolf estimator shrinks the covariance towards a target matrix T as 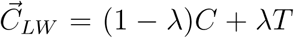. In [32], the authors proposed shrinkage towards an identity matrix. A better alternative is to shrink towards the group mean covariance matrix. The Ledoit-Wolf shrinkage can also be applied in tangent space as shown in Equation 7.

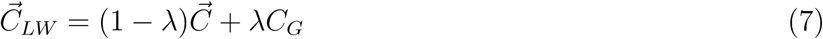

This shrinkage is isotropic and therefore does not take into consideration the distribution of the population. Recently, another method called PoSCE was proposed [33] in which the population prior is not based on just the mean, but uses information from the probabilistic distribution of covariances. The prior dispersion is calculated as the mean outer product of the parametrized tangent space connectivity estimates as 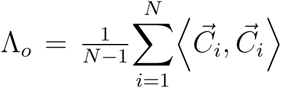 and Λ = *λI* is likelihood covariance where *λ* is the shrinkage control parameter. The shrunk covariance matrices in the tangent space can then be calculated as in Equation 8.

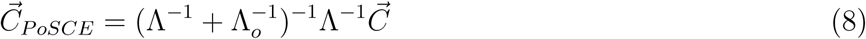

These isotropic and non-isotropic shrinkage techniques could ideally be applied to any projected connectivity estimate (covariance, full correlation, partial correlation, Tikhonov partial correlation or SGGGM partial correlation) in the tangent space. Theoretically, Tikhonov partial correlation and SGGGM connectivity estimates are already well-regularized so they may not require any further shrinkage in the tangent space, but we have tested all possible variants in this study (regularization in the ambient space and then further regularization in the tangent space). Moreover, we have introduced concepts of three spaces as detailed below:

- **Ambient Space**: The estimated functional connectivity estimates (Section 2.2) were not projected into the tangent space. The classifier/predictor was directly applied to connectivity estimates in the ambient space.
- **Tangent Space**: The various functional connectivity estimates (covariance, full correlation, partial correlation, Tikhonov partial correlation or SGGGM partial correlation as defined inSection 2.2) were projected to tangent space following Section 2.2.2, but no shrinkage was applied in the tangent space. The classifier/predictor was applied on the parameterized tangent space connectivity estimates.
- **Tangent Shrinkage Space**: The shrunk tangent space in which functional connectivity estimates were projected to tangent space and then shrinkage was applied in the tangent space (Section 2.2.3). In this case, the classifier/predictor was applied on the parameterized shrunk tangent space connectivity estimates. Most of the analyses carried out in this article are based on using PoSCE tangential space shrinkage, however Figure A.19 depicts results results comparing PoSCE shrinkage with Lediot-Wolf shrinkage in the tangent space.

### 2.3. Finding the best classifier/predictor

The last step is to fed these covariance matrices to predictor/classifier^7^ as features to predict non-imaging variables.

#### Elastic Net

The first classifier/predictor we tested is Elastic Net [56], which is a regularized regression method that combines the penalties of lasso (*L*_1_) and ridge (*L*_2_) methods. *L*_1_ encourages a sparser model (more zeros) but fails to accomplish group variable selection (where strongly correlated features are selected or discarded together) [57]. Alternatively, *L*_2_ encourages the grouping effect and removes the limitation on the number of chosen variables but it does not achieve sparsity. *L*_2_ is a reasonable approach to handle the ill-posed problems that result from extremely correlated features. Elastic net aims to overcome the limitations of both *L*_1_ and *L*_2_ penalties. The data matrix X is *s*×*p*, where *s* is the number of subjects and p is the number of features, and *y* is the response vector, such as age or sex. The functional connectivity matrices are symmetric and only values above the diagonal need to be retained and vectorized, and hence *p* = *d*(*d*-1)/2, where *d* is number of parcels. We then applied univariate pre-feature selection, and only retained the top 50% features (based on correlation with the target variable). In Equation 9, ‖_1_ corresponds to the *L*_1_ norm penalty and the quadratic part (the second term) is the is *L*_2_ norm.

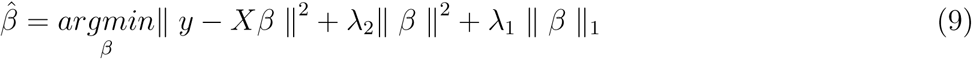

#### BrainNetCNN

The second classifier/predictor, BrainNetCNN, is specifically developed to leverage the topological structure of networks [60]. It consists of edge-to-edge (E2E), edge-to-node (E2N) and node-to-graph (N2G) convolutional filters, which are all designed to analyze functional connectivity to predict the behavioral information. The main contribution of BrainNetCNN is using a cross shape filter (the dimension of some of the filters are set to be *d* × 1 and others 1 × *d*, whereas, the overall size of input connectivity matrix is *d* × *d*) instead of the traditional box shape filter. There are primarily three main layers used in this network, E2E, E2N and N2G layers. The E2E layer considers the weights of all the edges which share a node together. Intuitively, the E2E layer resembles a convolutional filter, as for an edge (j,k) in a functional connectivity matrix, it combines the signal with the signal from direct neighbours (edges connected to either node j or node k), but applies a cross shape filter instead of a box shape filter. The output of the E2E layer is fed into the E2N layer. The E2N layer is equivalent to convolving the functional connectivity matrix with a spatial convolutional 1D row filter and a 1D column filter (and adding their results). It gives a unique output value for each node, j, as it takes the average of incoming and outgoing weights of each edge associated to node j. Finally, the N2G graph layer is used, which is similar to the fully connected layer. The N2G layer reduces the spatial dimensionality, and outputs a single scalar for weighted combination of nodes per feature map. The output features from the N2G layer are then fed into a predictor to provide final prediction (non-imaging variables). The architecture of the BrainNetCNN is shown in Figure A.13 (sub-figure A).

#### Recurrent Convolutional Neural Network

A Convolutional neural network (CNN) is a feed-forward network inspired by the brain’s visual system; however, in comparing it with the brain, CNN lacks the recurrent connections which are abundant in the brain. Inspired by [58], we have designed a recurrent convolutional neural network (RCNN) by incorporating recurrent connections in each convolutional layer. The activities of RCNN evolve over time (the activity of each unit is modulated by the activities of its neighboring units), which provide the advantage of an increased receptive field, and results in a greater awareness of contextual information. We have implemented a 2D RCNN to extract features from functional connectivity matrices. The most important module of RCNN is the recurrent convolution layer (RCL). The net input *z*_*ijk*_ at unit (i,j) on kth feature map is given by

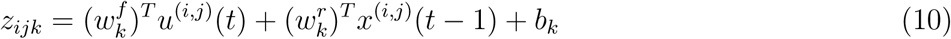

In Equation 10, *u*^(*i,j*)^(*t*) is the feed forward input, as in a CNN, while 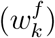 is the vectorized feed forward weights. The second term of the equation arises because of the recurrent connections and *x*^(*i,j*)^(*t* − 1) represents the recurrent input and 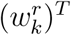 is associated with recurrent weights. In other words, *u*^(*i,j*)^(*t*) and *x*^(*i,j*)^(*t* − 1) are centered at (i, j), and are the vectorized square patches of the feature maps in the present and foregoing layer respectively. The last term of the equation, (*b*_*k*_), is the bias. *z*_*ijk*_ will pass through the rectified linear activation function *f* (*z*_*ijk*_) and then local response normalization (LRN) will be applied on it, which imitates lateral inhibition in cortex. In neurobiology, lateral inhibition refers to the capacity of an excited neuron to reduce the activity of its neighbors. Similarly here, feature maps will challenge each other for maximum representation. The architecture of 2D RCNN is shown in Figure A.12. Equation 10 describes the behaviour of one RCL (recurrent block) in Figure A.12. Unfolding the RCL for t = 0 will result in a purely feed-forward network. In Figure A.12, we have unfolded each RCL for t = 3, leading to a feed-forward network with largest depth of t+1 = 4 and smallest depth of t = 1.Each RCL block has convolution, ReLU, batch normalization and addition layers (collectively named as a residual block). Moreover, each residual block receives input directly from feed-forward and recurrent connections. The recurrent input evolves over iterations but the feed-forward input remains the same for all iterations. The overall 2D RCNN is composed of four RCL blocks, and between each RCL block, only feed forward connections are used. The output from the last RCL is fed into fully connected layers.

#### Graph Convolutional Neural Network

The standard convolution is limited to data on a Euclidean grid; however, the graph convolutional neural network (GraphCNN) extends beyond traditional CNNs to handle data that is supported on a graph. GraphCNN exploits the Laplacian of the graph and is built on spectral filters, graph coarsening, and efficient pooling, which are all based on established tools in graph signal processing [59]. We followed the approach of [61] and constructed the graph *G* = (*V, E, A*) where *d*_*x*_ = *V* is the set of vertices, *E* is the edges and *A* is the adjacency matrix. GraphCNN takes input as a feature matrix 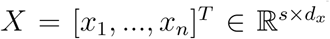, where *s* is number of nodes and *d* is number of features for each node. In our case, each subject is represented by a node and corresponding features are computed by vectorizing the functional connectivity matrix. The other input to the GraphCNN is an adjacency matrix, 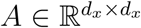 which is essentially a graph between features. We defined the adjacency matrix based on the similarity estimate between features. For constructing the graph, the Euclidean distance *d*_*ij*_ = ‖*x*_*i*_−*x*_*j*_‖_2_ was used as a distance function, a Gaussian kernel 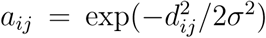 was applied and a *k* nearest neighbours graph was formulated. The architecture of GraphCNN is shown in Figure A.13 (sub-figure B).

##### 2.3.1. Deconfounding the Data

Many participant-related confounds exist in resting state data, which can corrupt associations between the imaging data and non-imaging data. Our main goal was to predict non-imaging variables based only on *pure* functional connectivity, hence we found it was crucial to minimize the impact of confounding variables. The basic confounds we controlled for were age, sex, ethnicity, weight, average head motion, brain size, intercranial volume, variables (x, y, z, table) related to bed position in scanner, imaging centre and confounds modelling slow date-related drift. We also included some non-linear effects, i.e., *age*^2^, *sex* × *age, sex* × *age*^2^. We evaluated the effect of various deconfounding schemes on X, a data matrix (i.e, subjects’ functional connectivity), and Y, a response vector (e.g, fluid intelligence). For X there are three possible strategies, (i) no deconfounding (X0), (ii) confound regression from data that is not contained within the cross-validation loop (X1), (iii) fold-wise confound regression (X2), following [62]. For Y, there are two options, (i) no deconfounding (Y0), or (ii) learning deconfounding regression weights from the training subset, and then applying these weights on the left-out validation subset (Y1). For example, deconfounded 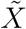 is calculated as 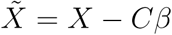 where C are confounds and *β* are regression weights. The *β* can be calculated as *β* = *C*^+^*X*, where *C*^+^ is the pseudoinverse of the confounds.

### 2.4. Training, validation and testing

For the Elastic Net, k fold nested cross-validation was performed (k=5). The subjects from both HCP and UKB data were divided into k folds. For each folds, a set was selected as the outer test set and the remaining k-1 sets were used as the outer training set. Then, an inner loop is used to tune the hyper-parameters via k-fold validation on the training set. The optimised model (using tuned hyper-parameters) was then evaluated on the test set and this procedure was repeated for all k test folds. In summary, the outer loop is used for model evaluation and the inner loop for the model selection phase. The important hyper-parameter to tune for Elastic Net is the weights on the ridge and lasso penalty. Lastly, we considered the family structure for HCP data and the family members were kept in the same folds. A summary figure explaining the process of nested cross-validation is shown in Figure A.34.

In case of CNNs, there are a number of hyper-parameters to tune, e.g., learning rate, number of hidden units, width of convolutional kernel, etc. For our benchmark study, we have hundreds of different variants of functional connectivity estimates as input, e.g., one possible variant of functional connectivity is estimated by following steps: step-1 (ICA parcellation), step-2 (SGGGM), step-3 (tangent space), step-4 (shrinkage in tangent space), step-5 (X1Y1 as deconfounding). This computationally makes it impossible to tune hyper parameters separately for each input. We randomly selected a few input variants (∼20) and tune the hyper-parameters for these selected inputs (for all non-imaging variables prediction). This gave us the estimate of hyper-parameter values for which CNNs perform optimally (convergence and relative higher prediction accuracy). We then fixed these hyper-parameters values for all non-imaging variables, prediction for both UKB and HCP datasets for all functional connectivity estimates. In Section 4, we performed external validation of the pipeline, and these hyper-parameters were still fixed on un-seen new datasets. This approach has the disadvantage that it can negatively affect CNN performance but it does avoid over-fitting concerns, and also tests the generalisability of the CNN architecture. The fine-tuning was performed within the training-validation-testing framework. The tuned hyper parameters for CNNs are reported in Table A.4.

We took advantage of large N of UKB (and HCP) and did not apply the inner cross-validation framework for CNN evaluation, but used the repeated training-validation-test splits. The subjects were split into training (72%), validation (8%) and testing (20%) sets. The training, validation and testing split was repeated 5 times; therefore, each model was trained five times and all subjects were tested once. The validation set here was used to assess performance of CNN, e.g., early stopping (when error on validation set grows but can select the previous optimum model) and model preparation (deep features selection) etc. It should be noted that any pre-processing of connectivity estimates (e.g., normalization) is performed within the cross-validation/training-validation-testing folds, and all CNN based results reported in this paper are on the testing data.

### 2.5. Statistical testing

#### Paired t-test

A paired t-test is performed to determine whether there is statistical evidence that the mean difference between paired observations (e.g., no shrinkage vs shrinkage approaches) on a particular outcome (i.e., fluid intelligence score prediction) is significantly different from zero. We have used symbols to report the test results; ns (not significant) when *P* > 0.05, * when *P* ≤ 0.05, ** when *P* ≤ 0.01 and *** when *P* ≤ 0.001.

#### Fisher transformation

To generate confidence intervals on the prediction correlations (i.e., continuous outputs e.g., age), we computed the Fisher transformation. If the prediction correlation is r, then Fisher transformation F(r) is *arctanh*(*r*). F(r) approximately follows a normal distribution, with mean = F(p) = *arctanh*(*p*), and standard error, 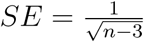, where n is sample size (i.e., number of subjects) and p is the true correlation coefficient. Then z-score is calculated as 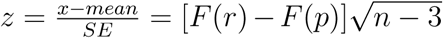. The confidence interval (CI) for p can be calculated as in Equation 11. The inverse Fisher transform is then calculated to bring the CI to the r units.

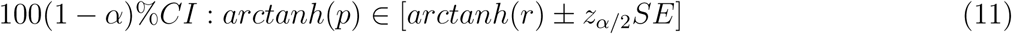

#### Wilson test

To draw the confidence intervals on the prediction accuracies (i.e., discrete outputs such as sex), we performed the Wilson test (suitable for the binomial distribution). The Wilson score interval is explained in [63, 64].

#### Repeated k-fold cross validation

We repeated 100 times the nested 5-fold cross validation on the top performing pipelines. We used the 99% confidence limits on these 100 prediction values to generate the confidence intervals. The rationale is to probe well the sampling variability and to test reproducibility of the top performing pipelines.

### 2.6. Imaging Data

The rfMRI data of 13301 subjects from UKB and 1003 subjects from the HCP were used in this analysis. The pre-processing pipelines for UKB and HCP are described in [65] and [66] respectively, although the main steps can be summarized as (1) motion correction, (2) removal of structural artifacts with ICA (Independent Component Analysis and FMRIB’s ICA-based X-noisefier (FIX), and (3) brain parcellation using any of the techniques covered in Section 2.1. The length of the scanning session for UKB is 6 minutes, and for HCP is 1 hour (4 sessions of 15 mins each). For UKB, we have 490 time-points and for HCP, we used data from all 4 sessions which gave us 4800 time points in total (1200 timepoints in each session).

### 2.7. Non-Imaging Data

Here, we have chosen four non-imaging variables: age, sex, fluid intelligence score and neuroticism score (the latter only for UKB), for following reasons:

#### Age

Improving prediction of age from brain data is of interest, to further clinical understanding of how age links to neurodevelopment and degeneration. For instance, comparing the discrepancy between an individual’s chronological age and the age predicted from neuroimaging data can serve as a biomarker for several age-linked brain diseases [67]. Age can be accurately predicted from structural MRI data [68]; however, prediction of age from functional connectivity estimates is not as accurate. Nevertheless, this represents a non-imaging variable that can be predicted reasonably strongly (as opposed to fluid intelligence and neuroticism).

#### Sex

Sex classification through functional connectivity is comparatively accurate, and so this variable provides a valuable evaluation parameter to compare benchmark performance of different models.

#### Fluid Intelligence

Fluid intelligence refers to the ability to reason and to solve new problems independently of previously acquired knowledge [69]. It is highly related to professional and educational success and represents overall performance across a broad range of cognitive abilities. A number of recent studies have predicted fluid intelligence from functional connectivity [70], but due to measurement variability, correlations remain rather weak. However, more precise and novel methods for the estimation and use of functional connectivity could lead to better predictions, and provide a more precise understanding of how brain networks relate to cognitive abilities.

#### Neuroticism

Mental health is being assessed in a number of ways within UKB. Participants answer a 12-item neuroticism questionnaire as part of their baseline assessment. We used the algorithm employed in [71] to calculate the score of neuroticism for each subject.

For the purpose of this paper, we have shown the prediction accuracy (for categorical variables) and prediction correlation (for continuous variables). There could be an argument in favor of variance (or coefficient of determination) over correlation. However, correlation is commonly used and well-understood by the readers and that was our rationale behind selecting correlation (as we want this pipeline paper to be directly comparable to existing literature work). Also, for the reported results in this paper, correlation or coefficient of determination carried exactly the same information.

## 3. Results

### 3.1. Parcellation and Functional Connectivity Estimation

Figure 2 shows the prediction accuracy/correlation for non-imaging measures from functional connectivity estimates across different parcellation schemes (Section 2.1) and functional connectivity estimation methods (Section 2.2). The results in Figure 2 were calculated by directly vectorizing the connectivity estimates without projection into tangent space. Elastic Net was used as a predictor/classifier with nested 5-fold cross-validation. The results illustrated in Figure 2 are a reduction of the full set of results, chosen to highlight the performance of methods employed in step 1 (brain parcellation) and step 2 (functional connectivity estimation) using the relatively robust Elastic Net as the predictor/classifier. The results shown in Figure 2 were calculated after removing the effect of confounds from data (deconfounding), and Figure A.14 shows prediction estimates without confound removal.

**Figure 2:**
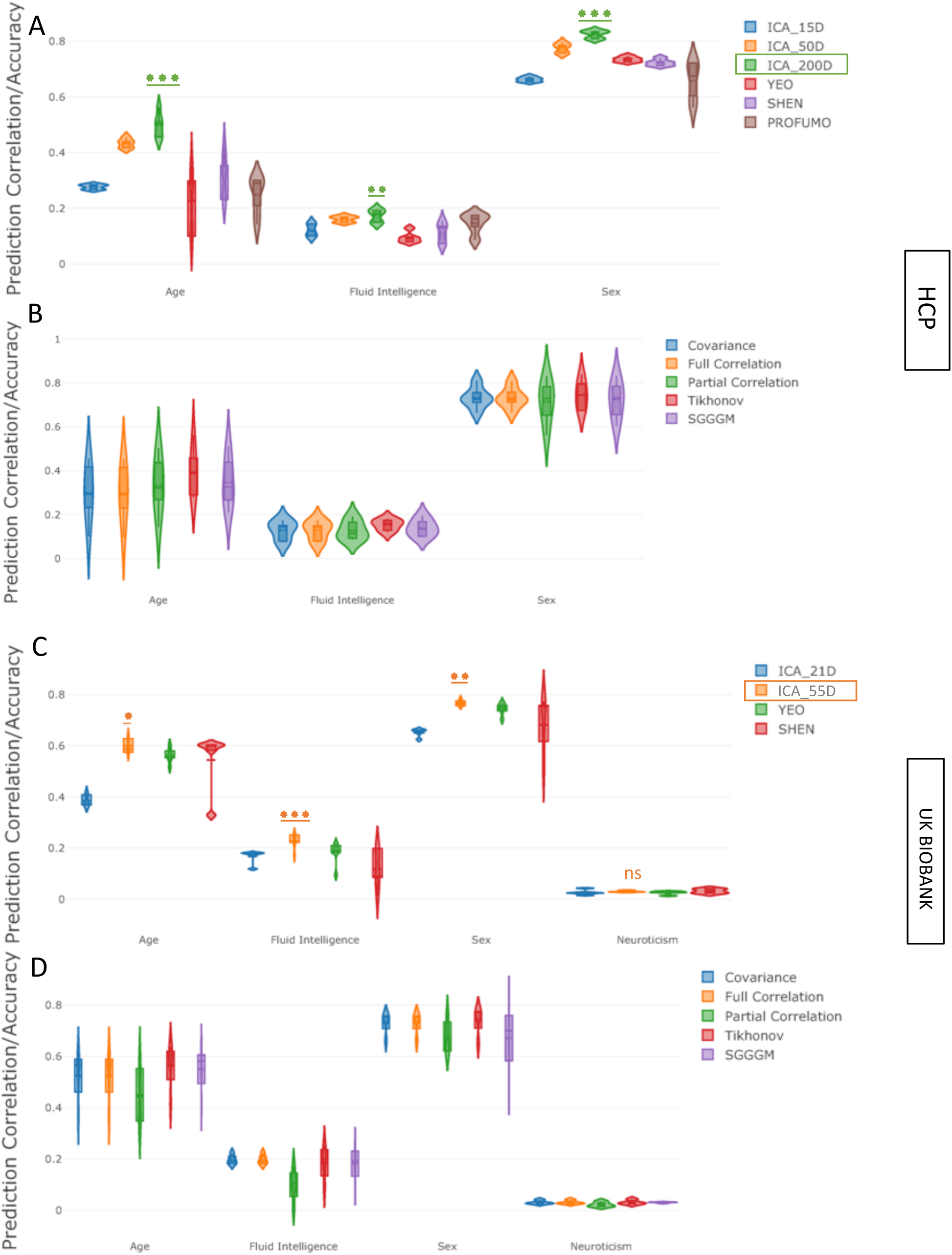
The impact of various parcellation strategies and functional connectivity estimation methods on prediction power for non-imaging variables **after deconfounding. [A**,**B] (HCP Data), [C**,**D] (UKB Data):** The violin plots in **A** and **C** show the prediction variability over 5 measures of functional connectivity estimates and in **B** and **D** show the prediction variability the over different parcellation schemes. For HCP, the ICA based parcellation schemes are *ICA*_15*D, ICA*_50*D* and *ICA*_200*D*, and for UKB are *ICA*_21*D* and *ICA*_55*D*, where D = the number of parcels. For both HCP and UKB, SHEN parcellation was 268D, YEO was 100D, and PROFUMO was 50D (for HCP only). The stars refer to comparison against the next-best method.

### 3.2. Tangent Space Projection and Shrinkage

A reference point is required when projecting functional connectivity estimates into the tangent space. This reference mean binds each of the connectivity estimates into the same tangential plane. The results in Figure A.15 show the prediction correlation for fluid intelligence using different reference means. For each mean, functional connectivity estimates from all parcellation schemes (Section 2.1) were estimated by using all functional connectivity estimation methods (Section 2.2). The predictor/classifier employed here for fluid intelligence prediction was Elastic Net using nested 5 fold cross validation. For simplicity, only fluid intelligence prediction scores are displayed in Figure A.15, as the prediction correlations for other non-imaging variables follows the same trend across various reference mean estimation techniques. The violin plots in Figure A.15 show the prediction variability over 4 different parcellation schemes and 5 measures of functional connectivity estimates in the tangent space. The results shown in Figure A.15 were calculated after deconfounding, and Figure A.16 shows prediction estimates without deconfounding.

Figure 3 (sub-figures A,C,E and G) illustrates how the prediction accuracy was modified by projecting connectivity estimates into the tangent plane. Again, we varied the parcellation schemes and functional connectivity estimation techniques, but fixed Elastic Net as the predictor/classifier. The results illustrated in Figure 3 (sub-figures A,C,E and G) are a reduction of the full set of results, chosen to highlight the performance of tangent space parameterization (Section 2.2.2). Figure 3 (sub-figures B,D,F and H) compares the effect of applying shrinkage in the tangent space (Section 2.2.3). The shrinkage applied was non-isotropic shrinkage towards the population dispersion (PoSCE). The Results in Figure 3 (sub-figures B,D,F and H) are based on functional connectivity estimates that have been projected into the tangent plane (Section 2.2.2). The violin plots in Figure 3 (sub-figures B,D,F and H) show the prediction variability over 4 different parcellation schemes and 5 measures of functional connectivity estimates in the normal tangent space (no shrinkage) versus tangent shrinkage space (PoSCE), (Section 2.2.3). Similarly to the previously displayed results, Elastic Net was used as the predictor/classifier for prediction of non-imaging measures. Essentially, Figure 3 shows side-by-side the effect of tangent space projection (from ambient space to tangent space) and tangent space shrinkage (staying within tangent space). The length of the scanning session in HCP is 60 minutes, Figure 3 also shows the prediction performance on the original 60 minutes scan, and on cut-down data (first 15 minutes and first 5 minutes of the scan for each subject). Moreover, the results shown in Figure 3 were calculated after deconfounding, and Figure A.17 shows similar prediction estimates without deconfounding (for the full length scanning session only).

**Figure 3:**
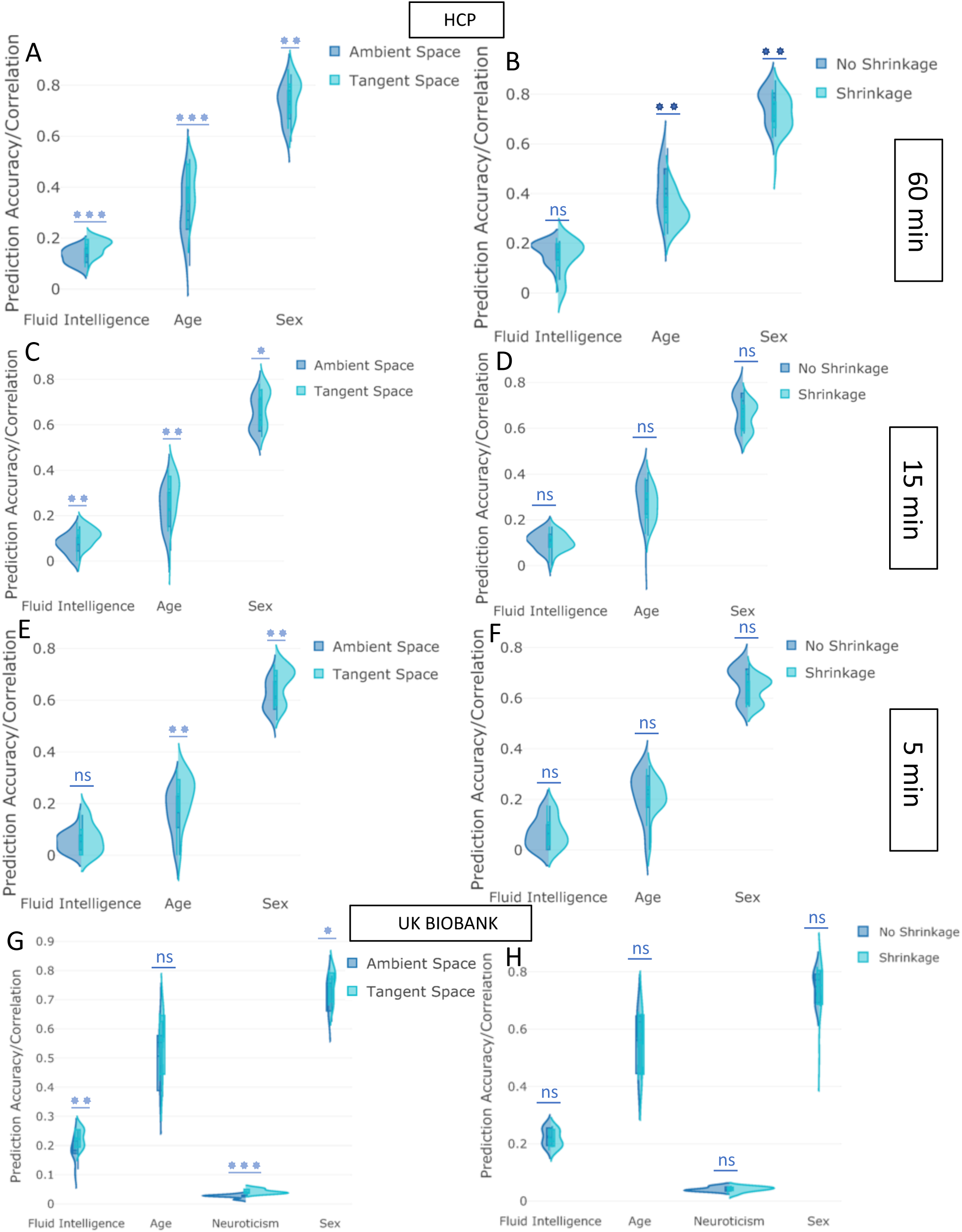
The impact of projecting to tangent space and applying shrinkage in tangent space on prediction power **after deconfounding. [A-F] (HCP Data), [G**,**H] (UKB):** The y-axis depicts the prediction accuracy/correlation for different behavioural measures. “Tangent Space” means that tangent space projection was applied on functional connectivity estimates (originally in the “Ambient Space”). The “Shrinkage” strategy means that non-isotropic PoSCE shrinkage was applied to connectivity estimates in tangent space before feeding to the predictor/classifier. “No Shrinkage” means that projected functional connectivity estimates in tangent space were directly fed to the predictor/classifier, and did not undergo PoSCE shrinkage. The violin plots show the prediction variability over 4 different parcellation schemes and 5 measures of functional connectivity estimates.

The Bland-Altman plot is shown in Figure A.18 to describe agreement between shrinkage versus no shrinkage approaches. The Figure 3 (B and H sub-figures) and Figure A.18 are precisely two different ways of visualizing the similar results. Figure A.19 compares the performance of isotropic (Ledoit-Wolf) versus non-isotropic (PoSCE) shrinkage in the tangent space and Figure A.20 compares the performance of isotropic (Ledoit-Wolf) versus no shrinkage strategy (projected functional connectivity estimates in the tangent space were directly fed to the predictor/classifier, and did not undergo any shrinkage). All the results displayed in Figure 3, A.18, A.19 and A.20 were calculated after deconfounding. The violin plots in Figure A.21 show the effect of varying the dimensionality of parcellation on shrinkage approaches for 5 measures of functional connectivity estimates, e.g., it compared the effect of tangent space shrinkage for higher dimensional ICA vs low dimensional ICA.

### 3.3. Deconfounding

We compared various deconfounding strategies (Section 2.3.1), by varying ICA dimensionality (Section 2.1.3) and using Elastic Net as the predictor/classifier. Figure 4 shows the prediction correlations of fluid intelligence from functional connectivity estimates for each deconfounding strategy. The violin plots in Figure 4 shows the prediction variability over each specific sICA dimensionality, 5 measures of functional connectivity estimates and 3 different spaces (ambient space, tangent space and tangent shrinkage space). Again, the results illustrated in Figure 4 are a reduction of the full set of results, chosen to highlight the performance of different deconfounding strategies. (Section 2.3)

**Figure 4:**
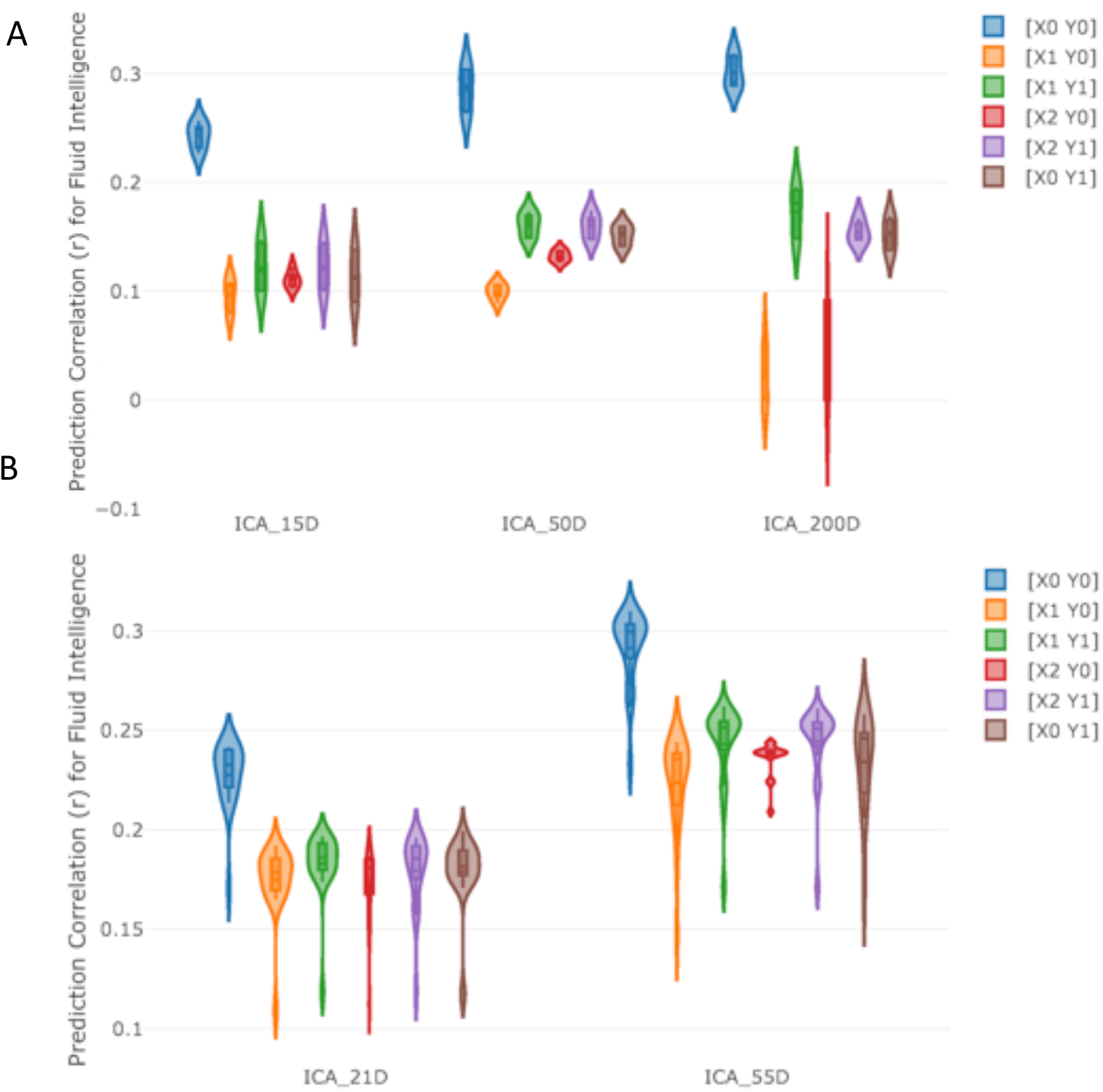
The impact of deconfounding strategies on the prediction power. **[A] (HCP), [B] (UKB)** This figure shows the prediction correlation for fluid intelligence score for various ICA based parcellation techniques across the 6 possible deconfounding strategies that were tested. The violin plots show the prediction variability over each specific sICA dimensionality, 5 measures of functional connectivity estimates and 3 different spaces (ambient space, tangent space and tangent shrinkage space). X is an input matrix (i.e, subjects’ functional connectivity), and Y, a response vector (e.g., fluid intelligence). The details of different deconfounding strategies are explained in Section 2.3.1.

### 3.4. Predictors/Classifiers Performance

Figure 5 compares the performance of different predictors (Section 2.3) on the prediction between functional connectivity and non-imaging variables. Inputs included functional connectivity estimates (Section 2.2) both in ambient space (without tangent space projection) and tangent space (Section 2.2.2). Estimates with tangent space parameterization included estimates with and without PoSCE shrinkage (tangent space and tangent shrinkage space). Therefore, the violin plots in Figure 5 show the prediction variability over 4 different parcellation schemes, 5 measures of functional connectivity estimates and 3 different spaces (ambient space, tangent space and tangent shrinkage space). Again, the results illustrated in Figure 5 are a reduction of the full set of results, chosen to highlight the performance of different classifiers/predictors (Section 2.3). The results in Figure 5 compared the performance of Elastic Net, 2D RCNN and BrainNetCNN (Section 2.3). We included GraphCNN initially, but dropped it due to consistently poor performance compared with the other models. Figure A.31 shows the comparative performance of GraphCNN and Elastic Net. The subset of results chosen for Figure A.31 were the ones where GraphCNN exhibited the best performance, although still underperforming in comparison to Elastic Net.

**Figure 5:**
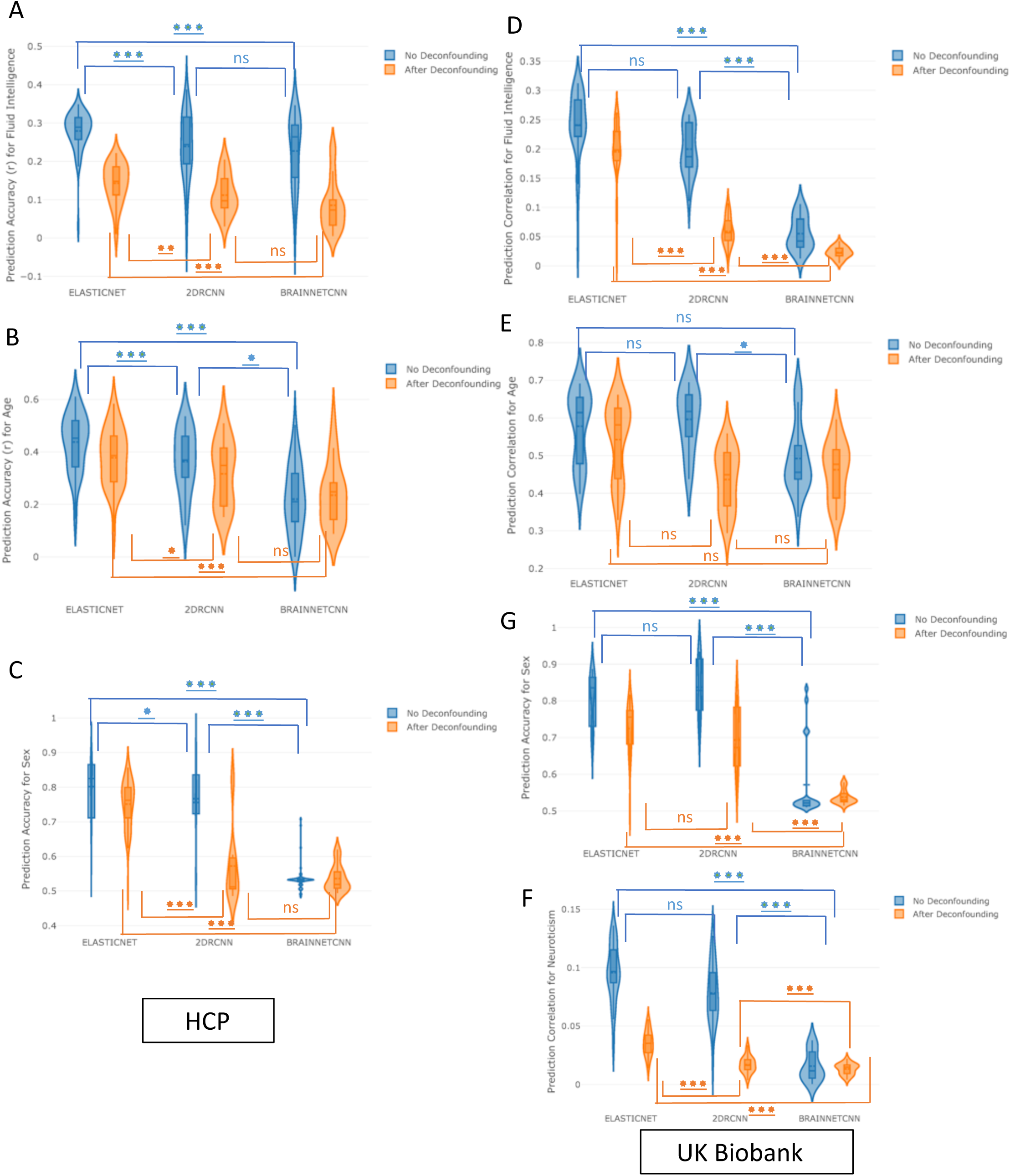
The performance of different classifiers/predictors on prediction power with and without deconfounding. **[A**,**B**,**C] (HCP Data), [D**,**E**,**F**,**G] (UKB Data):** The prediction accuracy/correlation for different non-imaging measures is depicted on the y-axis. The violin plots show the prediction variability over 4 different parcellation schemes, 5 measures of functional connectivity estimates and 3 different spaces (ambient space, tangent space and tangent shrinkage space).

### 3.5. Top Performing Configurations

Figures 6 and 7 show the top ten methods in terms of predictive power, with strategies from different steps identified. The results shown in Figure 6 and 7 were calculated after deconfounding, and Figures A.22 and A.23 show prediction estimates without deconfounding. The error bars in Figures 6, 7, A.22 and A.23 were generated using the Fisher transformation (for continuous output) and Wilson test (for discrete output) (Section 2.5). The highlighted red blocks show the recommended pipelines (rationale explained in Section 5), and the red dotted lines highlight the point when the error bar of pipeline after the dotted line is out of range from the error bar of the top (first) pipeline.

**Figure 6:**
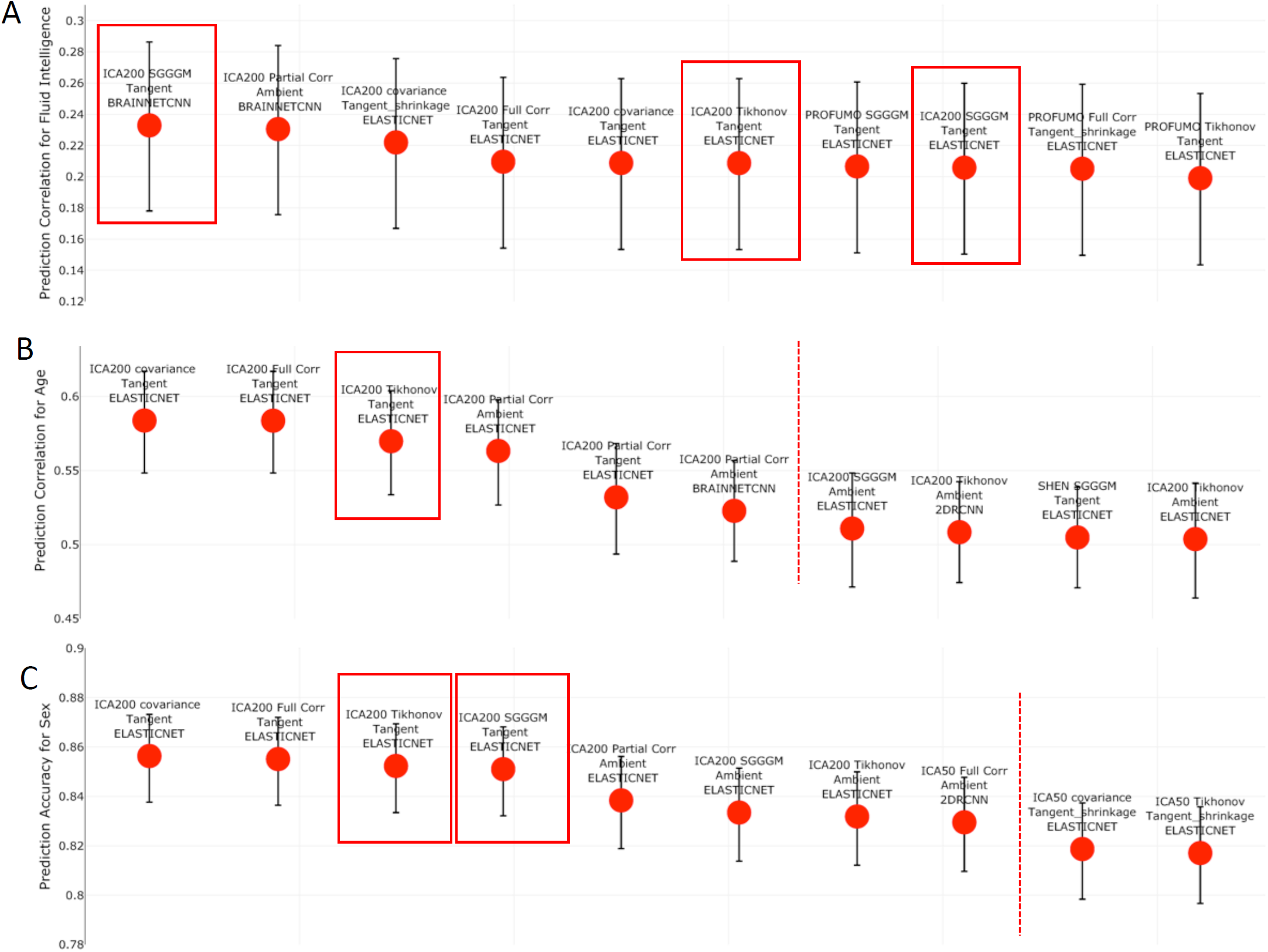
The top performing ten configurations for the prediction of each non-imaging variable by dataset **after deconfounding. [A**,**B**,**C] (HCP Data)** Each data point represents a different configuration strategy that may vary in terms of parcellation strategy, the functional connectivity estimation method, whether tangent space parameterization was employed, whether tangent space regularization was employed, and the predictor that was used. The first word indicates the parcellation strategy, and the second word refers to the functional connectivity estimation method. The third word refers to the geometry in which classifier/predictor is applied, ambient referring to non-tangent space and tangent referring to the projected covariance matrices in tangent space. If non-isotropic shrinkage was applied after projecting covariance matrices to tangent space, the fourth word will be “shrinkage”. The last word indicates the type of classifier/predictor that was used. The highlighted red blocks show the recommended pipelines (rationale explained in Section 5), and red dotted lines highlight the point when the error bar of pipeline after the dotted line is out of range from the error bar of the top (first) pipeline.

**Figure 7:**
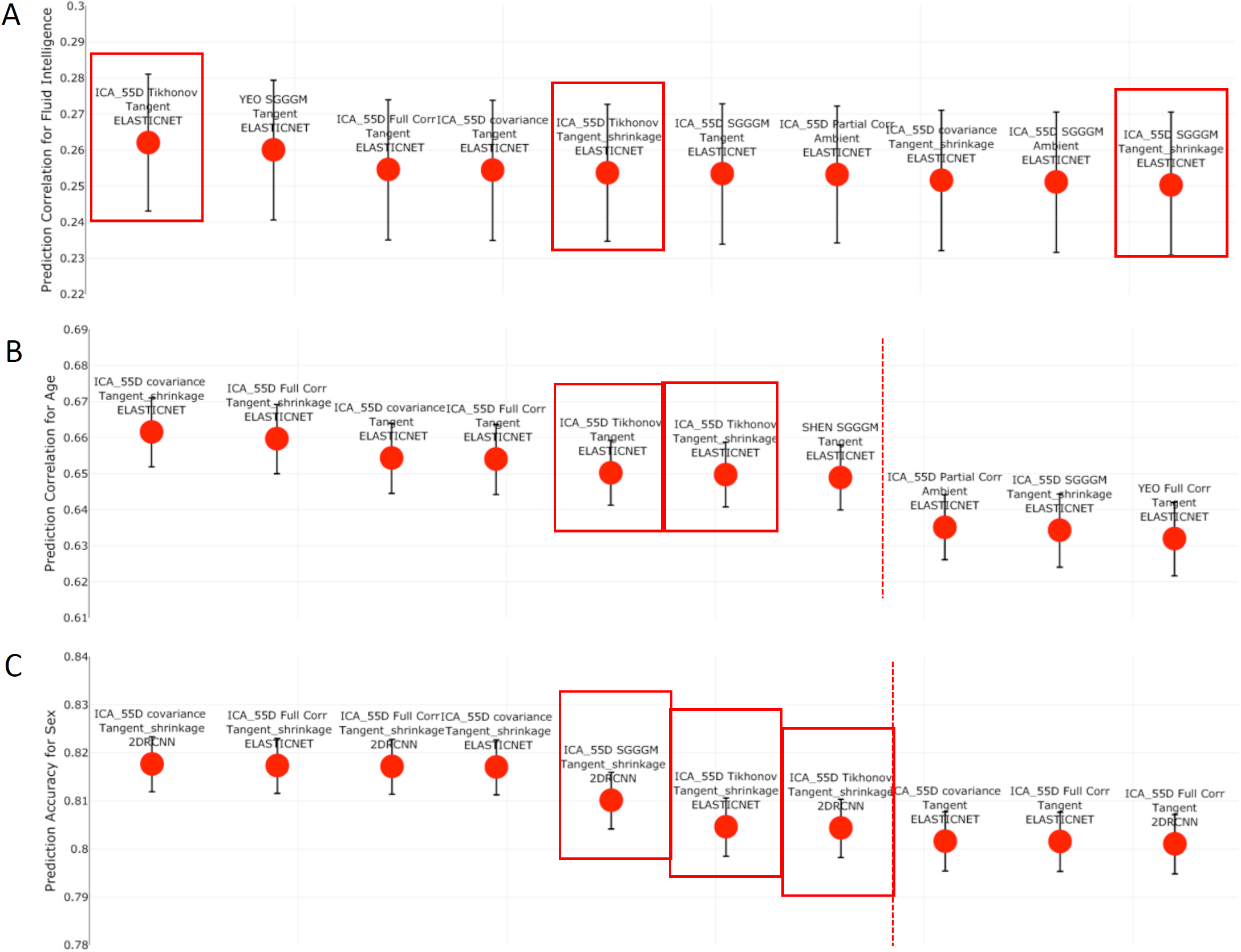
The top performing ten configurations for the prediction of each non-imaging variable by dataset **after deconfounding. [A**,**B**,**C**,**D] (UKB Data)** Each data point represents a different configuration strategy that may vary in terms of parcellation strategy, the functional connectivity estimation method, whether tangent space parameterization was employed, whether tangent space regularization was employed, and the predictor/classifier that was used. The first word indicates the parcellation strategy, and the second word refers to the functional connectivity estimation method. The third word refers to the geometry in which classifier is applied, ambient referring to non-tangent space and tangent referring to the projected covariance matrices in tangent space. If non-isotropic shrinkage was applied after projecting covariance matrices to tangent space, the fourth word will be “shrinkage”. The last word indicates the type of classifier/predictor that was used. The highlighted red blocks show the recommended pipelines (rationale explained in Section 5), and red dotted lines highlight the point when the error bar of pipeline after the dotted line is out of range from the error bar of the top (first) pipeline.

### 3.6. Joint Results

Figures 8, 9 and 10 show the combination of various techniques at different stages of the pipeline for fluid intelligence, sex, and age prediction respectively for both HCP and UKB data after deconfounding. Figures A.24, A.25, A.26 show similar results for fluid intelligence, sex and age prediction respectively but without deconfounding. In each of these figures, a few parameters were fixed across the different pipeline combinations (Euclidean mean as reference mean, non-isotropic shrinkage as shrinkage in tangent space). Figure A.27 provides results for the prediction of neuroticism score in UKB data, and results are displayed both before and after controlling for confounds.

**Figure 8:**
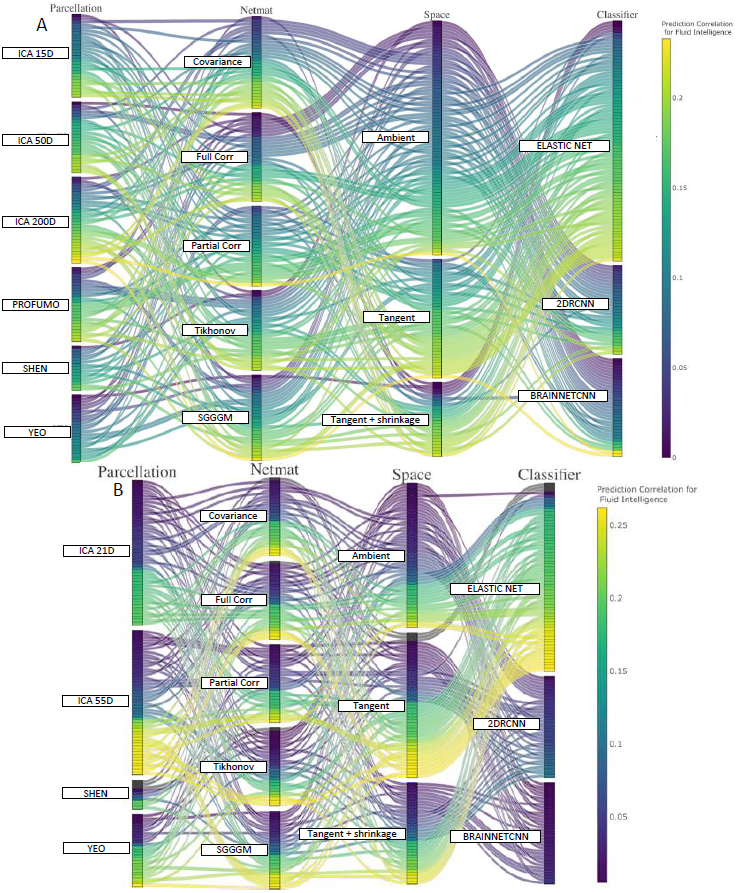
This parallel coordinates plot provides a visualization of all possible combinations of options in the pipeline to predict fluid intelligence scores from functional connectivity **after deconfounding. [A] (HCP Data), [B] (UKB Data):** The lines are color-coded according to their prediction performance. The vertical ordering is chosen with respect to prediction performance for each sub-block (in ascending order of prediction correlation/accuracy). For example, the lower (from bottom) the position of lines in a block, the higher the prediction accuracy of that variant (and vice-versa). With respect to the vertical ordering of sub-blocks (e.g., ambient or tangent space), that is chosen by hand (but it should not make any difference as the crossing of different lines does not explain any meaningful information here).

**Figure 9:**
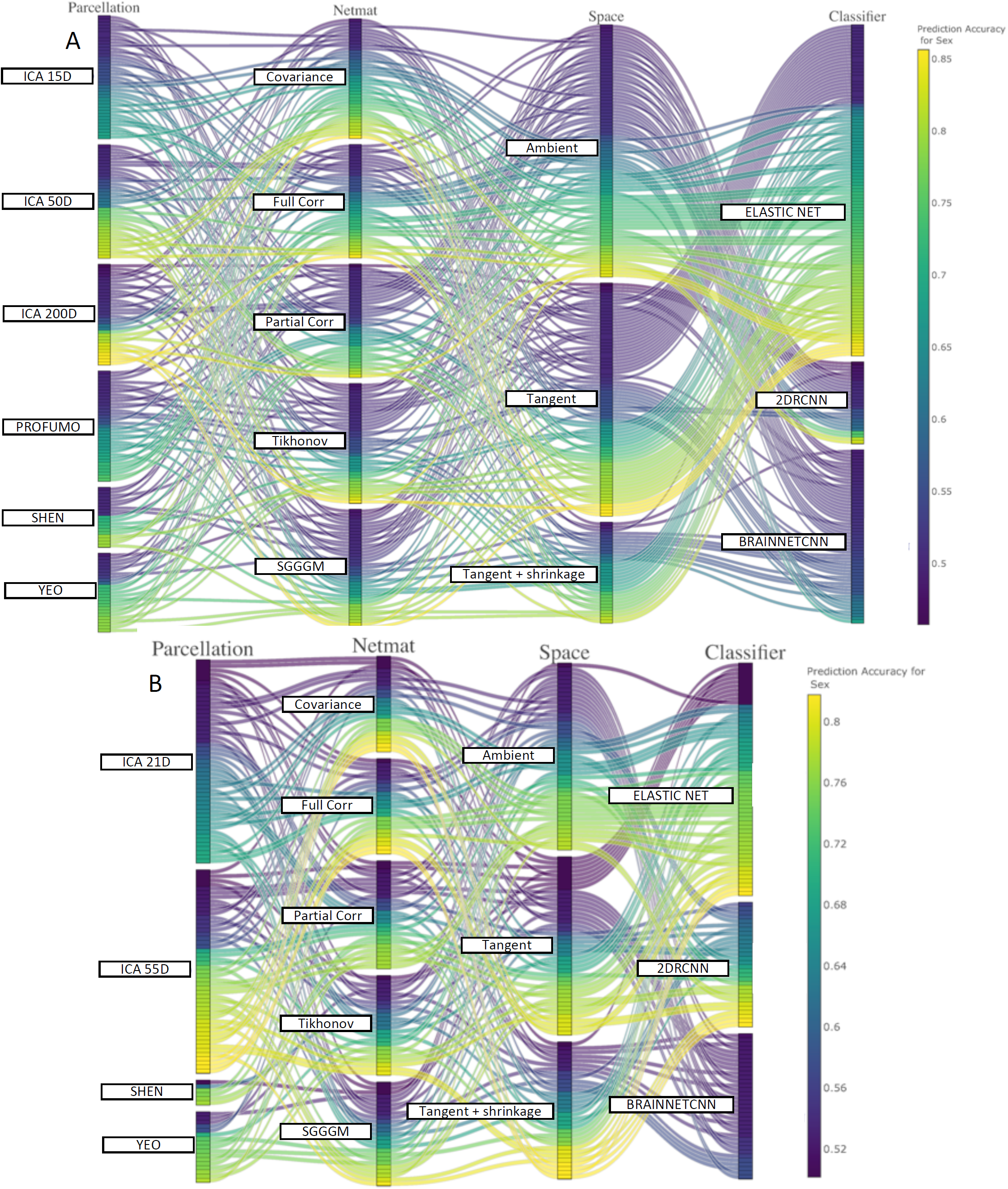
This parallel coordinates plot provides a visualization of all possible combinations of options in the pipeline to pipeline to predict sex from functional connectivity **after deconfounding. [A] (HCP Data), [B] (UKB Data):** The lines are color-coded according to their prediction performance.

**Figure 10:**
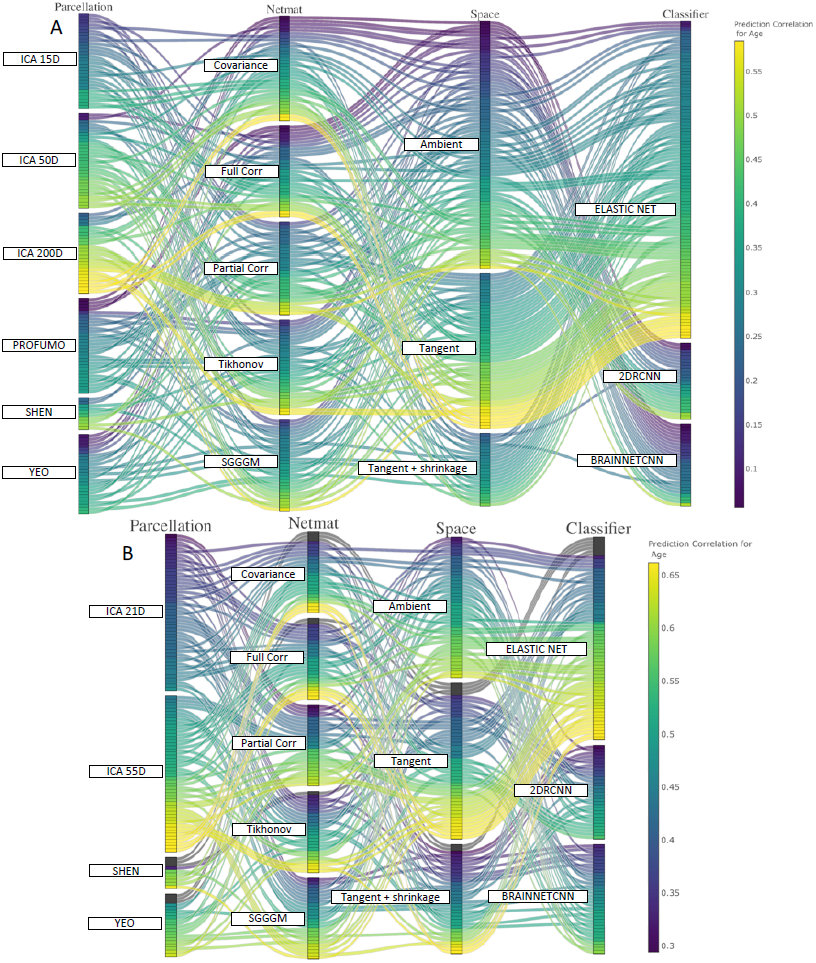
This parallel coordinates plot provides a visualization of all possible combinations of options in the pipeline to pipeline to predict age from functional connectivity **after deconfounding. [A] (HCP Data), [B] (UKB Data):** The lines are color-coded according to their prediction performance.

### 3.7. Deconfounding revisited

All the results illustrated in this paper were using X1Y1 or X1Y0 deconfounding strategy (apart from Figure 4 where all six deconfounding strategies were employed). We have applied the X1Y1 deconfounding for age, fluid intelligence and neuroticism score prediction and X1Y0 for sex prediction (X1Y1 and X1Y0 are detailed in Section 2.3.1 and the reasoning for this is discussed in Section 5). Confounds (particularly head size and motion) are clearly a serious problem if not dealt with for fluid intelligence score prediction. Head size is positively correlated with fluid intelligence and head motion is negatively correlated (detailed further in Table A.5).For neuroticism score prediction, the confounds which have the most effect on prediction power are age, table position, imaging-centre and date-drift related confounds. This could be because of non-linear interactions between imaging-centre/date-drift related confounds, age, table-position and neuroticism. For age, the crucial confound is head-motion (positively correlated with age), although other than head-motion, we did not find prediction was significantly affected by any confounds. For sex prediction, the crucial confounds are age, height, weight, head motion and head size. For fluid intelligence, neuroticism and age prediction, the deconfounded results are calculated after regressing out all the potential confounding variables. However, in case of sex prediction, for the deconfounded results, we regressed out all the confounds from data except head-size and body density related confounds. The rationale is that sex is a major casual factor for head-size, and head-size causes some confounding effects in the imaging. Regressing out head-size reduces the overall sex prediction accuracy but allows one to focus on the non-headsize sex effects in the imaging. However, one important goal of this work is to optimize functional connectivity estimation methods (i.e., allow maximum predictors/classifiers to converge to make performance of different models comparable). Therefore, we didn’t include head-size and body density related variables as confounds when predicting sex. Most of the results without removing the effect of confounds (i.e., higher prediction accuracy/correlation as compared to deconfounded results in the main text) are shown in the Inline supplementary material.

## 4. Results Evaluation

### 4.1. Evaluation on Independent Datasets

We evaluated the proposed network modelling methods on Autism Brain Imaging Data Exchange (ABIDE) [72] and Addiction Connectome Preprocessed Initiative (ACPI) datasets ^8^.

#### ABIDE

We used the pre-processed resting-state fMRI data from the Preprocessed Connectome Project pipeline (C-PAC) [73], without global signal regression and without band-pass filtering. The processed resting-state fMRI data is available for 871 subjects (autism and control). We applied most of the pipeline combinations on ABIDE including all parcellation strategies (except PROFUMO), functional connectivity estimation techniques, tangent space parameterization and shrinkage techniques (non-isotropic shrinkage), and predictors/classifiers (except GraphCNN). We pre-selected our recommended pipelines based on the generally top performing techniques as shown in Figures 6, 7, A.22 and A.23 (rationale further explained in Section 5); parcellation (ICA), functional connectivity estimation (Tikhonov or SGGGM), tangent space parameterization (yes) and tangent space shrinkage (optional). Regarding recommended predictor/classifier, Elastic Net tends to be more stable but CNN-based architectures can out-perform Elastic Net sometimes, therefore; we tested all the predictors/classifiers. The confounds were not provided for the ABIDE data-set so results reported are based without deconfounding (X0Y0). ABIDE data is comparatively more noisy as compared to HCP and UKB data; the number of time points varies between 115 and 295. We employed predictors to predict age of participants (min = 6.47, max = 64).

#### ACPI

We used the pre-processed resting-state fMRI data from Multimodal treatment study of Attention Deficit Hyperactivity Disorder. The data was processed using the Preprocessed Connectome Project pipeline (C-PAC); with images registered using Advanced Normalization Tools (ANTS), without any scrubbing, and without global signal regression. We applied most of pipeline combinations on ACPI data including all parcellation strategies (except PROFUMO), functional connectivity estimation techniques, tangent space parameterization and shrinkage techniques, and predictors/classifiers (except GraphCNN). The confounds were not provided for ACPI data-set so results reported here are without deconfounding (X0Y0). The number of time points in ACPI dataset for each participant is 179 and the total number of subjects is 126. We employed predictors/classifier to predict whether a subject smokes or not (binary classification).

The results for predicting age on ABIDE and smoking status on ACPI for the chosen subset of pipelines is shown in Table 3. The top 3 best performing pipelines for both ABIDE and ACPI are highlighted in red. Figures A.33 and A.29 show the full set of results for age and smoking status prediction for both ABIDE and ACPI data respectively.

**Table 3:**
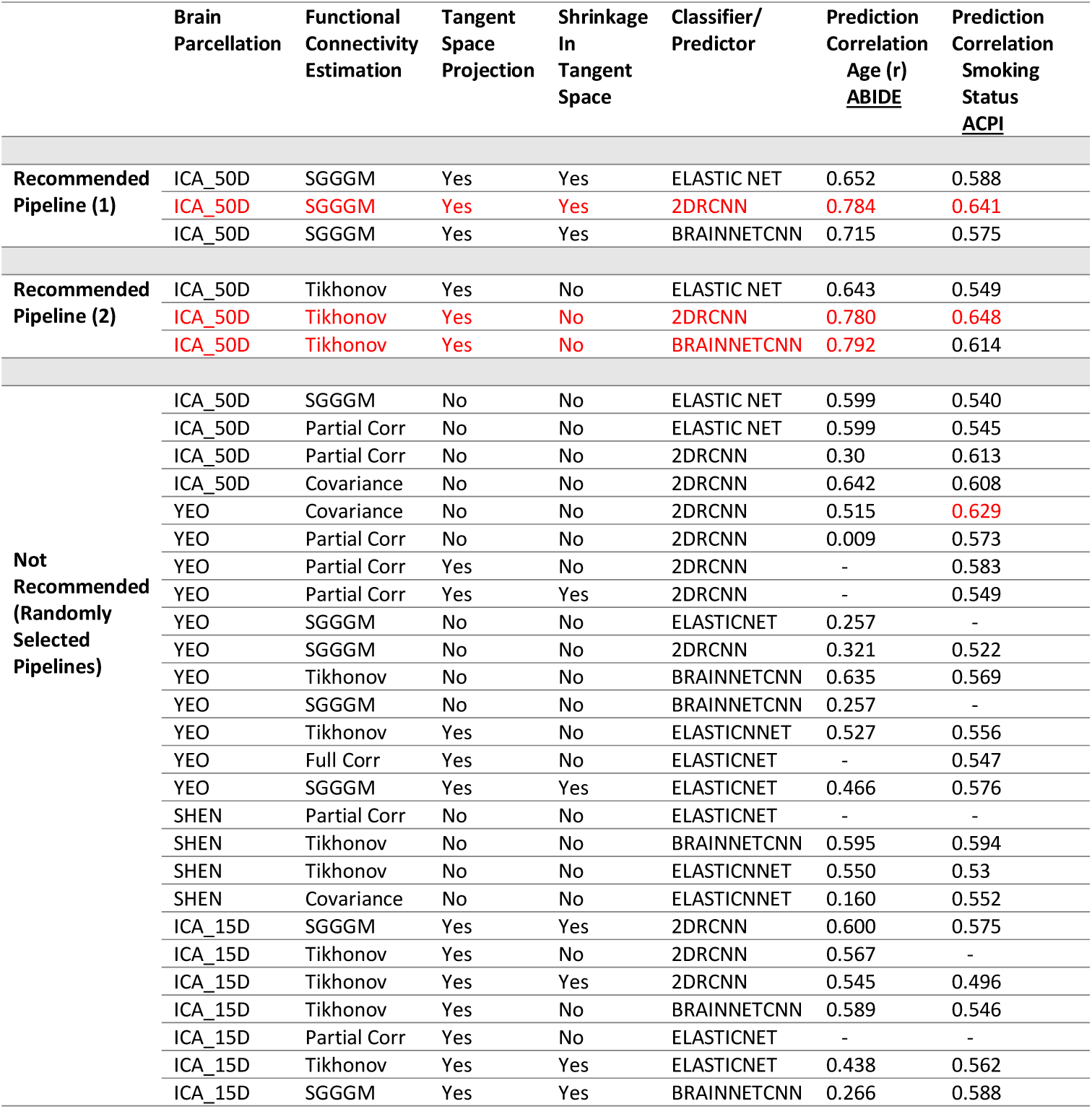
Predicting age for ABIDE participants and smoking status for ACPI participants. The top 3 best performing pipelines for both ABIDE and ACPI is highlighted in red. The missing lines indicate that predictors/classifier did not converge.

|  | Brain<br>Parcellation | Functional<br>Connectivity<br>Estimation | Tangent<br>Space<br>Projection | Shrinkage<br>In<br>Tangent<br>Space | Classifier/<br>Predictor | Prediction<br>Correlation<br>Age (r)<br><u>ABIDE</u> | Prediction<br>Correlation<br>Smoking<br>Status<br><u>ACPI</u> |
| --- | --- | --- | --- | --- | --- | --- | --- |
| **Recommended**<br>**Pipeline (1)** | ICA_50D | SGGGM | Yes | Yes | ELASTIC NET | 0.652 | 0.588 |
|  | ICA_50D | SGGGM | Yes | Yes | 2DRCNN | 0.784 | 0.641 |
|  | ICA_50D | SGGGM | Yes | Yes | BRAINNETCNN | 0.715 | 0.575 |
| **Recommended**<br>**Pipeline (2)** | ICA_50D | Tikhonov | Yes | No | ELASTIC NET | 0.643 | 0.549 |
|  | ICA_50D | Tikhonov | Yes | No | 2DRCNN | 0.780 | 0.648 |
|  | ICA_50D | Tikhonov | Yes | No | BRAINNETCNN | 0.792 | 0.614 |
| **Not**<br>**Recommended**<br>**(Randomly**<br>**Selected**<br>**Pipelines)** | ICA_50D | SGGGM | No | No | ELASTIC NET | 0.599 | 0.540 |
|  | ICA_50D | Partial Corr | No | No | ELASTIC NET | 0.599 | 0.545 |
|  | ICA_50D | Partial Corr | No | No | 2DRCNN | 0.30 | 0.613 |
|  | ICA_50D | Covariance | No | No | 2DRCNN | 0.642 | 0.608 |
|  | YEO | Covariance | No | No | 2DRCNN | 0.515 | 0.629 |
|  | YEO | Partial Corr | No | No | 2DRCNN | 0.009 | 0.573 |
|  | YEO | Partial Corr | Yes | No | 2DRCNN | - | 0.583 |
|  | YEO | Partial Corr | Yes | Yes | 2DRCNN | - | 0.549 |
|  | YEO | SGGGM | No | No | ELASTICNET | 0.257 | - |
|  | YEO | SGGGM | No | No | 2DRCNN | 0.321 | 0.522 |
|  | YEO | Tikhonov | No | No | BRAINNETCNN | 0.635 | 0.569 |
|  | YEO | SGGGM | No | No | BRAINNETCNN | 0.257 | - |
|  | YEO | Tikhonov | Yes | No | ELASTICNET | 0.527 | 0.556 |
|  | YEO | Full Corr | Yes | No | ELASTICNET | - | 0.547 |
|  | YEO | SGGGM | Yes | Yes | ELASTICNET | 0.466 | 0.576 |
|  | SHEN | Partial Corr | No | No | ELASTICNET | - | - |
|  | SHEN | Tikhonov | No | No | BRAINNETCNN | 0.595 | 0.594 |
|  | SHEN | Tikhonov | No | No | ELASTICNET | 0.550 | 0.53 |
|  | SHEN | Covariance | No | No | ELASTICNET | 0.160 | 0.552 |
|  | ICA_15D | SGGGM | Yes | Yes | 2DRCNN | 0.600 | 0.575 |
|  | ICA_15D | Tikhonov | Yes | No | 2DRCNN | 0.567 | - |
|  | ICA_15D | Tikhonov | Yes | Yes | 2DRCNN | 0.545 | 0.496 |
|  | ICA_15D | Tikhonov | Yes | No | BRAINNETCNN | 0.589 | 0.546 |
|  | ICA_15D | Partial Corr | Yes | No | ELASTICNET | - | - |
|  | ICA_15D | Tikhonov | Yes | Yes | ELASTICNET | 0.438 | 0.562 |
|  | ICA_15D | SGGGM | Yes | Yes | BRAINNETCNN | 0.266 | 0.588 |

### 4.2. Comparison with other commonly used methodologies

#### Connectome-based predictive modelling

There is another pipeline in the literature known as Connectome-based predictive modelling (CPM) [95] that is mainly comprised of three steps: (i) parcellating the brain using the SHEN functional atlas, (ii) estimating connectivity using full correlation, (iii) correlating each edge in the connectivity matrix with behavioral measures, selecting only the most correlated N edges, averaging across these N edges to give a single predictor value for each subject, and then fitting a linear model (e.g., robust regression, polynomial curve fitting) to predict non-imaging variables.

#### Random forests

Random forests algorithm are an ensemble learning method and work as a large collection of decorrelated random trees [74]. We have compared the prediction performance of Elastic Net and Random Forest for sex prediction, and results are shown in Figure A.32.

#### Dictionary Learning

Massive online dictionary learning (MODL) was used to extract the time-series data [75] on ABIDE data-set only. These time-series were originally used in [75]. We have compared the prediction performance of ICA and Dictionary Learning for a subset of tests, and results are shown in Figure A.33.

#### Higher Dimensional YEO Parcellation

The YEO parcellation is available at multiple resolution levels. For most analysis in this work, we have used d = 100 parcels, but we have compared a subset of methodologies for YEO parcellation with d = 200 and d = 400 parcels. The results are shown in Figure 11 (sub-figure D).

**Figure 11:**
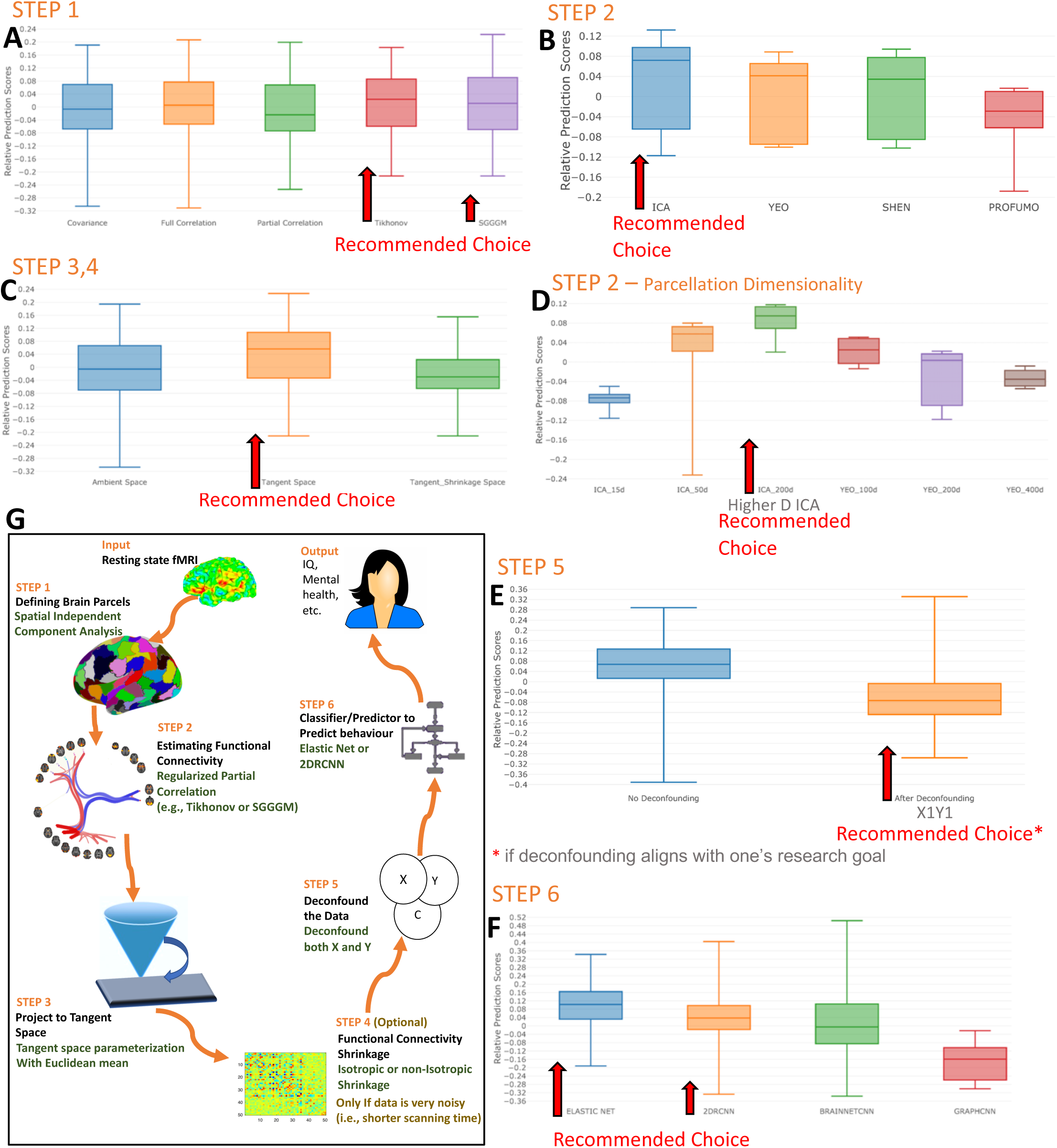
The relative impact of different steps of pipeline on the non-imaging variables prediction performance for participants in UKB, HCP, ABIDE, and ACPI datasets. Each sub-figure highlights the recommended options which are selected based on quantitative and qualitative analysis. The recommended optimal methodologies from each step of the pipeline are, (1) sICA for parcellation, (2) Tikhonov or SGGGM for functional connectivity estimation, (3) projecting the connectivity estimates to tangent space, (4) optional shrinkage in tangent space, (5) deconfounding the data (both the target and predictor variables) and (6) Elastic Net or 2D RCNN as a predictor/classifier.

## 5. Discussion

A number of research groups are increasingly focussing on rfMRI connectivity, and developing a plethora of methods [76, 77]. These methods range from neural-level simulations to abstract graph theoretic summaries of a connectivity matrix [78, 79]. Between these two extremes, we also have several popular network modelling methods (applied to real fMRI data) that can mainly be categorized into the study of functional vs effective connectivity (an over-simplistic division) [79, 80, 81]. In the latter, one is generally attempting to estimate both strengths and the dominant direction of information flow (causality); this aims to be more meaningful than functional connectivity, but is hard to estimate robustly on resting data, and such methods can only cope with fewer parcels. Then, we have functional connectivity (presented in this paper), which comes with the benefits of being simpler, well conditioned and being able to handle more nodes. Functional connectivity has also faced some criticism, as even with partial correlation, one can try to estimate the direct connections, but not the directionalities. Ideally, we would like to work at the more sophisticated bio-physically interpretable level, but considering the current limitations, functional connectivity is a pragmatic choice, particularly for resting-fMRI observational studies (because there are no external interventions such as in task-fMRI). In this paper, we aimed to identify an optimal pipeline to estimate and analyze functional connectivity, entailing few major steps, and in the following section, we will discuss the overall recommendations (and rationale behind these).

We started this work with the hypothesis that different tasks (non-imaging variables to predict) and different datasets may tell different stories, and for that reason, we carefully chose a few different tasks (age, sex, fluid intelligence, neuroticism and smoking status) and four different datasets. We wanted to evaluate trends across different tasks and across different datasets independently. Once we had considered these independent results, we aimed to identify “one best/recommended” pipeline. The remainder of this section evolves systematically, starting from HCP and UKB datasets, and then testing on ABIDE and ACPI datasets, and lastly deducing recommendations which are reliable in all scenarios (consolidated results across all tasks and datasets). Lastly, we illustrate that the recommendations which are based on individual scenarios, and recommendations based on consolidated results are very similar.

### The best parcellation strategy

The data-driven parcellation techniques we explored for identification of brain parcels included sICA, PROFUMO, SHEN and YEO. PROFUMO yielded less predictive power than sICA identified parcels, as shown in Figure 2. This is consistent with results in [82], where PROFUMO network matrices tend to be quite empty, in part because of the fairly strong temporal independence between probabilistic functional modes. Using sICA, we investigated if the number of parcels (ICA dimensionality) affected the predictive power (illustrated in Figure 4 and briefly discussed). Our results showed that increasing dimensionality leads to an increase in predictive power. However, higher predictive power using sICA could actually be attributable to cross-subject variations in the spatial configuration of functional brain regions that are represented as changes in functional connectivity [82]. Therefore, although PROFUMO yielded lower predictive power, it still may provide a *purer* estimate, as it should not contain as much potential confounding anatomical information. sICA also outperformed SHEN and YEO parcellations in terms of predictive power as illustrated in Figure 2 (sub-figures A and C). SHEN and YEO parcellations can be affected if the reality is that of overlapping functional organisation, due to incorrect assumption of hard parcels leading to mixing of extracted time-series [83]; however, further detailed work is require to understand and address this issue. To conclude, in terms of quantitative assessment, sICA outperformed other parcellations.

### The best functional connectivity estimation method

When functional connectivity estimates are left in ambient space, without projection into tangent space, our results revealed estimation using regularised partial correlation estimation yields higher predictive power than does using full correlation/covariance and unregularised partial correlation as illustrated by the mean of the violin plots in sub-figures B and D in Figure 2. For example, for age prediction task in UKB, average correlation using covariance, full correlation, unregularised partial correlation, Tikhonov and SGGGM is 0.52, 0.52, 0.45, 0.56, and 0.55 respectively. Using partial correlation results in the removal/weakening of a considerable number of marginal and negative connections. The marginal connections were most likely due to global effects or indirect connections. Hence, partial correlation also enhanced the sparsity in the functional connectivity matrix. Partial correlation involves the inversion of covariance matrix, and to stably invert the covariance matrix, regularization should be normally applied to the covariance matrices [40]. We employed Tikhonov regularization, as it is an efficient and stable method for estimation of a regularized partial correlation matrix. We found this method resulted in increased predictive power as compared to simple partial correlation (particularly for UKB, as duration of scanning session is shorter than with HCP) as illustrated in Figure 2 (sub-figure D). However, we found a more sophisticated, recent regularization method, SGGGM, also provided reasonable predictive power (on par with Tikhonov). Through incorporation of group information when deriving intra-subject connectivity estimates, this method enables more accurately data-adaptive regularisation. This method produced the robust predictions, although optimised tuning of the parameters of SGGGM (such as sparsity level and group consistency) is essential, as the performance of SGGGM is very sensitive to these parameters. Therefore, to find optimal values for these parameters, a comprehensive grid search strategy must be applied (and, for validity, this needs to be done within cross-validation). To conclude, both in terms of qualitative and quantitative assessment, Tikhonov and SGGGM outperformed other connectivity estimation techniques.

### Is tangent space parameterization important

To test how tangent space parameterization affects predictive power, we projected different functional connectivity estimates into tangent space. As functional connectivity matrices being projected in to tangent space should always be positive definitive, estimates containing values outside of the positive definitive manifold ideally require adjustment. Functional connectivity estimates through covariance or SGGGM are always positive definitive, therefore did not require additional processing. However, partial correlation and Tikhonov connectivity matrices are not always positive definitive. To adjust for these values, we calculated the eigen-decomposition of the matrix. Any eigenvalues close to zero were adjusted to a fixed, small non-zero value. Our results demonstrated that the performance of most functional connectivity estimation techniques were reasonably improved by projecting them into tangent space, as illustrated in Figure 3 (subfigures A,C,E and G).These convincing results suggest the importance for future studies to consider the application of Riemannian geometry principles (e.g., tangent space parametrization) to functional connectivity estimates. When applied to different functional connectivity estimation methods, we found tangent space parametrization increased the predictive ability of almost all estimation techniques. Our results however failed to delineate a single connectivity estimate that was best improved by this parameterization. Projection of these estimates into tangent space required a reference mean, and so we also compared the performance of different reference means. Specifically, we compared the performance of Euclidean, harmonic, log determinant, log Euclidean and Riemannian means as reference means for tangent space parameterization. In terms of prediction accuracy/correlation, our results demonstrate the Euclidean mean tends to have the most stability (less standard deviation), as shown in Figure A.15 and A.16 (although the differences were not large between any of the reference mean methods). Therefore, we implemented tangent space parameterization using the Euclidean mean as a reference mean for proceeding analyses. The main advantage of tangent space parameterization is that it moved the functional connectivity estimates into a space where the following computations (e.g., prediction) could be done in a Euclidean framework with a corresponding boost in efficiency. To conclude, both in terms of qualitative and quantitative assessment, tangent space parameterization improves the predictive power.

### Regularization in tangent space

The functional connectivity estimates often have high estimation variability. After projecting functional connectivity estimates to tangent space, we evaluated three possible regularization strategies, (i) No shrinkage, (ii) applying the non-isotropic Population Shrinkage Covariance Estimator (PoSCE) shrinkage, (iii) applying isotropic Ledoit-Wolf shrinkage. We found that applying the PoSCE shrinkage technique as compared to no shrinkage slightly improved the performance in terms of prediction accuracy for UKB data as illustrated from top pipelines in Figure 7 and Figure A.23, but this improvement is not significant, as illustrated in Figure 3 (sub-figure H). Moreover, It did not yield meaningful improvements for HCP data, but deteriorated the performance as shown in Figure 3 (sub-figure B,D and F). We also tested the effect of tangent space projection and shrinkage on cut-down HCP data (from 60 minutes to 15 and 5 minutes), and found out that it is the tangent space projection which significantly improved the performance as illustrated in Figure 3 (sub-figures C and E), but there is no significant performance change using tangent space shrinkage as illustrated in Figure 3 (sub-figures D and F). We also tested the effect on PoSCE shrinkage by varying the number of parcels, and did not find any significant improvement as illustrated in Figure A.21.However, our analyses also revealed that optimization of the shrinkage parameter in PoSCE was not straightforward. For PoSCE, we chose the shrinkage parameter which resulted in the strongest prediction for the non-imaging variables (within an appropriate cross-validation framework). We then compared non-isotropic PoSCE shrinkage to Ledoit-Wolf isotropic shrinkage, and found that PoSCE did not yield a meaningful advantage compared with Ledoit-Wolf shrinkage (as illustrated in Figure A.19). Lastly, we compared the isotropic Ledoit-Wolf shrinkage method with the no shrinkage approach, and found that applying simple Ledoit-Wolf shrinkage improved performance for HCP data, but did not significantly improve performance for UKB data (as illustrated in Figure A.20). These results imply that simple isotropic shrinkage is more suitable for less noisy data (e.g., HCP data), and aggressive PoSCE can slightly improve performance for data with comparatively high inter- and intra-subject variance (e.g., UKB data, as duration of scanning session is shorter than with HCP). To conclude, we are recommending shrinkage in tangent space as an optional step which could be applied if data is noisy.

### Deconfounding

Many imaging studies fail to control for confounds in their analyses, which is problematic in providing replicable and interpretable results that are not attributable to extraneous variables. It is correct that in some cases the confounds may be useful to keep in the data - that is one reason why we show most results with and without confounds. However, some confounds (e.g., head motion) are generally considered to be an important factor to remove. On the other hand, age is a good example of being a factor that the experimenter may or may not want to remove - although in the context of alzheimer’s disease prediction, presumably one would include it as an additional predictor, as opposed to simply ignoring it (or deconfounding it), in order to both obtain optimal prediction and most interpretable prediction parameters. For an example, in UKB fluid intelligence prediction, one could carry out the intermediate option of deconfounding for all confounds except for head motion or age, which would give intermediate prediction correlations (in between the two extremes already presented). We have shown some of these intermediate-scenario results in Table A.5.

In our analyses, we found the optimal deconfounding strategy was X1Y1 (and X1Y0 in some cases). The reason we recommend X1Y1 over X2Y1 is because X2Y1 combination consists of performing fold-wise confound regression on the X (the functional connectivity estimates), and fold-wise deconfounding the target variable Y. However, the deconfounding strategy X1Y1 (deconfounding X once, outside of cross-validation) is more aggressive than X2Y1 (X2Y1 is less aggressive as it used cross-validation to limit the parts of the confounds actually used). For some non-imaging target variables prediction (e.g., sex) there is no chance that confound variables would feed into them (e.g., sex could not be corrupted because of head motion or age). In such cases, it is recommended to deconfound only the imaging variables, X (i.e., functional connectivity estimates) and not to deconfound the target variable, Y (i.e., sex); therefore, X1Y0 (or X2Y0) deconfounding strategy should be used for such target variables. The pretext of supporting X1Y0 over X2YO is the same (X1Y0 is more aggressive as compared to X2Y0). Our analysis have revealed that X0Y1 (deconfounding only Y), also theoretically yields plausible results; however, for some cases (e.g., sex prediction), it is not applicable. Based on our analysis, we have the following recommendations, (i) deconfound both X and Y when there is a chance that confounds can corrupt both the imaging data and non-imaging data, such as fluid intelligence score or any given mental health variable, and (ii) deconfound only X when there is no chance of confounds feeding into target variable, such as sex prediction (otherwise, this could cause Berkson’s paradox [84, 85] ^9^.

### The best classifier/predictor

Figure 5 illustrates the performance of different classifiers/predictors in combination with different parcellation and estimation techniques. Elastic Net is a simple regularized regression technique, whereas the other models tested are based on convolutional neural networks. The prediction of age, fluid intelligence and neuroticism is a continuous prediction task, whereas sex prediction is a binary classification problem. 2D RCNN consistently demonstrated the best performance for sex prediction (e.g., UKB data). 2D RCNN and Elastic Net were comparable for the prediction of fluid intelligence score (e.g., HCP data). For the prediction of neuroticism, our results demonstrated that ElasticNet provided the highest predictive power. Across the prediction of all variables, BrainNetCNN underperformed in comparison to other classifiers/predictors, although it is possible that further optimization of parameters could lead to higher accuracy. GraphCNN consistently underperformed in comparison with the other models (results presented in Figure A.31). It should be considered that we tested these deep learning architectures (2D RCNN, BrainNetCNN, and GraphCNN) for hundreds of different functional connectivity estimates, which impose practical limitations on our ability to tune hyper-parameters for each possible configuration. This indirectly suggests that only the most stable architecture will show consistent results across different connectivity estimates. Moreover, this suggests the most robust, off-the-shelf types of neural networks must minimize sensitivity to hyper-parameter tuning, to maximize generalizability and minimize overfitting. Overall, we have demonstrated that a carefully designed 2D Recurrent Convolutional Network or BrainNetCNN (top performing for fluid intelligence prediction in HCP data) can compete with (and in some cases outperform) a simpler regression model. These results contrast with [87], where kernel based regression (non-parametric classical machine learning algorithm [88]) demonstrated better performance than deep neural networks.

### Overall Results (HCP and UKB)

In the previous sections, we recommended optimal methodologies from each step of the pipeline. Our recommendation from each step is, (1) sICA for parcellation, (2) Tikhonov or SGGGM for functional connectivity estimation, (3) projecting the connectivity estimates to tangent space, (4) optional shrinkage in tangent space, (5) deconfounding the data (X1Y1 or X1Y0) and (6) Elastic Net or 2D RCNN as a predictor/classifier. It is also important to investigate these methods simultaneously (to identify potential interactions between different pipeline decisions). Figures 6, 7, A.22 and A.23 illustrate the top performing pipelines out of hundreds of combinations. The top performing methods which repeatedly stand out are in-fact the joint combination of these methodologies; i.e., “ICA SGGGM Tangent Elastic Net” or “ICA Tikhonov Tangent shrinkage 2DRCNN”, etc. It is to be noted that our recommendations are not solely based on quantitative results but also on the qualitative assessment of each step as explained through-out the Discussions (e.g., one has more parcels than time points, Tikhonov or SGGGM may be able to give stable results). The error bars drawn (using Fisher Transformation/Wilson test) in Figures 6, 7,A.22 and A.23 show that for sex and age prediction, these recommended pipelines are significantly different from the other non-recommended pipelines. To make sure that all non-imaging variables prediction are reproducible and significant, we ran repeated cross-validation on the top performing pipelines. With a 99% confidence interval, the fluid intelligence scores prediction correlation is ∼0.35 ± 0.006, for age it is ∼0.58 ± 0.005 and for sex it is ∼0.92 ± 0.001. This highlights that the identified optimum pipelines are significantly different and reproducible.

### Results External Evaluation (ABIDE and ACPI)

To test the generalisability of our recommendations, we ran our methodological techniques on ABIDE and ACPI datasets. We first compared our recommended pipelines with randomly chosen non-recommended pipelines, and results are presented in Table 3. It is to be noted that these datasets have fewer subjects and are comparatively more noisy as compared to HCP and UKB. As shown in Table 3, recommended pipelines outperformed the other pipelines significantly (particularly for participants’ age prediction task on ABIDE). This supports that the recommended steps are generalizable. Moreover, Figures A.33 and A.29 show the full set of results for age and smoking status prediction for ABIDE and ACPI data respectively. These results support our recommendations in general, although the performance of CNNs (both 2D RCNN and BrainNetCNN) exceeded our expectations. The results reported on CNNs are without hyper-tuning (same fixed parameters as used in HCP/UKB). This could possibly be explained by CNNs being more powerful on noisier data (relatively speaking), may be similar in effect to data augmentation [89]. We have also tested our methodological choices with some other commonly used techniques. We compared the performance of the CPM predictive model with Elastic Net and found that Elastic Net performs better (as shown in Figure A.30). We also compared Elastic Net with Random Forests on a subset of results, and found that Elastic Net performs better (as shown in Figure A.32).

### Combined Results across all tasks and datasets

In this paper, we followed a step-wise approach and have presented our results systematically, starting from HCP and UKB datasets, and then testing on ABIDE and ACPI datasets. Lastly, we condensed the results across all four datasets (UKB, HCP, ABIDE and ACPI) and all tasks (non-imaging variables prediction), and have shown the consolidated results in Figure 11. The relative prediction scores shown in Figure 11 are defined as the prediction score of each pipeline relative to the average across all pipelines. The recommendations which we have found on individual datasets are similar to our conclusions deduced from consolidated results: (1) the mean of relative prediction scores using sICA is 0.07, compared to YEO (0.04), SHEN (0.03) and PROFUMO (−0.03), (2) the mean of relative prediction scores using Tikhonov and SGGGM is 0.02, compared to covariance and Full correlation (0.0), and partial correlation (−0.02), (3) the mean of relative prediction scores using tangent space is 0.05, compared to ambient space (−0.02) and tangent sharinkage space (−0.04), (4) the mean of relative prediction scores after deconfounding is -0.05, which is negative, but we believe the results without applying deconfounding are inflated due to the shared confounds, and lastly (5) the mean of prediction score using Elastic Net and 2D RCNN is 0.11 and 0.04 respectively, compared to BrainNetCNN (0.0), and GraphCNN (−0.16). To conclude, there is an approximately “optimal” pipeline (or set of recommendations) which performed reasonably well across all tasks whether analysed individually or in a consolidated fashion.

### Dimensionality of Parcellation Revisited

Finally, it is important to consider the impact of the number of parcels across various parcellation methods, although it is hard to fully disentangle different choices of dimensionality from different parcellation methods. We have nevertheless attempted to draw recommendations by comparing YEO parcellation with varying the number of parcels (d = 100, 200, 400) with sICA (d = 15, 50, 200) as shown in Figure 11 (sub-figure D). We found that higher dimensional sICA (d=200) outperforms lower dimensional sICA (d=50), which is also illustrated in Figure 4, and briefly discussed above the “best parcellation strategy” section. Moreover, lower dimensional sICA (d=50) outperforms both lower and higher dimensional YEO parcellation (d = 100 or 400). Lastly, we found that prediction performance is not always improved by increasing the dimensionality of parcellation method, as better results were obtained by lower dimensional YEO (d=100) compared to higher dimensional YEO (d = 200 or 400). This was also the main rationale behind showing lower dimensional YEO (d=100) results throughout the main text. It is also important to comment on the dimensionality of sICA for UKB data, as in theory higher ICA dimensionality might have been useful for UKB data, but we can not explore high dimensionality with ICA as the number of non-artefact (neuronal signal) components barely grows at all (more than the existing 55). As discussed above, we believe this is because of the limitations of volumetric analysis, possibly combined with the 6-minute limited duration of the acquisition in UKB (as we do not face such limitations in case of HCP).

### Comparison with Literature

Somewhat similar evaluations of various methods (in this problem domain) have previously been carried out in [90, 91]. In [90], authors proposed that matrix whitening transform and parallel transport could be utilized to project covariance matrices into a common tangent space and evaluated this method on twenty four healthy subjects. These former results build a foundation for estimation of functional connectivity using manifold operations, as higher prediction accuracy was achieved using tangent space parameterization.

[91] evaluated different modelling choices and recommended a three stage pipeline: (i) ICA or MODL for brain parcellation, (ii) tangent space embedding for functional connectivity estimation and (iii) use of non-sparse models like SVM for non-imaging variables prediction. For brain parcellation, [91] applied a number of parcellation techniques that differed from our pipeline, however also found that sICA yielded the best performance (alongside with MODL). In terms of step-1 (brain parcellation), we compared sICA with MODL, and found that the performance of these is similar (although sICA outperforms for age prediction, as shown in Figure A.33). We have not included MODL in main analysis as it is already thoroughly evaluated in [91] (and it remains a valuable tool for brain parcellation).

Lastly, we have applied deep learning architecture such as CNNs to analyze functional connectivity whereas [91] utilized traditional machine learning algorithms. Similarly to the last step of our pipeline, [87] compared the performance of three CNNs to kernel regression. The three CNNs tested included a generic fully-connected feedforward network, BrainNetCNN and Graph CNN. This study reported that CNNs did not outperform kernel regression across a wide range of behavioral and demographic measures. However, our results found that our proposed 2D RCNN (architecture illustrated in Figure A.12), which has not been previously evaluated against these methods, outperforms the regularized regression and existing CNNs for some tasks (e.g., Figure A.25).

## 6. Conclusion

Our results have demonstrated that within data driven parcellation, ICA provides the most predictive power for non-imaging variables. For estimating functional connectivity, we have shown that regularized partial correlation like SGGGM or Tikhonov outperforms unregularized partial correlation or full correlation. To correctly apply mathematical formulations on functional connectivity estimates, it is recommended to take into account that these measures should ideally be worked with within the Riemannian manifold. To do so, Riemannian space is approximated with an associated tangent space and then functional connectivity estimates are mapped to the tangent space. We have reported in this paper that this tangent space parameterization results in a significant increase in predictive power of functional connectivity estimates, as previously shown in [90, 91]. Our results have also demonstrated that additional shrinkage is not necessarily required on well regularized connectivity estimates in tangent space; however, could be applied for noiser data. Lastly, we evaluated various classifiers for prediction of non-imaging variables from connectivity estimates and concluded that a carefully designed deep learning based architecture (2D RCNN) can be a valuable tool for analyzing functional connectivity. However, Elastic Net probably performs better at present overall.

## 7. Acknowledgements

Computation used the Oxford Biomedical Research Computing (BMRC) facility, a joint development between the Wellcome Centre for Human Genetics and the Big Data Institute supported by Health Data Research UK and the NIHR Oxford Biomedical Research Centre.

We are extremely grateful to all UK Biobank (Data Access Application 8107), Human Connectome Project, ACPI, and ABIDE study participants, who generously donated their time to make this resource possible. UK Biobank (including the imaging enhancement) has been generously supported by the UK Medical Research Council and the Wellcome Trust. Human Connectome Project, WU-Minn Consortium (Principal Investigators: David Van Essen and Kamil Ugurbil; 1U54MH091657) funded by the 16 NIH Institutes and Centers that support the NIH Blueprint for Neuroscience Research; and by the McDonnell Center for Systems Neuroscience at Washington University. The ACPI is primarily supported by a grant supplement (R01 MH094639) provided to PI Milham by the National Institute on Drug Abuse (NIDA). Additional support to ACPI is provided by the Child Mind Institute and the Nathan Kline Institute. For ABIDE, the primary support for the work by Adriana Di Martino was provided by the (NIMH K23MH087770) and the Leon Levy Foundation and primary support for the work by Michael P. Milham and the INDI team was provided by gifts from Joseph P. Healy and the Stavros Niarchos Foundation to the Child Mind Institute, as well as by an NIMH award to MPM (NIMH R03MH096321).

We additionally thanks Gael Varoquaux, Giles Colclough, Mehdi Rahim, Thomas E. Nichols, and Todd Constable for helpful discussions.

Lastly, U.P is funded by an MRC Mental Health Data Pathfinder award (PI Clare Mackay) MC PC 17215. S.M.S and M.W.W receive support from the Wellcome Trust.

## Appendix A. Inline Supplementary Figures

**Table A.4:**
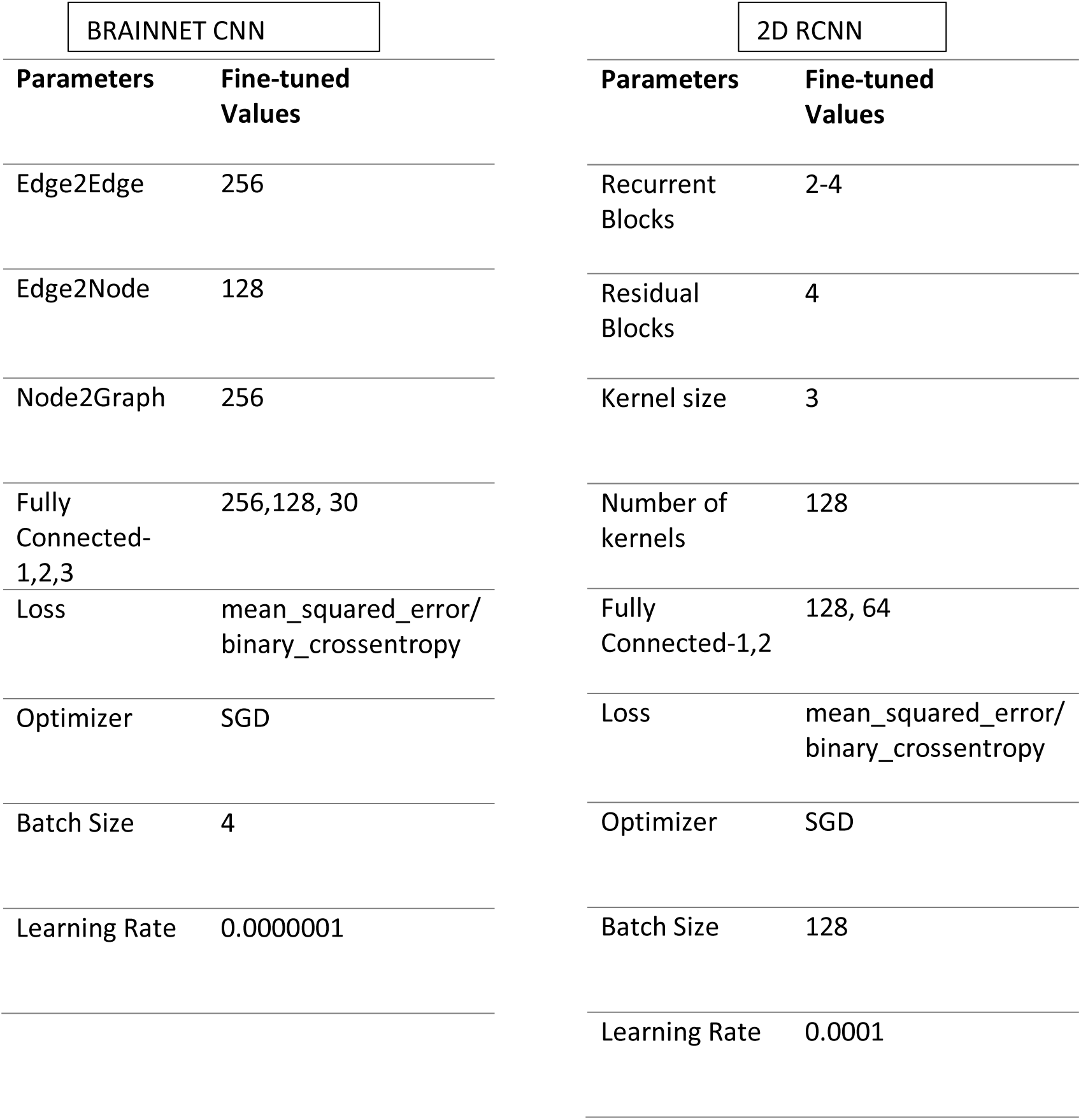
The tuned hyper-parameters for BrainNetCNN and 2DRCNN.

**Table A.5:** The impact of deconfounding each confound separately on the prediction of fluid intelligence in the HCP data.

| **Method** | **r value** |
| --- | --- |
| No Deconfounding | 0.2934 |
| Deconfounding Age | 0.2815 |
| Deconfounding Age + Sex | 0.2720 |
| Deconfounding Sex | 0.2714 |
| Deconfounding ethnicity | 0.2919 |
| Deconfounding Height | 0.2739 |
| **Deconfounding rfMRI motion** | 0.2552 |
| **Deconfounding FS IntraCranial Volume** | 0.2240 |
| **Deconfounding FS BrainSeg Volume** | 0.2378 |
| Deconfounding all variables | 0.1926 |

**Figure A.12:**
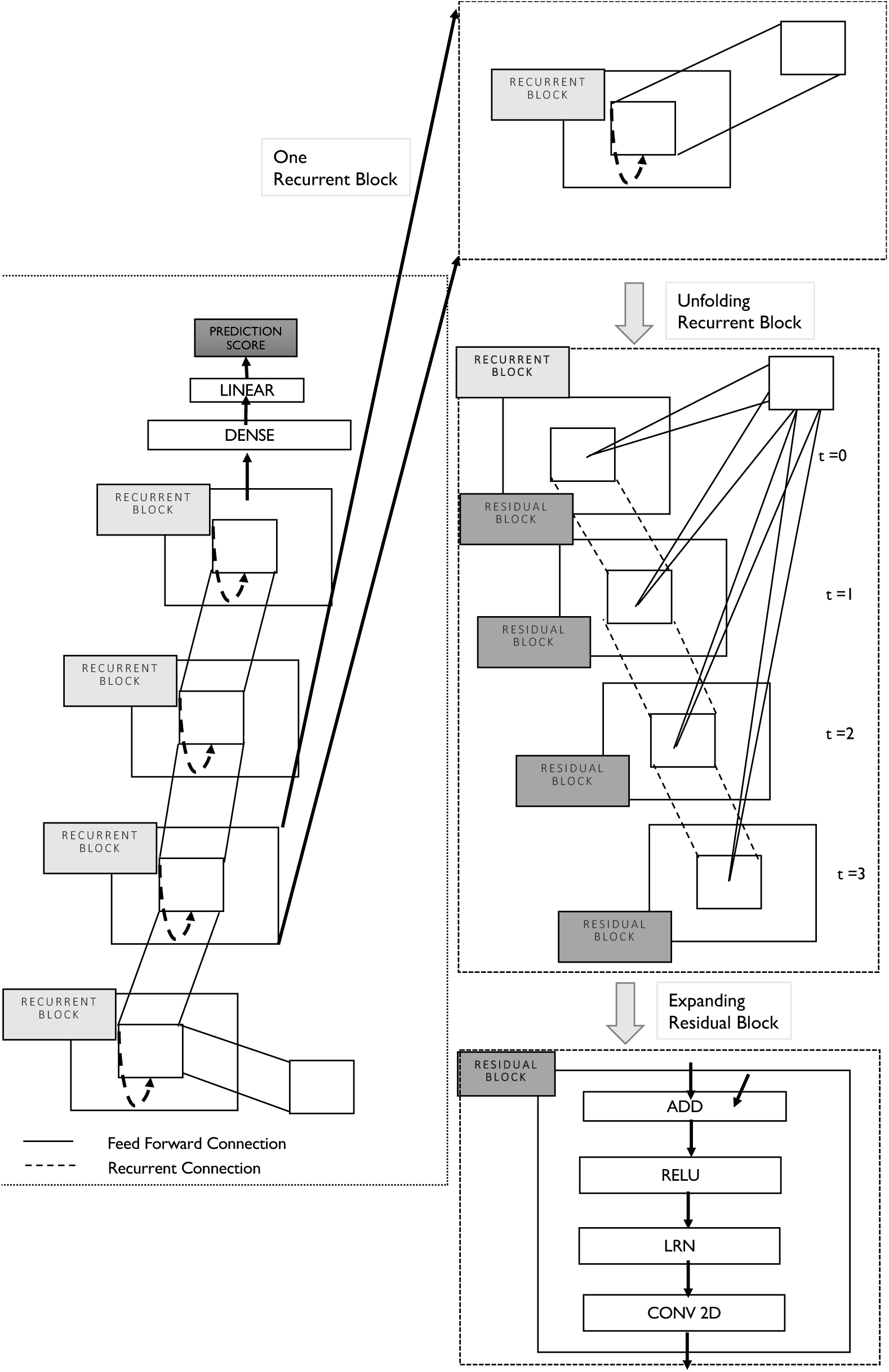
The proposed architecture of 2D RCNN. A recurrent block is unfolded on the right for t = 3 time steps. At t = 0, it is only a feed-forward network with a single residual block. At t =3, it has depth of 4 with additional recurrent connections. Each residual block is further composed of addition, batch normalization, convolution and activation layers.

**Figure A.13:**
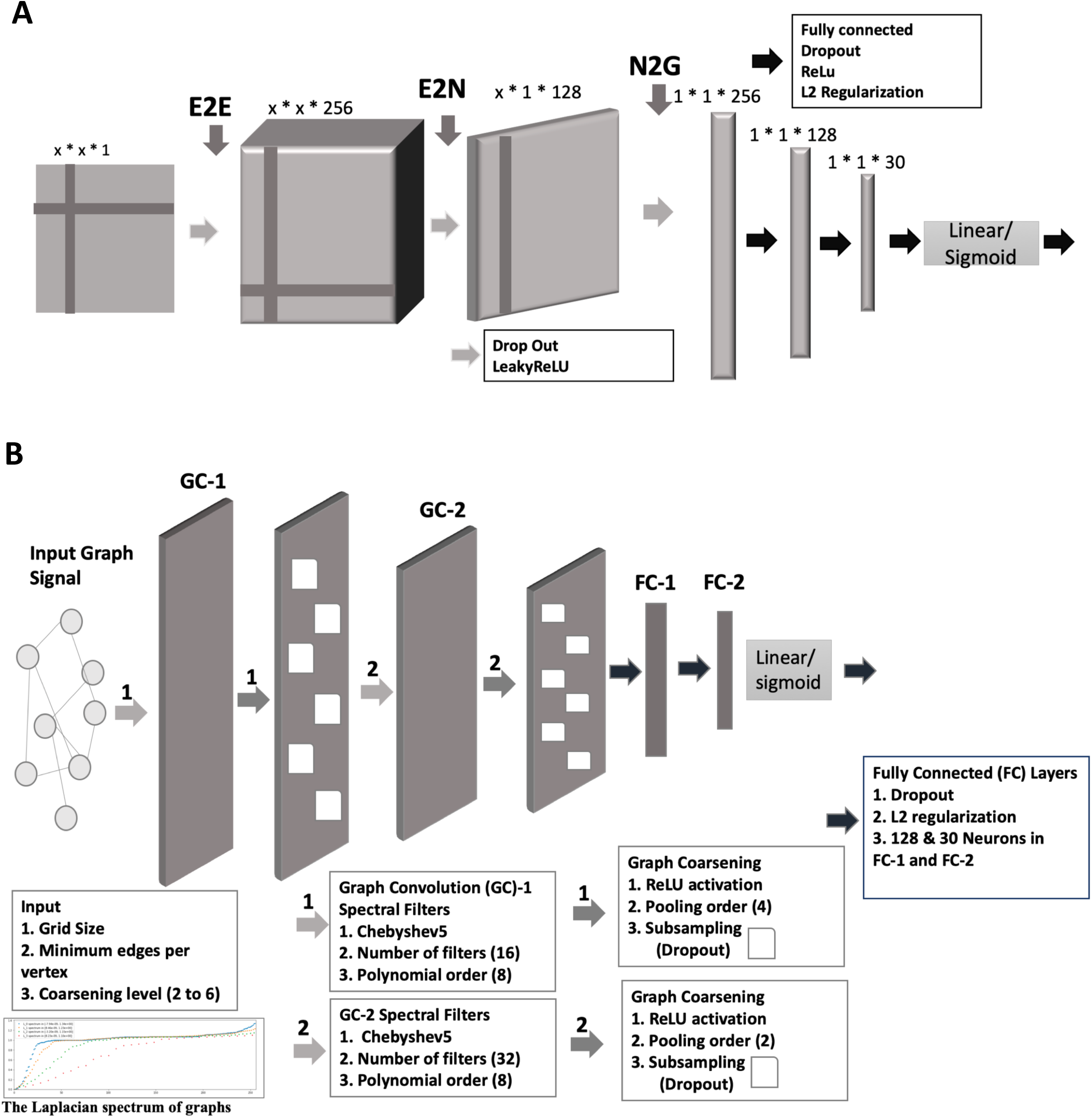
The architectures of DNNs: **(A)** The BrainNetCNN architecture, where the E2E layer considers weights of all neighbouring edges (adjacent brain regions), the E2N layer convolves netmats with 1D convolutional filter producing a single output for each node and finally the N2G layer reduces dimensionality. **(B)** the schematic of GraphCNN, which can be summarized as 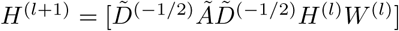, where A is the adjacency matrix, W is weight matrix and D is degree matrix. Input graph signals pass through a set of convolution, pooling and fully connected layers resulting in producing a single output score corresponding to non-imaging variable for each subject’s functional connectivity matrix.

**Figure A.14:**
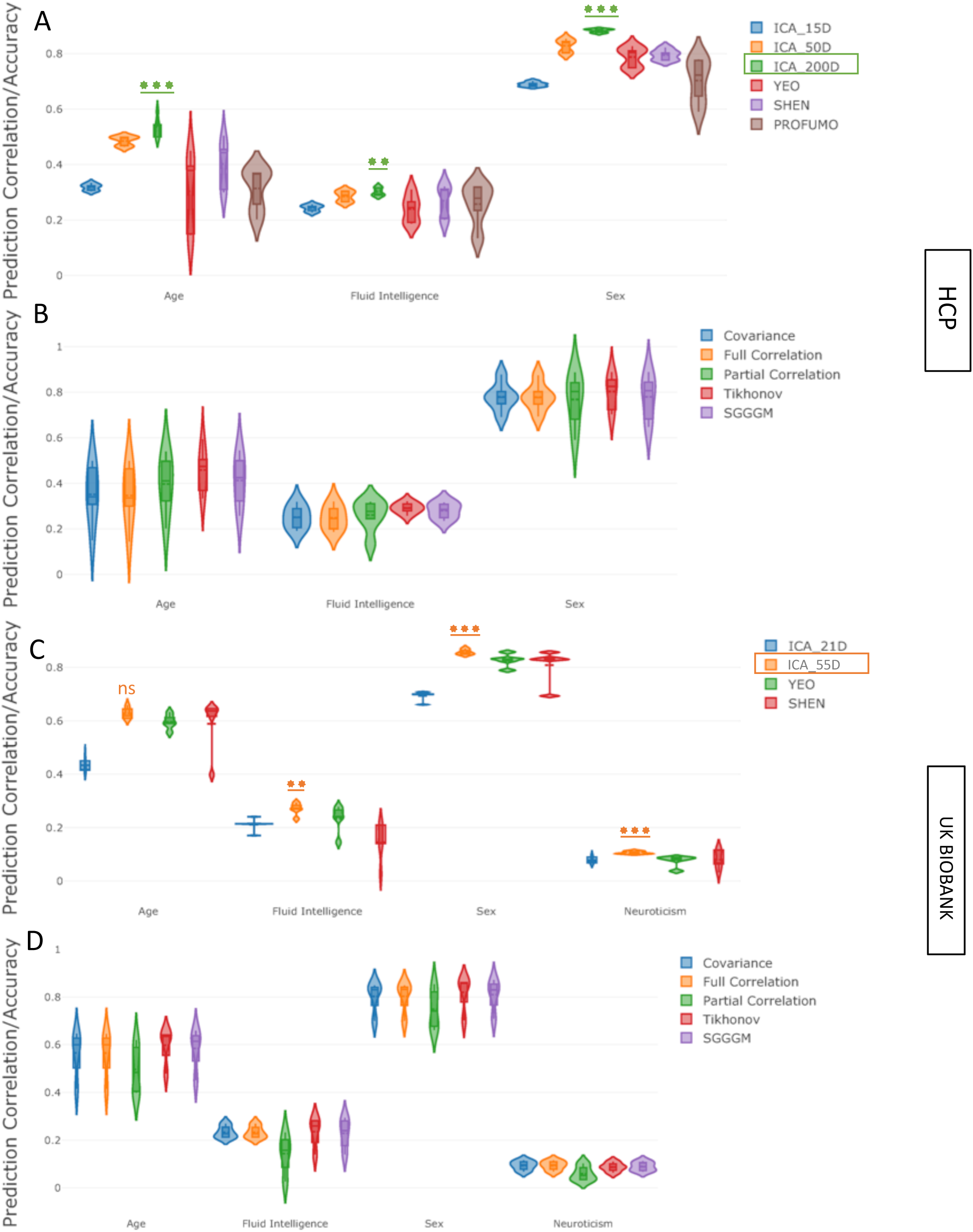
The impact of various parcellation strategies and functional connectivity estimation methods on prediction power for non-imaging variables **without deconfounding. [A**,**B] (HCP Data), [C**,**D] (UKB Data):** The violin plots in **A** and **C** show the prediction variability over 5 measures of functional connectivity estimates and in **B** and **D** show the prediction variability the over different parcellation schemes. For HCP, the ICA based parcellation schemes are *ICA*_15*D, ICA*_50*D* and *ICA*_200*D*, and for UKB are *ICA*_21*D* and *ICA*_55*D*, where D = the number of parcels. For both HCP and UKB, SHEN parcellation was 268D, YEO was 100D, and PROFUMO was 50D (for HCP only). The stars refer to comparison against the next-best method.

**Figure A.15:**
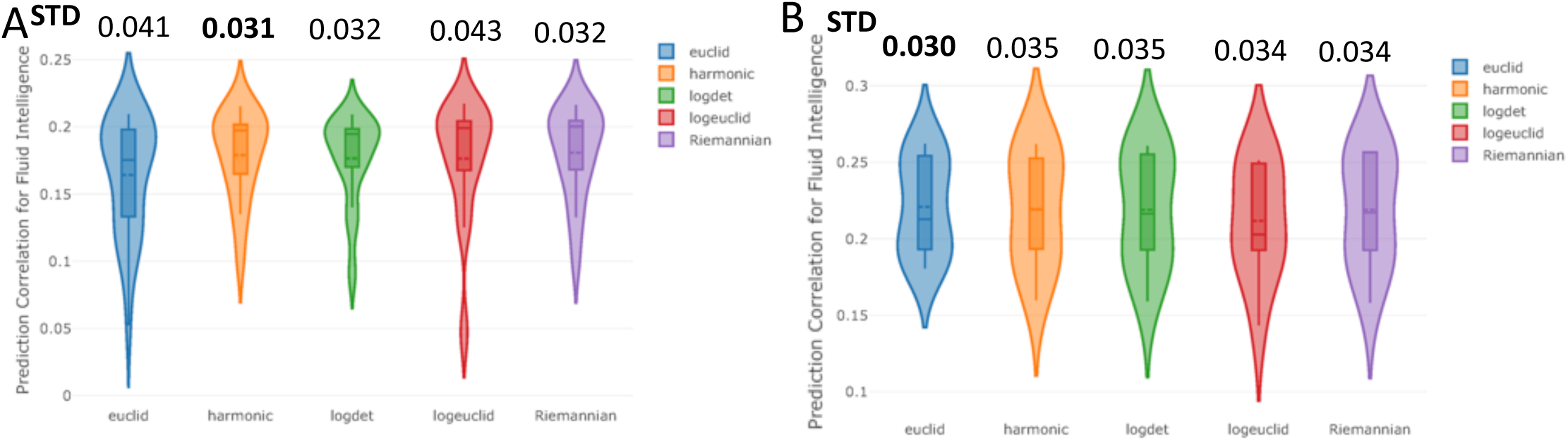
The impact of various reference mean estimation techniques during tangent space parameterization on prediction of non-imaging variables **after deconfounding. [A] (HCP Data), [B] (UKB Data)** These figures show the prediction correlation for fluid intelligence scores when functional connectivity estimates are projected into tangent space, using different reference means. The violin plots represent the prediction variability over 4 different parcellation schemes and 5 measures of functional connectivity estimates in the tangent space.

**Figure A.16:**
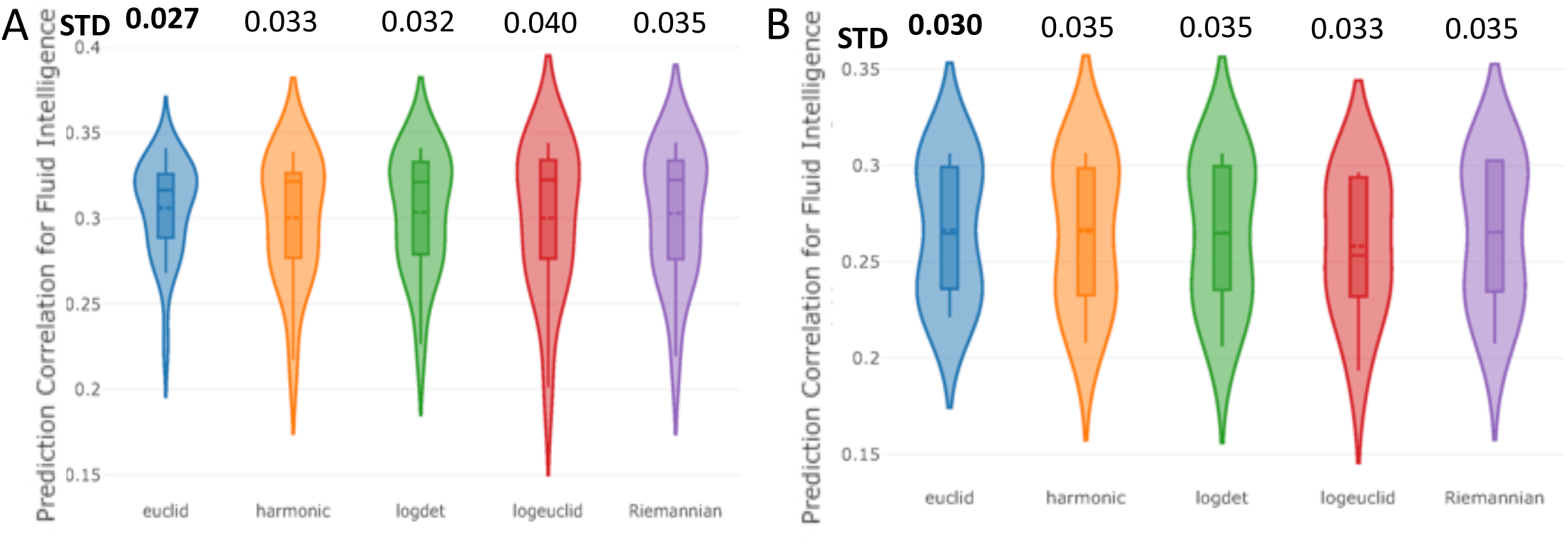
The impact of various reference mean estimation techniques during tangent space parameterization on prediction of non-imaging variables **without deconfounding. [A] (HCP Data), [B] (UKB Data)** These figures show the prediction correlation for fluid intelligence scores when functional connectivity estimates are projected into tangent space, using different reference means. The violin plot represents the prediction variability over 3 different parcellation schemes and 5 measures of functional connectivity estimates in the tangent space.

**Figure A.17:**
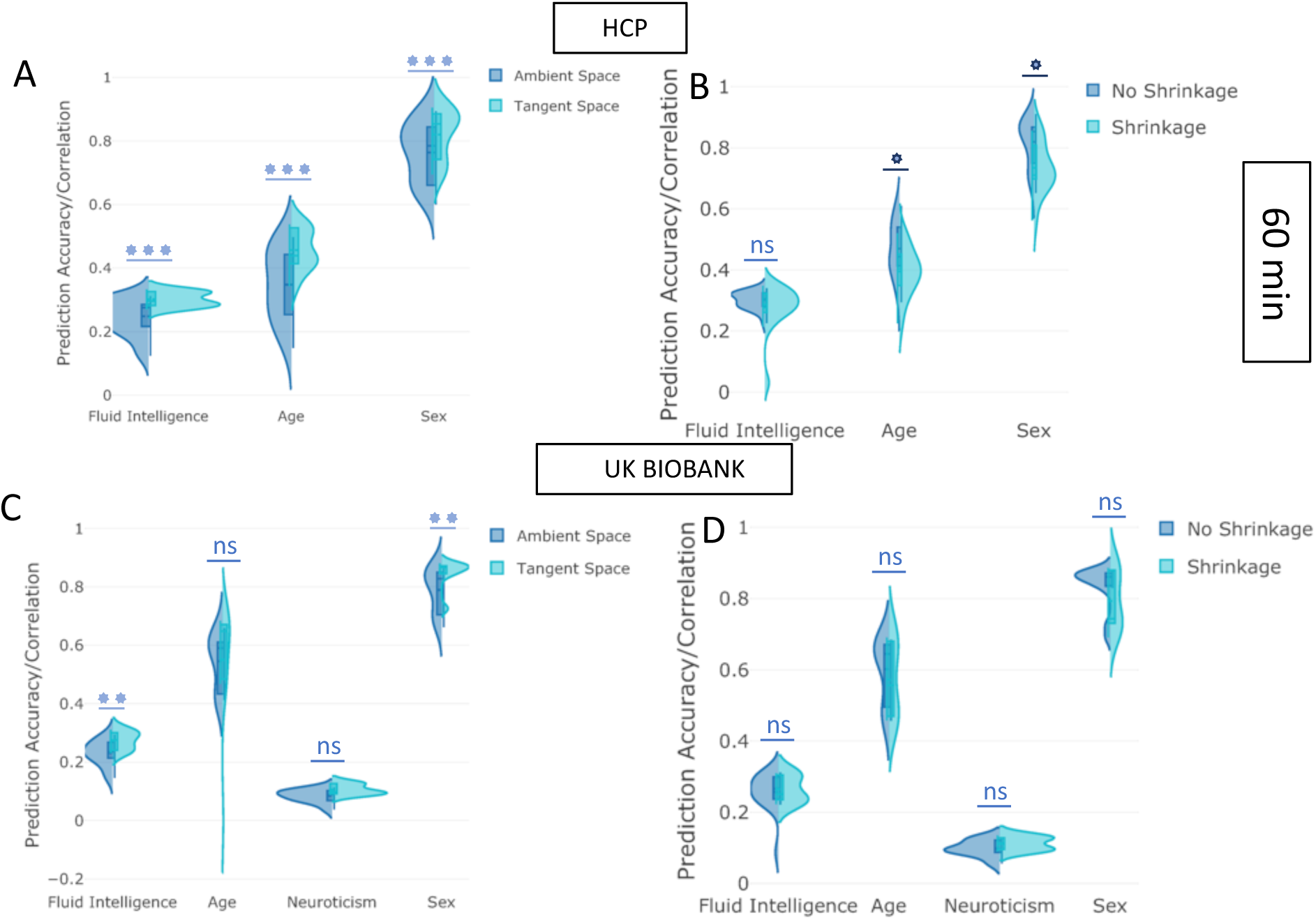
The impact of projecting to tangent space and applying shrinkage in tangent space on prediction power **before deconfounding. [A-B] (HCP Data), [C**,**D] (UKB):** The y-axis depicts the prediction accuracy/correlation for different behavioural measures. “Tangent Space” means that tangent space projection was applied on functional connectivity estimates (originally in the “Ambient Space”). The “Shrinkage” strategy means that non-isotropic PoSCE shrinkage was applied to connectivity estimates in tangent space before feeding to the predictor/classifier. “No Shrinkage” means that projected functional connectivity estimates in tangent space were directly fed to the predictor/classifier, and did not undergo PoSCE shrinkage. The violin plots show the prediction variability over 4 different parcellation schemes and 5 measures of functional connectivity estimates.

**Figure A.18:**
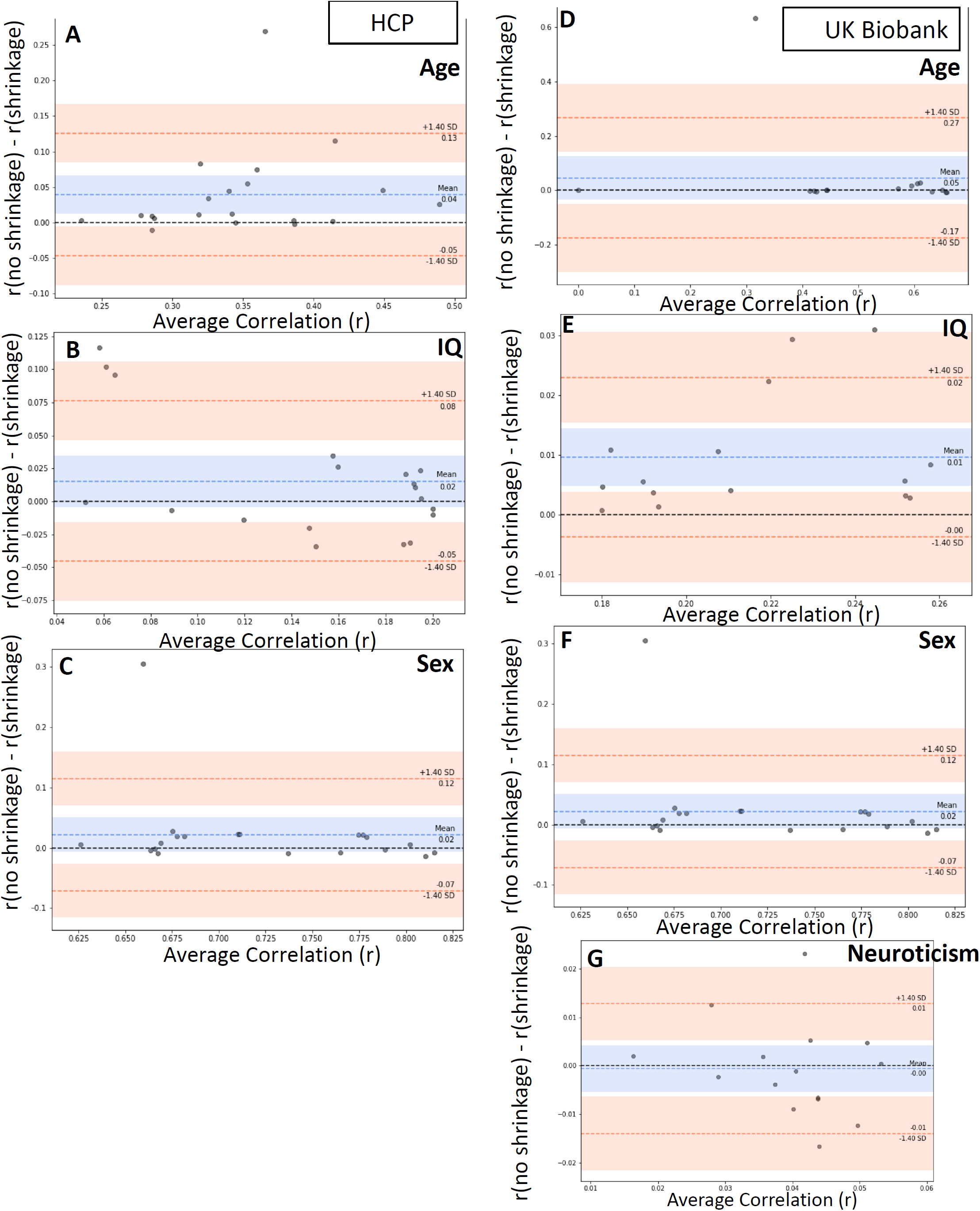
The impact of applying shrinkage in tangent space on prediction power **after deconfounding. [A**,**B**,**C] (HCP Data), [D**,**E**,**F**,**G] (UKB Data):** The y-axis depicts the difference of two methods (no shrinkage -shrinkage) and x-axis depicts the mean of methods ((no shrinkage + shrinkage) /2).

**Figure A.19:**
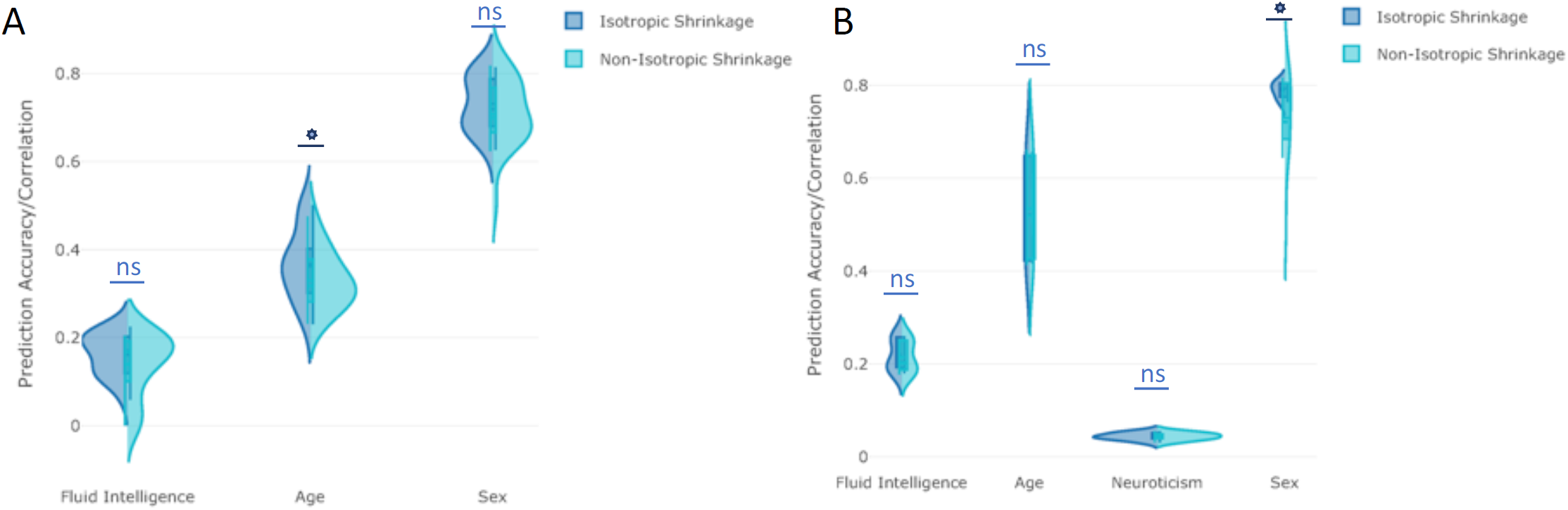
The impact of isotropic versus non-isotropic shrinkage in tangent space on prediction power for non-imaging variables **after deconfounding. [A] (HCP Data), [B](UKB)**: The y-axis depicts the prediction accuracy/correlation for different non-imaging measures. Isotropic Shrinkage means that Ledoit-Wolf shrinkage was applied to projected functional connectivity estimates in tangent space before they were fed to a predictor/classifier. The Non-Isotropic Shrinkage strategy means that PoSCE shrinkage was applied to connectivity estimates in tangent space before they were fed to a predictor/classifier.

**Figure A.20:**
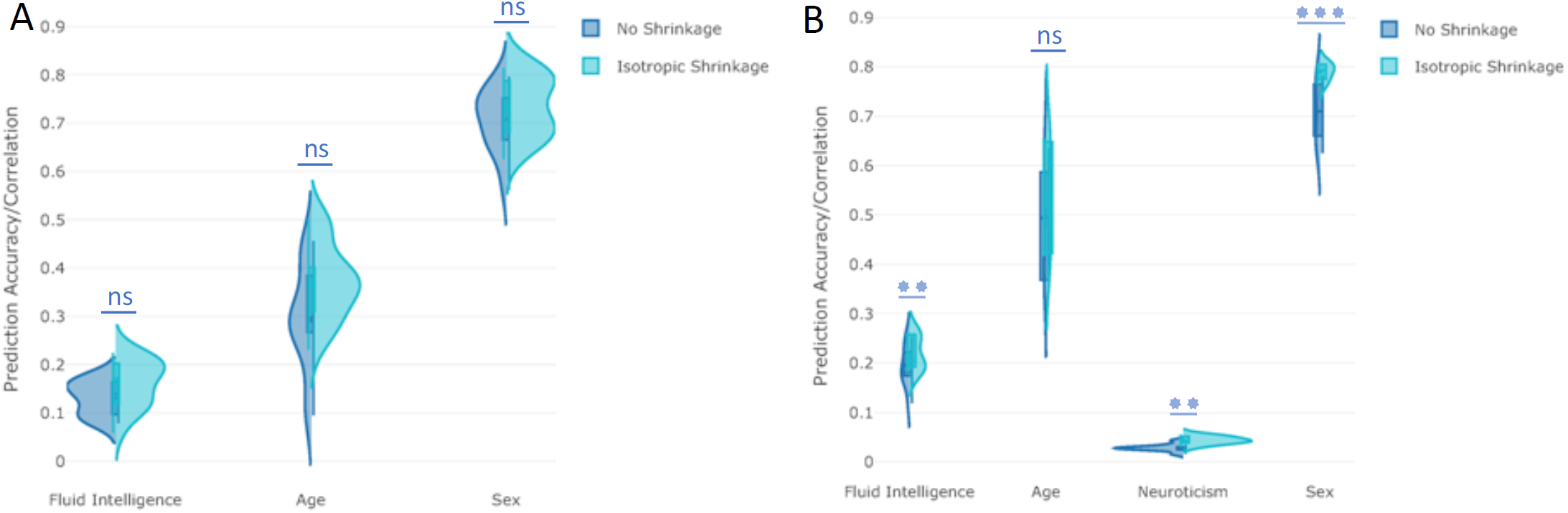
The impact of applying isotropic shrinkage in tangent space on prediction power after deconfounding. **[A] (HCP Data), [B](UKB)**: The y-axis depicts the prediction accuracy/correlation for different non-imaging measures. Isotropic Shrinkage means that Ledoit-Wolf shrinkage was applied to projected functional connectivity estimates in tangent space before they were fed to a predictor/classifier. “No Shrinkage” means that projected functional connectivity estimates in tangent space were directly fed to the predictor/classifier, and did not undergo Ledoit-Wolf shrinkage.

**Figure A.21:**
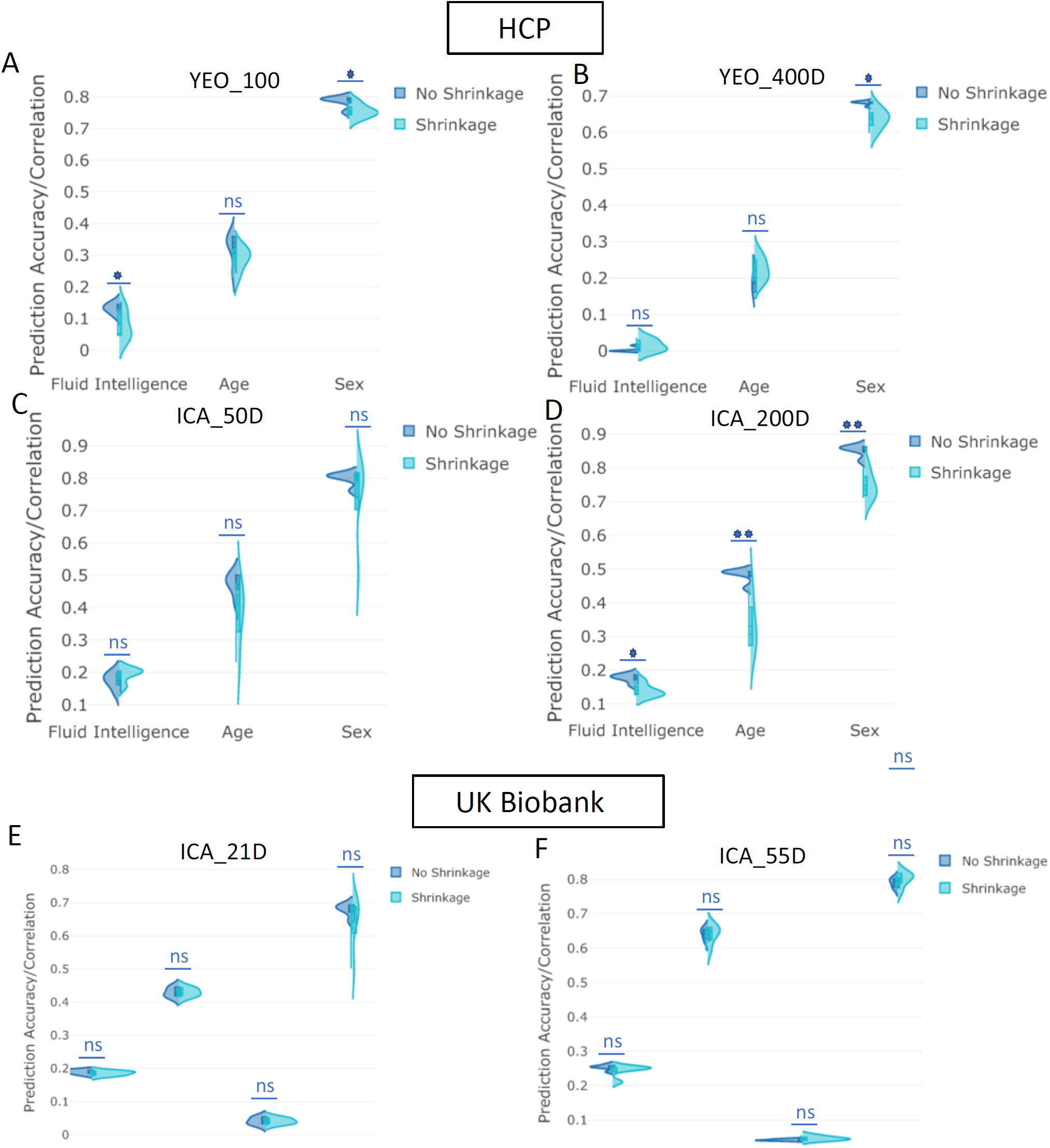
The impact of applying shrinkage in tangent space on prediction power varying the number of parcels. **[A-D] (HCP Data), [E**,**F](UKB)**: The y-axis depicts the prediction accuracy/correlation for different behavioural measures. The “Shrinkage” strategy means that non-isotropic PoSCE shrinkage was applied to connectivity estimates in tangent space before feeding to the predictor/classifier. “No Shrinkage” means that projected functional connectivity estimates in tangent space were directly fed to the predictor/classifier, and did not undergo PoSCE shrinkage. The “YEO_400D” means that 400D YEO parcellation was applied. The violin plots show the prediction variability over 5 measures of functional connectivity estimates for the mentioned parcellation scheme (e.g., “YEO_400D”)

**Figure A.22:**
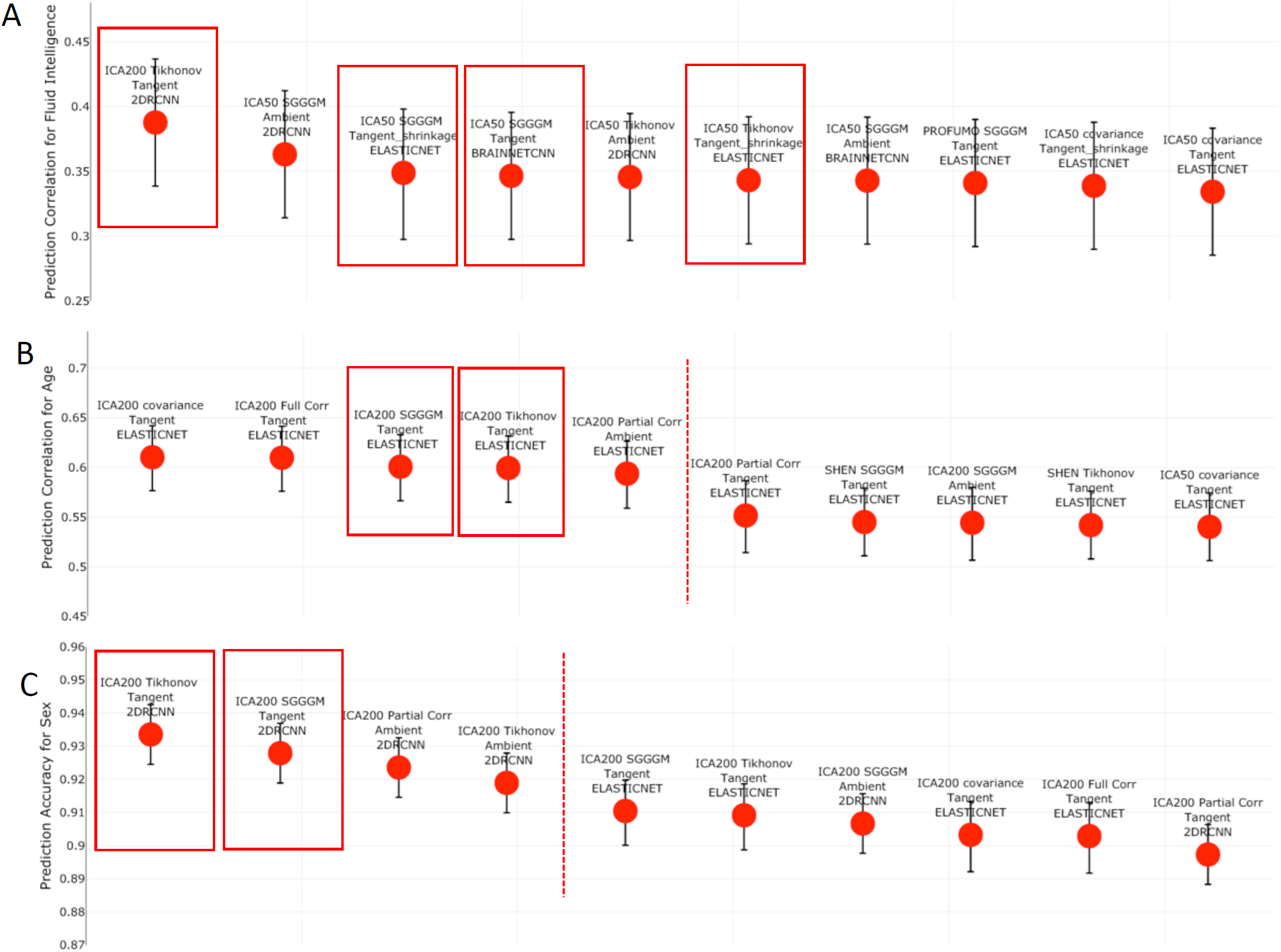
The top performing ten configurations for the prediction of each non-imaging variable by dataset **without deconfounding. [A**,**B**,**C] (HCP Data)** Each data point represents a different configuration strategy that may vary in terms of parcellation strategy, the functional connectivity estimation method, whether tangent space parameterization was employed, whether tangent space regularization was employed, and the predictor/classifier that was used. The first word indicates the parcellation strategy, and the second word refers to the functional connectivity estimation method. The third word refers to the geometry in which classifier is applied, ambient referring to non-tangent space and tangent referring to the projected covariance matrices in tangent space. If non-isotropic shrinkage was applied after projecting covariance matrices to tangent space, the fourth word will be “shrinkage”. The last word indicates the type of classifier/predictor that was used. The highlighted red blocks show the recommended pipelines (rationale explained in Section 5), and red dotted lines highlight the point when the error bar of pipeline after the dotted line is out of range from the error bar of the top (first) pipeline.

**Figure A.23:**
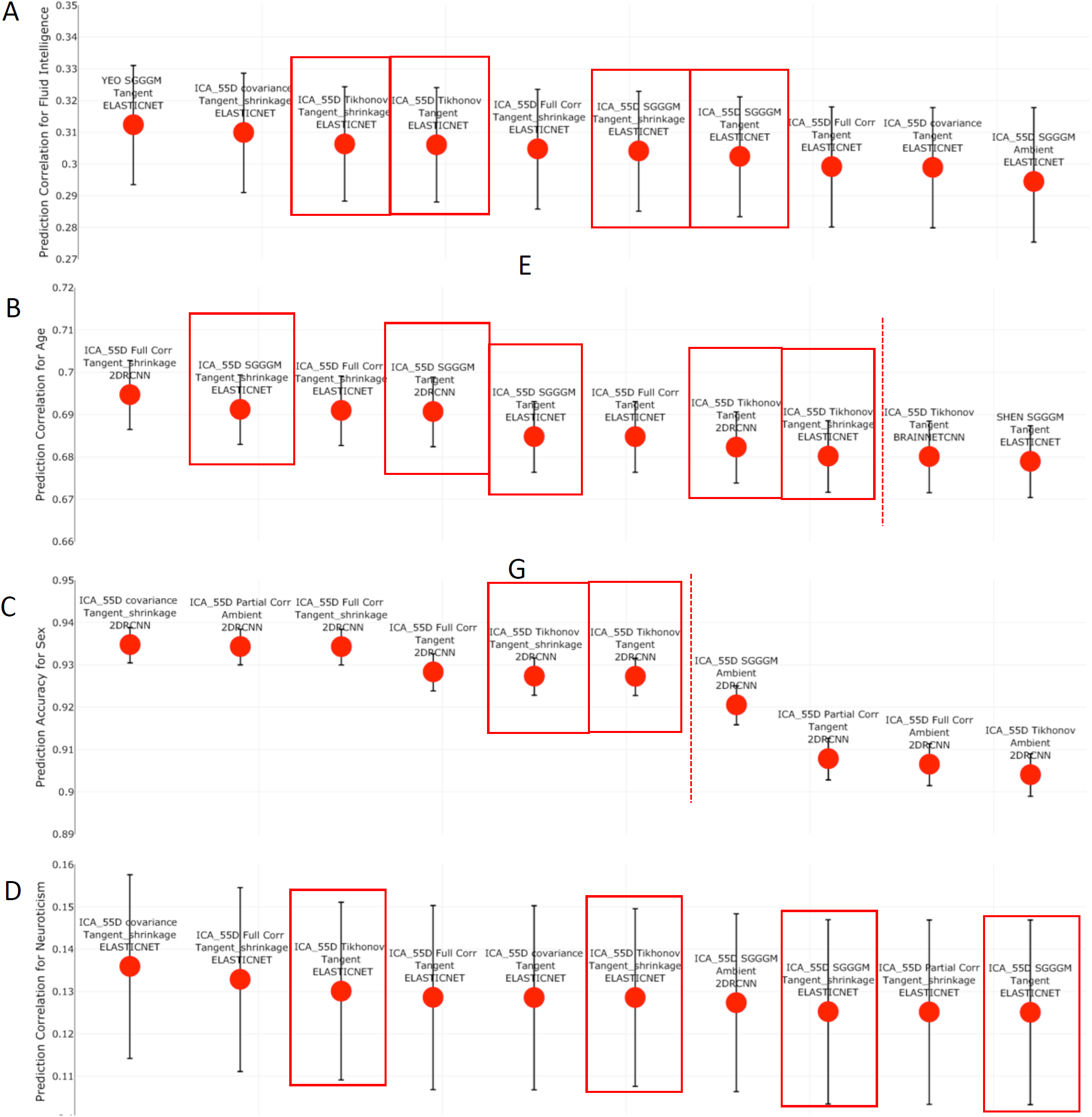
The top performing ten configurations for the prediction of each non-imaging variable by dataset **without deconfounding. [A**,**B**,**C**,**D] (UKB Data)** Each data point represents a different configuration strategy that may vary in terms of parcellation strategy, the functional connectivity estimation method, whether tangent space parameterization was employed, whether tangent space regularization was employed, and the predictor/classifier that was used. The first word indicates the parcellation strategy, and the second word refers to the functional connectivity estimation method. The third word refers to the geometry in which classifier/predictor is applied, ambient referring to non-tangent space and tangent referring to the projected covariance matrices in tangent space. If non-isotropic shrinkage was applied after projecting covariance matrices to tangent space, the fourth word will be “shrinkage”. The last word indicates the type of classifier/predictor that was used. The highlighted red blocks show the recommended pipelines (rationale explained in Section 5), and red dotted lines highlight the point when the error bar of pipeline after the dotted line is out of range from the error bar of the top (first) pipeline.

**Figure A.24:**
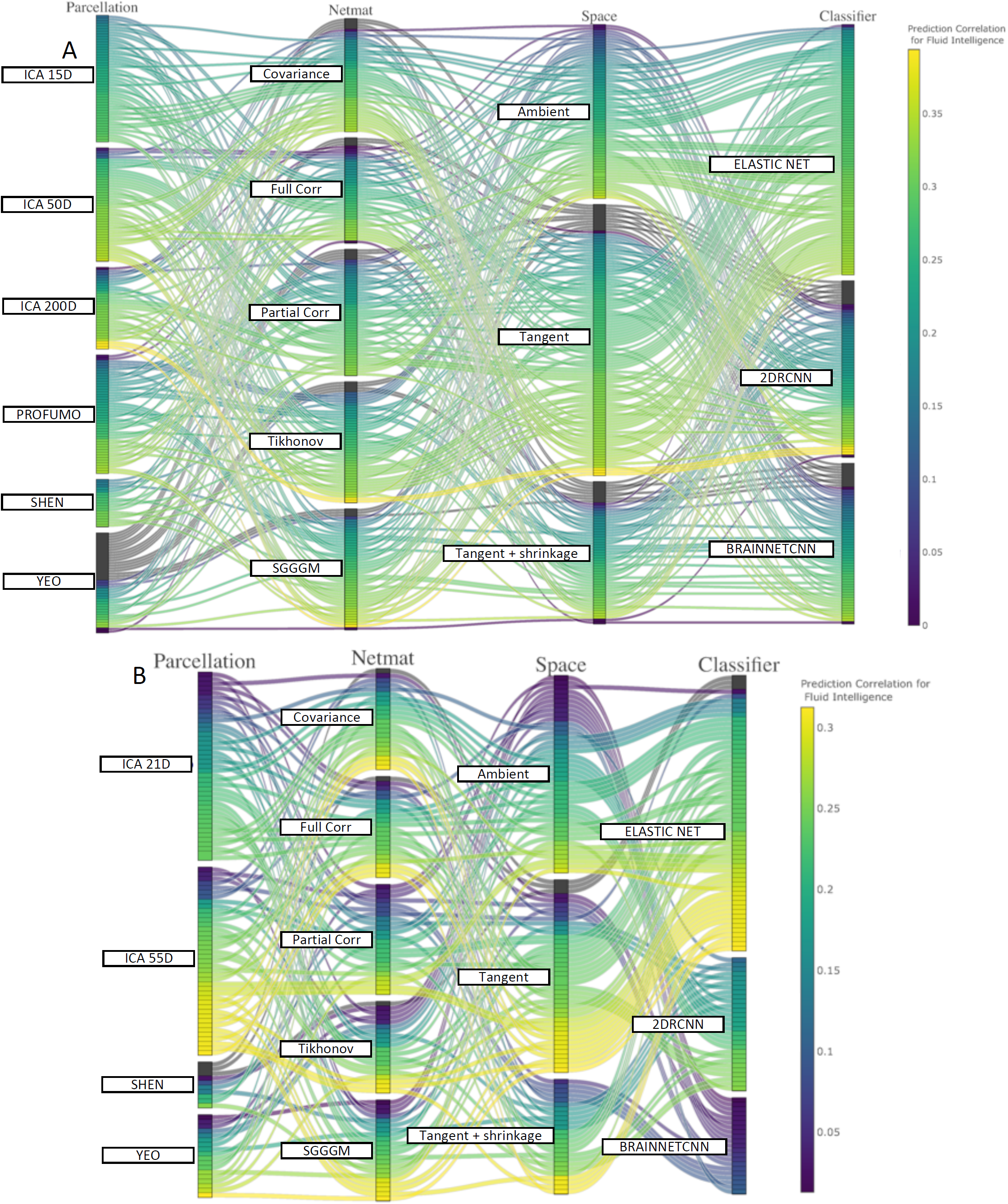
This parallel coordinates plot provides a visualization of all possible combinations of options in the pipeline to predict fluid intelligence scores from functional connectivity **without deconfounding. [A] (HCP Data), [B] (UKB Data):** The lines are color-coded according to their prediction performance.

**Figure A.25:**
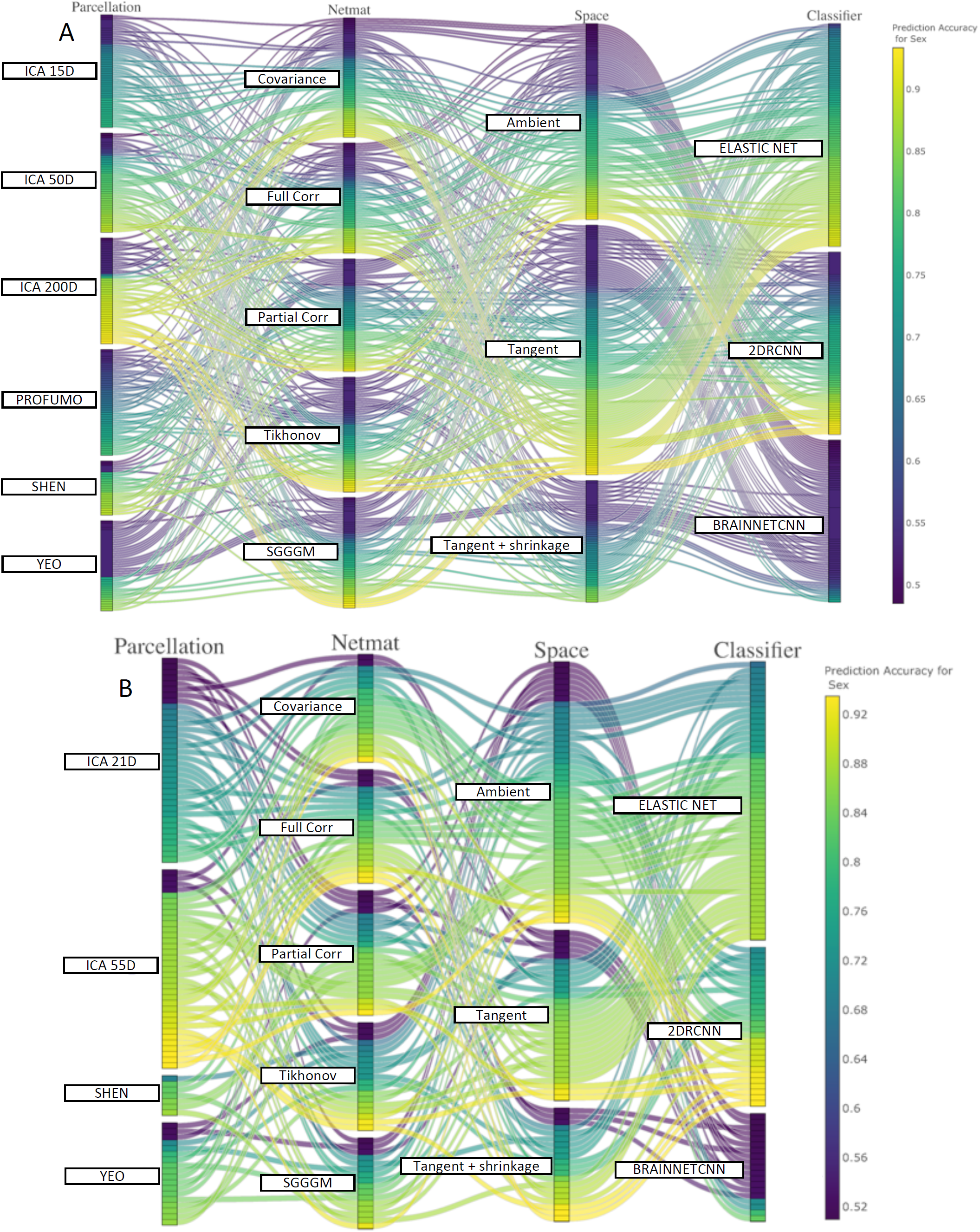
This parallel coordinates plot provides a visualization of all possible combinations of options in the pipeline to pipeline to predict sex from functional connectivity **without deconfounding. [A] (HCP Data), [B] (UKB Data):** The lines are color-coded according to their prediction performance.

**Figure A.26:**
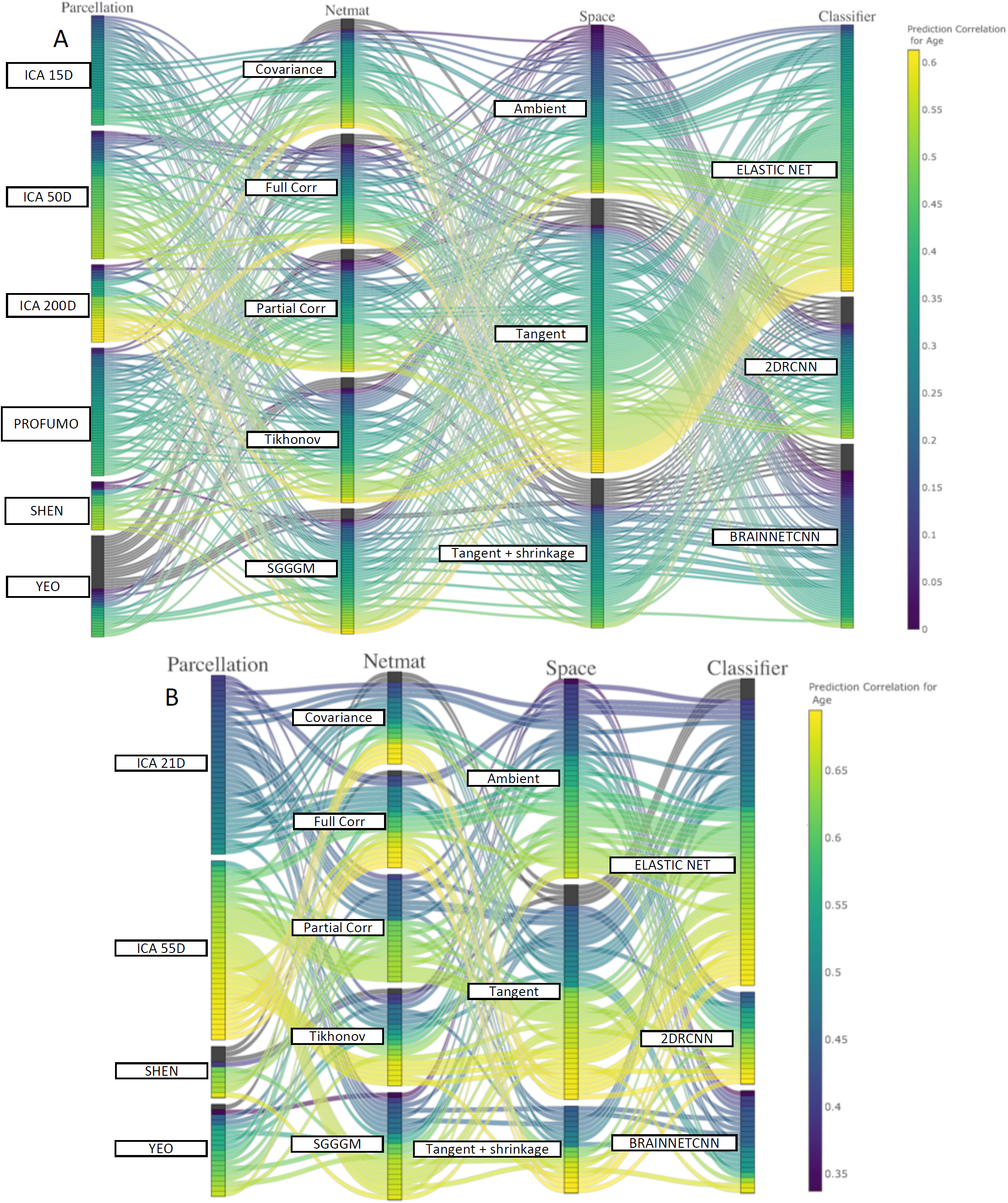
This parallel coordinates plot provides a visualization of all possible combinations of options in the pipeline to pipeline to predict age from functional connectivity **without deconfounding. [A] (HCP Data), [B] (UKB Data):** The lines are color-coded according to their prediction performance.

**Figure A.27:**
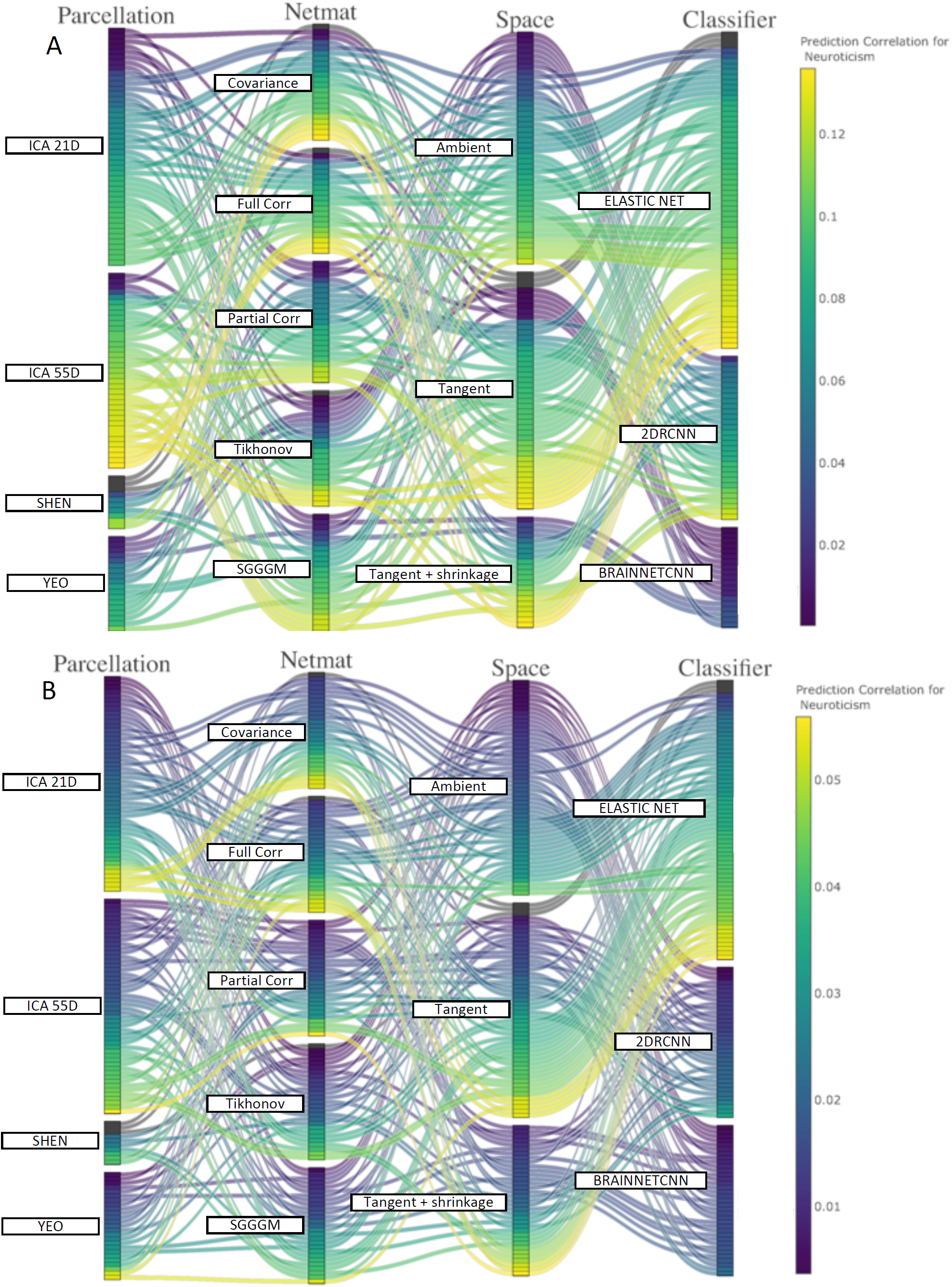
This parallel coordinates plot provides a visualization of all possible combinations of options in the pipeline to predict neuroticism score from functional connectivity. **[A]** shows the result before confounds removal and **[B]** shows the result after regressing out the confounds.

**Figure A.28:**
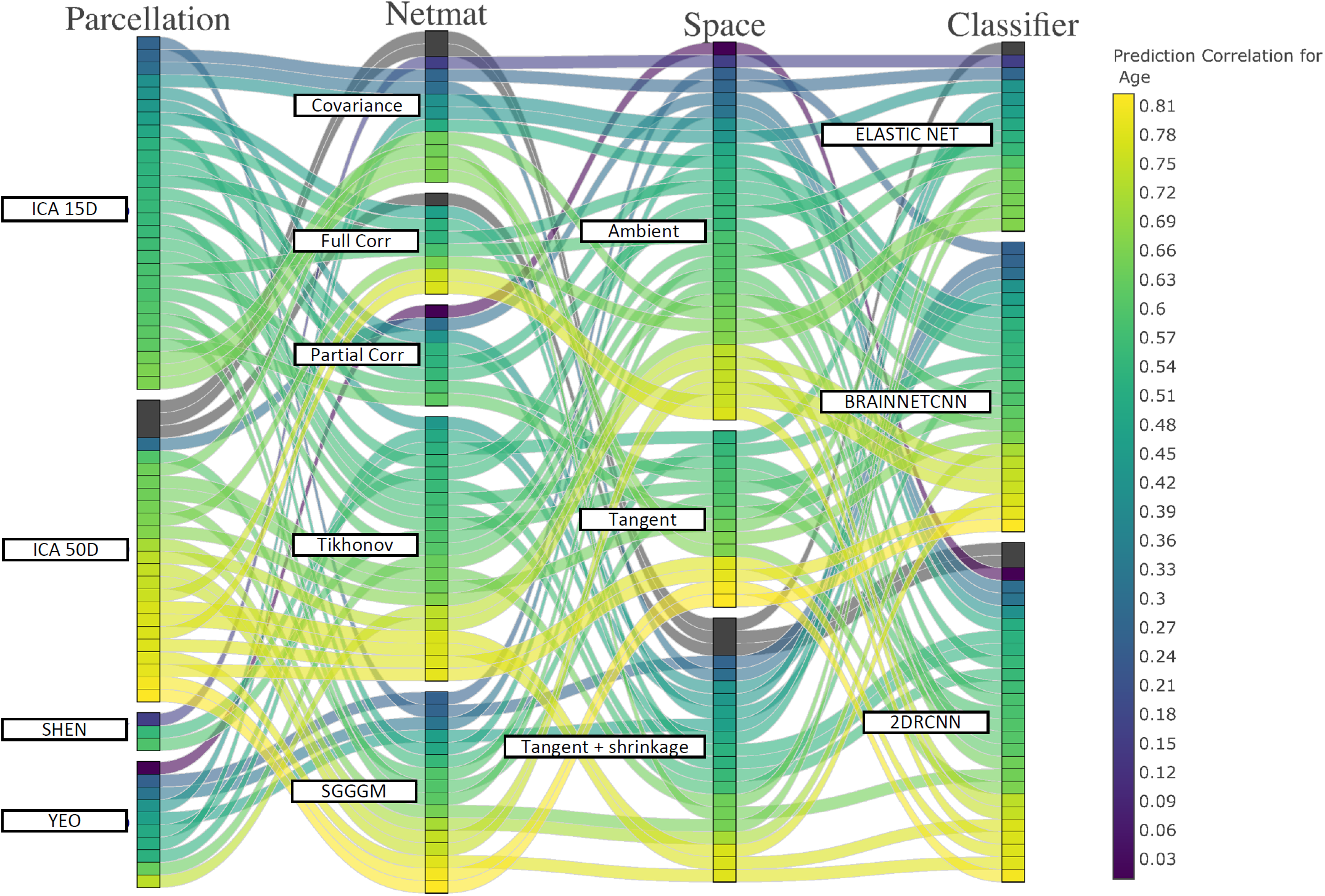
**(ABIDE)** This parallel coordinates plot provides a visualization of all possible combinations of options in the pipeline to predict age score from functional connectivity. The lines are color-coded according to their prediction performance.

**Figure A.29:**
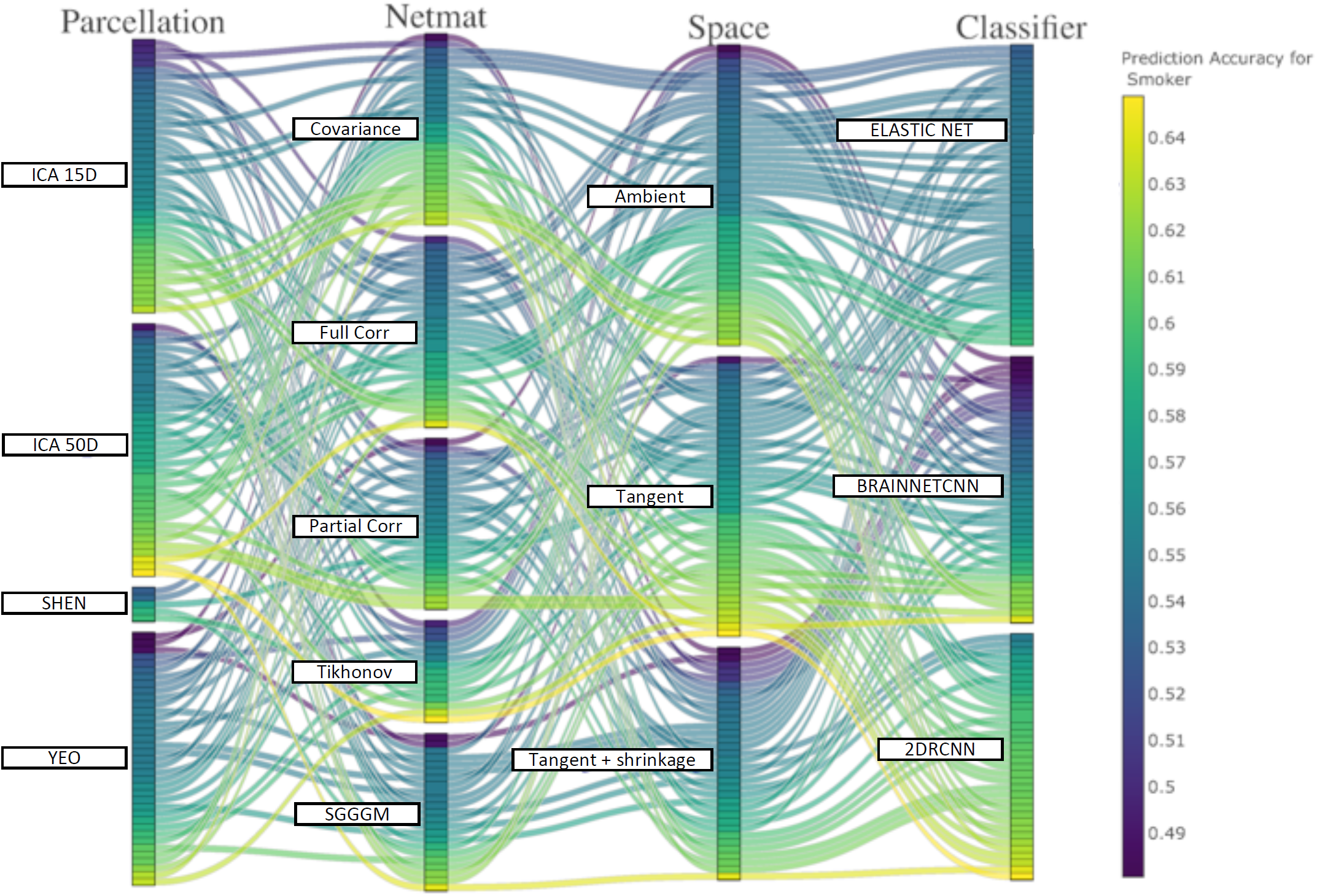
**(ACPI)** This parallel coordinates plot provides a visualization of all possible combinations of options in the pipeline to predict smoking status from functional connectivity. The lines are color-coded according to their prediction performance.

**Figure A.30:**
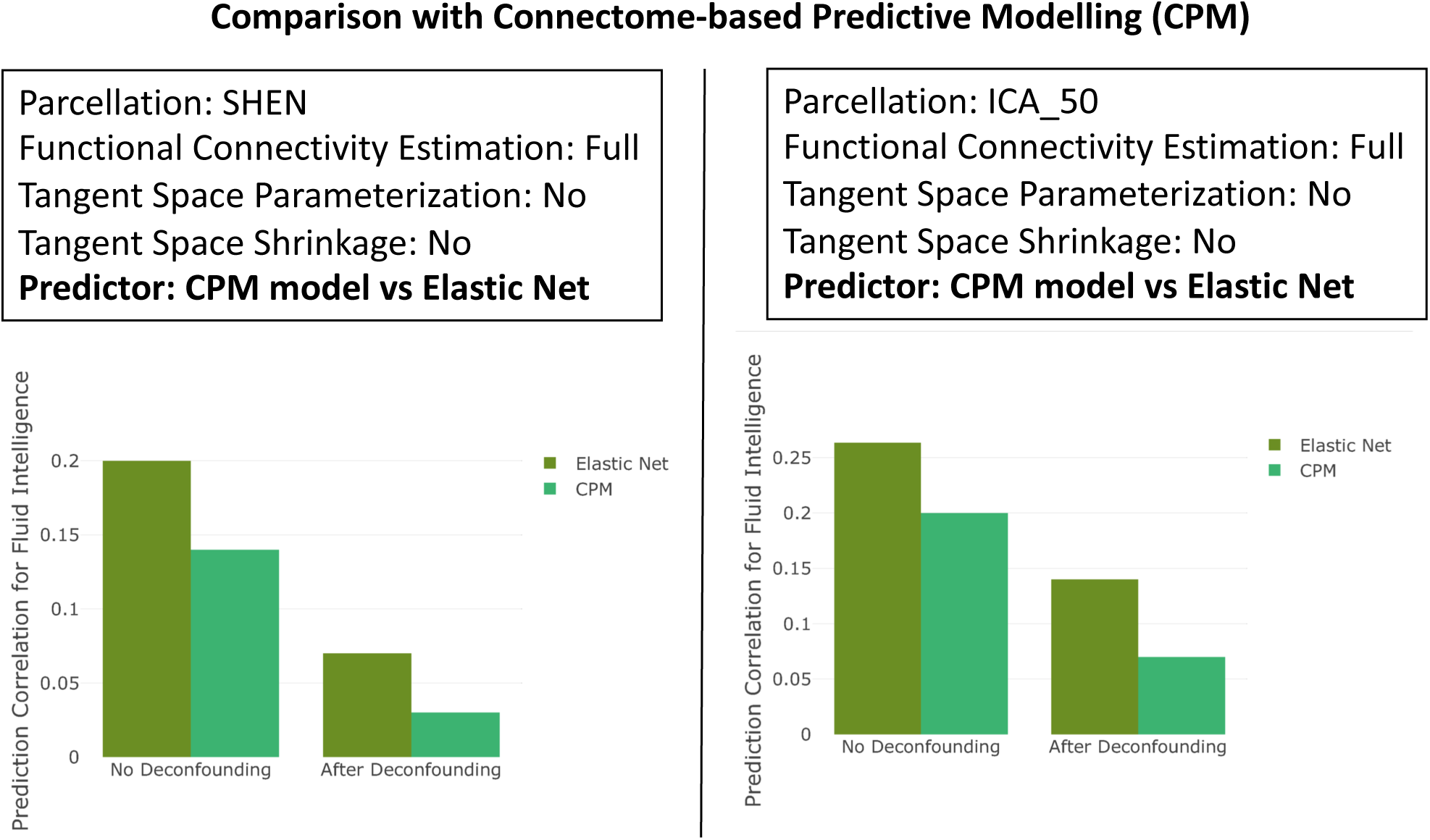
The comparison of Elastic Net and CPM prediction performance. CPM predictor/classifier is based on a model that averages connectivity edges from a subset of all edges.

**Figure A.31:**
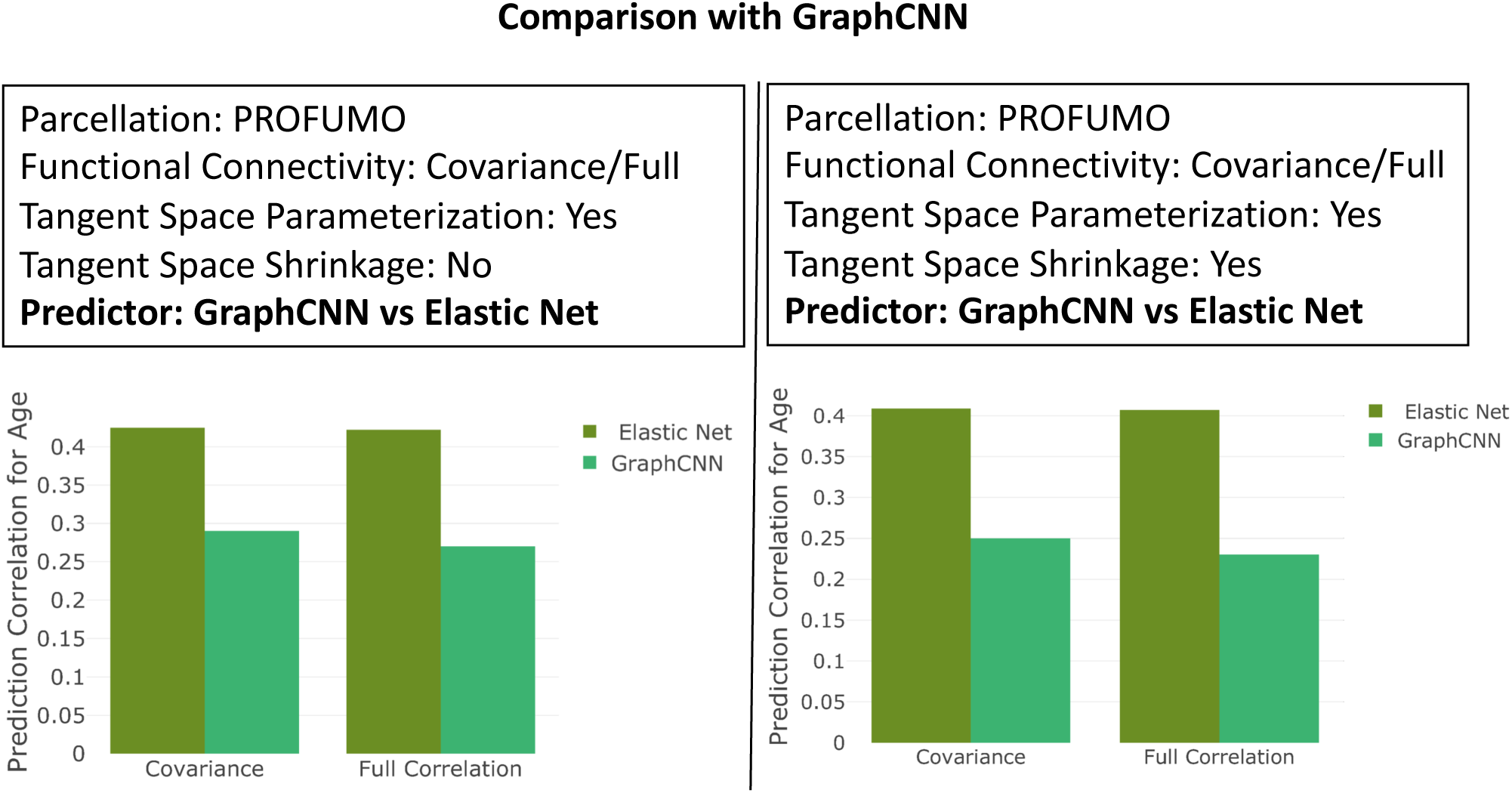
**(HCP)** The comparison of Elastic Net and GraphCNN prediction performance. We have chosen a subset (e.g., parcellation (PROFUMO), functional connectivity estimation (Full/Covariance)) from all available configurations to illustrate the performance of GraphCNN.

**Figure A.32:**
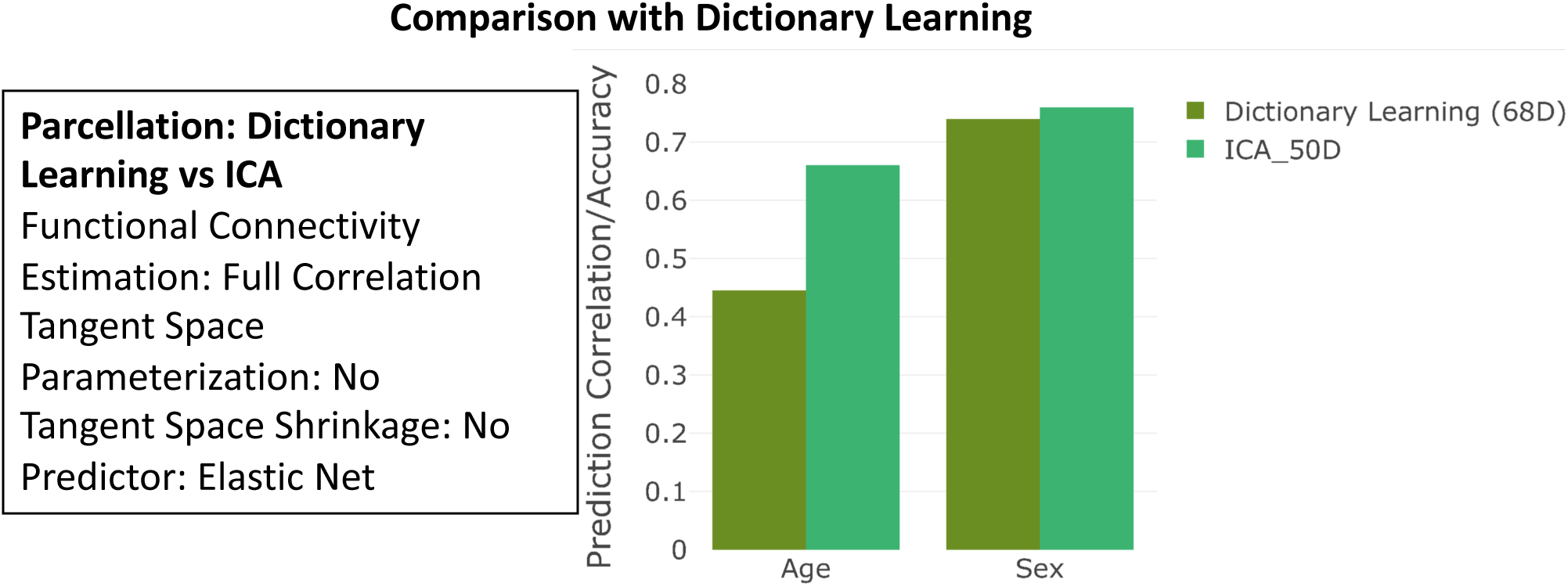
**(ABIDE)** The comparison of ICA and Dictionary Learning prediction performance. We have chosen a subset (e.g., parcellation (ICA and Dictionary Learning), functional connectivity estimation (Full) from all available configurations to illustrate the comparison.

**Figure A.33:**
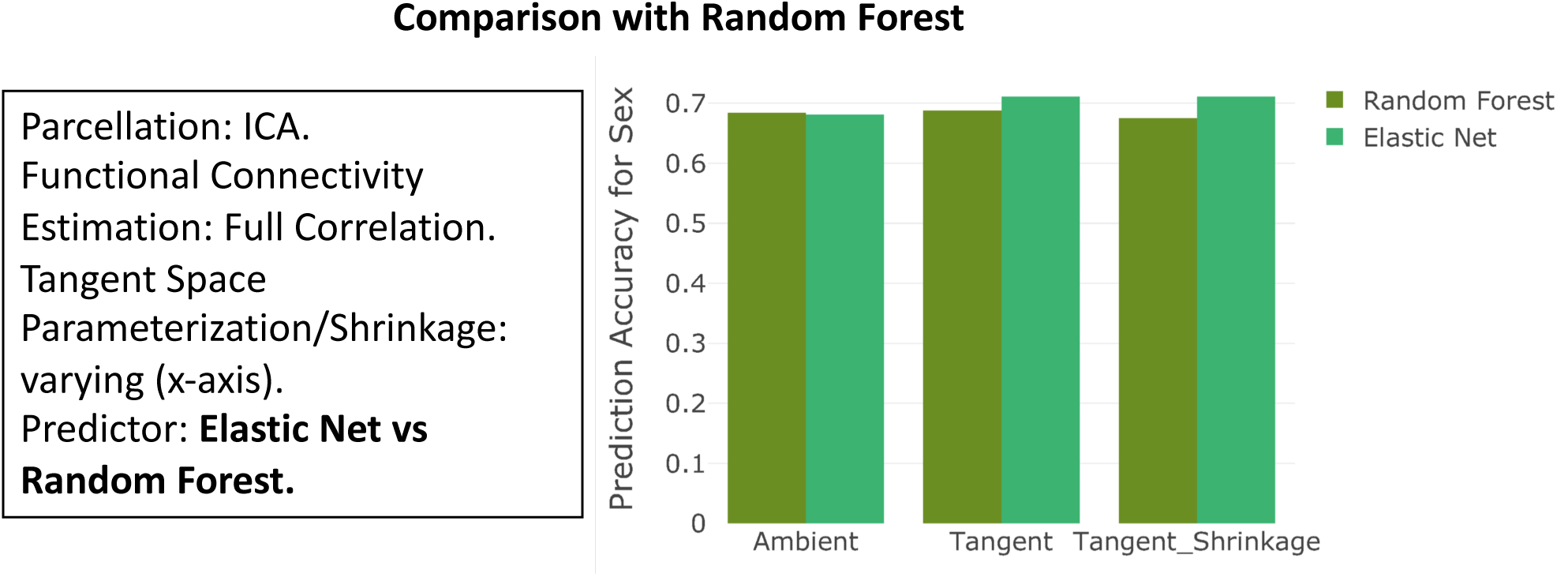
**(HCP)** The comparison of Elastic Net and Random forest prediction performance. We have chosen a subset (e.g., parcellation (ICA), functional connectivity estimation (Full)) from all available configurations to illustrate the comparison.

**Figure A.34:**
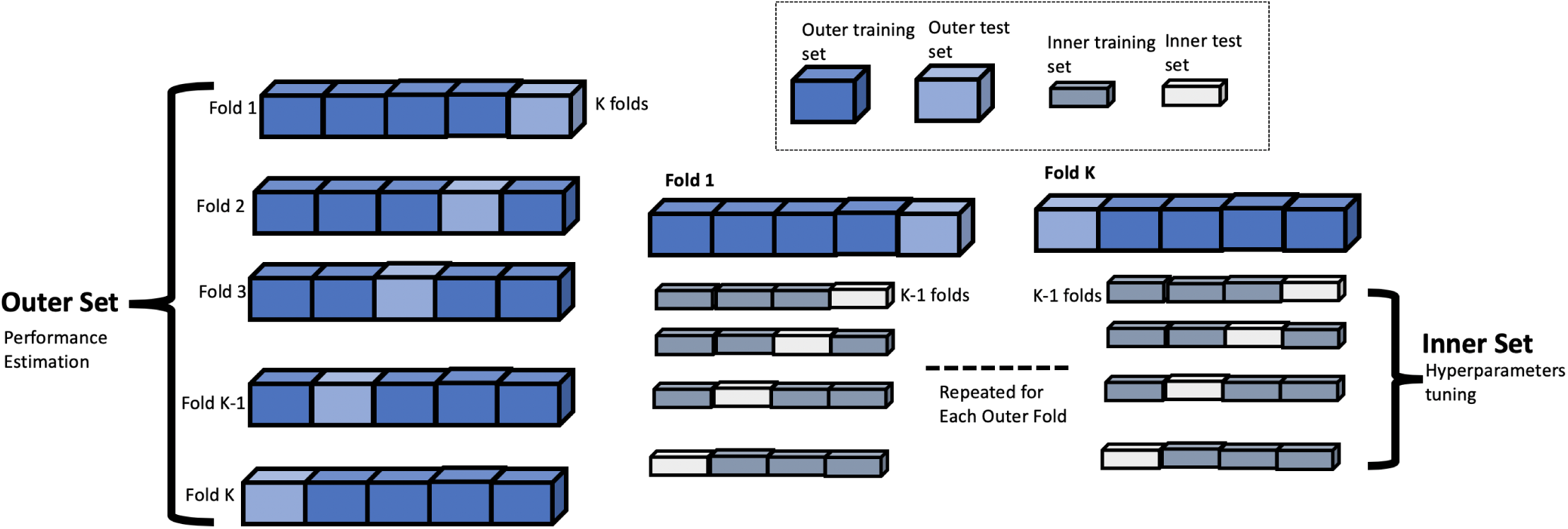
A summary figure explaining the process of nested cross-validation.

## Footnotes

2 To apply sICA, we first generated the group-PCA (Principal Component Analysis) output using MIGP (MELODICs Incremental group-PCA) [44] using data from all subjects. This comprises the top few thousands (1000-5000) weighted spatial eigenvectors from a group-averaged PCA, which is a very close approximation to the original data (fully concatenating all subjects’ time-series). Lastly, group sICA is run on the output of group-PCA.

3 Regarding the selection of artefactual components; this was part of the central processing done previously on behalf of UKB [36]. The decisions were made on the basis of spatial layout of components, and average temporal power spectrum.

4 For UKB, the ICA dimensionality cannot go up further (e.g., d = 200), because for larger dimensionalities, the additional components appear as artefactual; the number of non-artefact components barely grows at all (more than the existing 55). We believe this is because of the limitations of volumetric analysis possibly combined with the 6-minute limited duration of the acquisition.

5 We have N subjects, and for each subject, we have d parcels. In the case of optimisation of the regularising parameters, we first calculated the difference between the regularised subject precision matrices (*d* × *d* × *N*) and the group average of the unregularised subject precision matrices (*d* × *d*). Then, we calculated the sum of the resulting matrices (after squaring matrix elements), and obtained a *d* × *d* resulting matrix. Then, we summed up the elements in the upper triangular of the resulting matrix, and lastly computed the square root.

6 In a broad sense, we have used the word covariance for different variants of the functional connectivity estimates. This should not be confused with empirical covariance, 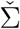.

7 *Predictor* is used sometimes as synonymous for input variable, and *classifier* is mostly used for categorical variables prediction. We have used *predictor/classifier* combination in an effort to make sure that the audience is not confused, and this implies prediction of both categorical and continuous non-imaging variables.

8 http://fcon_1000.projects.nitrc.org/indi/ACPI/html/index.html

9 The description of Berkson’s paradox is: “If there exist a causal structure *X* →*C* ←*Y*, where X and Y are not directly connected. If we try to account for (condition on) C in a partial correlation analysis (i.e., testing for conditional independence), by regressing C out of X and Y, we induce a negative correlation between X and Y” [79]. In the context of the discussion above, if there is no chance of confounds (C), being fed into a target variable (Y), but still by conditioning on C, we might create a spurious association between functional connectivity (X) and Y.

